# Isolating the role of corticosterone in the hypothalamic-pituitary-gonadal genomic stress response

**DOI:** 10.1101/2020.10.08.330209

**Authors:** Suzanne H. Austin, Rayna Harris, April M. Booth, Andrew S. Lang, Victoria S. Farrar, Jesse S. Krause, Tyler A. Hallman, Matthew MacManes, Rebecca M. Calisi

**Affiliations:** Department of Neurobiology, Physiology, and Behavior, University of California Davis, 1 Shields Avenue, Davis, CA, USA, 95616; Department of Molecular, Cellular and Biomedical Sciences, The University of New Hampshire, 434 Gregg Hall, Durham, NH, USA, 03824; Department of Biology, University of Nevada, Reno, 1664 N. Virginia Street, Reno, NV, USA, 89557-0314; Department of Fisheries and Wildlife, Oregon State University, 104 Nash Hall, Corvallis, OR 97331

**Author notes:** Department of Integrative Biology, Oregon State University, 3029 Cordley Hall, Corvallis, OR, USA, 97331; Department of Fisheries and Wildlife, Oregon State University, 104 Nash Hall, Corvallis, OR 97331.

## Abstract

The negative impacts of stress on reproduction have long been studied. A large focus of investigation has centered around the effects of the adrenal steroid hormone corticosterone (CORT) on a system of tissues vital for reproduction, the hypothalamus of the brain, the pituitary gland, and the gonads (the HPG axis). Investigations of the role of CORT on the HPG axis have predominated the stress and reproductive biology literature, potentially overshadowing other influential mediators. To gain a more complete understanding of how elevated CORT, characteristic of the stress response, affects the activity of the HPG axis, we experimentally examined its role at the level of the genome in both male and female rock doves (*Columba livia*). We exogenously administrated CORT to mimic circulating levels during the stress response, specifically 30 min of restraint stress, an experimental paradigm known to increase circulating corticosterone in vertebrates. We examined all changes in genomic transcription within the HPG axis as compared to both restraint-stressed birds and vehicle-injected controls, as well as between the sexes. We report causal and sex-specific effects of CORT on the HPG stress response at the level of the transcriptome. Restraint stress caused 1567 genes to uniquely differentially express while elevated circulating CORT was responsible for the differential expression of 304 genes. Only 108 genes in females and 8 in males differentially expressed in subjects who underwent restraint stress and those who were given exogenous CORT. In response to CORT elevation characteristic of the stress response, both sexes shared the differential expression of 5 genes, *KCNJ5, CISH, PTGER3, CEBPD*, and *ZBTB16*, all located in the pituitary. The known functions of these genes suggest potential influence of elevated CORT on immune function and prolactin synthesis. Gene expression unique to each sex indicated that elevated CORT affected more gene transcription in females than males (78 genes versus 3 genes, respectively). To our knowledge, this is the first study to isolate the role of CORT in HPG genomic transcription during a stress response. These results provide novel targets for new lines of further investigation and therapy development. We present an extensive and openly accessible view of the role corticosterone in the HPG genomic stress response, offering novel gene targets to inspire new lines of investigation of stress-induced reproductive dysfunction. Because the HPG system is well-conserved across vertebrates, these data have the potential to inspire new therapeutic strategies for reproductive dysregulation in multiple vertebrate systems, including our own.

## INTRODUCTION

Unpredictable perturbations or perceived threats in the environment can activate various endocrine cascades to promote behavioral and physiological coping mechanisms associated with survival. The hypothalamic-pituitary-adrenal (HPA) axis plays a central role in mediating a portion of these coping mechanisms (reviewed in Romero and Wingfield, 2015). Parvocellular neurons in the hypothalamus of the brain release hormones such as arginine vasotocin (AVT; in birds and lizards) and corticotropin releasing hormone (CRH) into the portal vasculature, which transports them to the anterior pituitary gland. There, AVT and CRH activate corticotrope cells to induce the release of adrenocorticotropic hormone (ACTH). ACTH travels through the bloodstream to the adrenal glands and binds to melanocortin 2 receptors (MC2R), which stimulates the synthesis of glucocorticoids [including corticosterone (*hereafter*, CORT) in adult birds and cortisol in many mammals]. CORT travels via the bloodstream to bind to its receptors, either nuclear or membrane bound, including the low affinity glucocorticoid receptor (GR) and high affinity mineralocorticoid (MR) receptor. The binding of CORT to nuclear receptors affects gene transcription throughout the body due to their global distribution (Romero and Wingfield, 2015). Receptor activation modifies metabolism, immune function, and induces behavioral and physiological changes (Sapolsky et al., 2000; Wingfield et al., 1998; Wingfield and Kitaysky, 2002).

When individual survival is favored over other energetically costly activities (Wingfield et al., 1998), the chronic depletion of bodily resources can inhibit other important biological processes (e.g., reproductive function, Sapolsky et al., 2000). The activation and regulation of reproduction and associated behaviors is mediated by a biological system of tissues consisting of the hypothalamus, pituitary, and gonads, referred to as the “HPG” axis. Its most well-studied endocrine cascade involves production and secretion of gonadotropin-releasing hormone (GnRH) from the preoptic area of the hypothalamus, which causes pituitary secretion of gonadotropins, luteinizing hormone (LH) and follicle stimulating hormone (FSH). LH and FSH travel through the bloodstream and act upon receptors in the gonads, stimulating gametogenesis and the synthesis of sex steroids, such as testosterone and estradiol. These sex steroids then bind with receptors within the HPG axis to facilitate reproduction and sexual behaviors, and they also act as a negative feedback control mechanism.

It has long been known that exposure to certain stressors, and the subsequent physiological response, can negatively impact sexual behavior and reproduction. The activation of the HPA axis affects HPG function at multiple levels because MR and GR are globally distributed and are expressed throughout the body (Angelier and Chastel, 2009; Geraghty and Kaufer, 2015). Elevation of CORT, a significantly influential and well-studied hormone in the stress response, can suppress the release of the reproductive hormone, GnRH, at the level of the brain, by interacting with gonadotropin inhibitory hormone [GnIH; also referred to as RFamide-related peptide (*RFRP)*], or with kisspeptin neurons (in mammals) (Calisi, 2014). This, in turn, reduces the rate of depolarization of the GnRH-1 neuron and subsequent GnRH-1 release, which causes the reduction of LH, FSH, and steroidogenic capacity and gametogenesis. Thus, activation of the HPA axis in response to environmental perturbations has the ability to affect each endocrine-producing tissue of the HPG axis, and therefore, reproduction.

Research on the influential role of CORT on the HPG axis has predominated much of the stress and reproductive biology literature, which has potentially overshadowed other influential mediators of stress. In this study, we used the model of the rock dove (*Columba livia*) to experimentally test the extent to which changes in gene expression within the HPG axis were explained by an increase in circulating CORT – a key characteristic of a stress response. Previously, our group described the genomic transcriptome community of sexually mature, non-breeding male and female doves (MacManes et al., 2017). Then, we tested how gene transcription within the HPG axis of both sexes was affected by activation of the HPA axis, the latter which we stimulated using a common restraint stress paradigm in which subject mobility is restricted for 30 min (Calisi et al., 2018). We reported a heightened and mostly sexually specific genomic stress response throughout the HPG axis. In this study, we isolated the role of elevated CORT in both sexes using the same restraint-stress paradigm to determine its causal and sex-typical effects on HPG gene transcription. We did this by exogenously administrating CORT to mimic circulating levels following 30 min of restraint stress. We then compared differential gene expression within the HPG following 30 min of exogenous CORT and 30 min of restraint stress. We report stress-induced and sex-specific changes in HPG genomic activity caused by elevated circulating CORT. We predicted a reduction in differentially expressed genes in response to exogenous CORT compared to restraint stress.

## RESULTS

### Corticosterone

Both restraint stress and exogenous CORT treatment increased circulating levels of CORT in both sexes (treatment: *F*_3, 71_ = 83.4, *P* < 0.001, sex: *F*_1, 71_ = 0.4, *P* = 0.279; Fig. 1). Circulating CORT concentrations did not differ between the control group of our previous restraint stress study and our current study [restraint-stress control group vs. CORT control group: least-squares mean back-transformed estimate (± SE) = 1.31 ± 0.22, (Dunnett-adjusted) *P* = 0.631; CORT treatment vs. vehicle control=14.69 ± 0.23, *P* < 0.001; restraint-stress treatment vs. CORT treatment= 0.53 ± 0.20, *P* < 0.001]. Median circulating CORT concentrations were higher in the exogenous CORT treatment group as compared to our previous restraint-stress treatment group (47.05 ng/mL ± 13.34 SD vs. 29.8 ng/mL ± 16.26 SD), though there was substantial overlap of values (restraint stress range: 1.3 – 64 ng/mL; Fig. 1).

**Figure 1.**
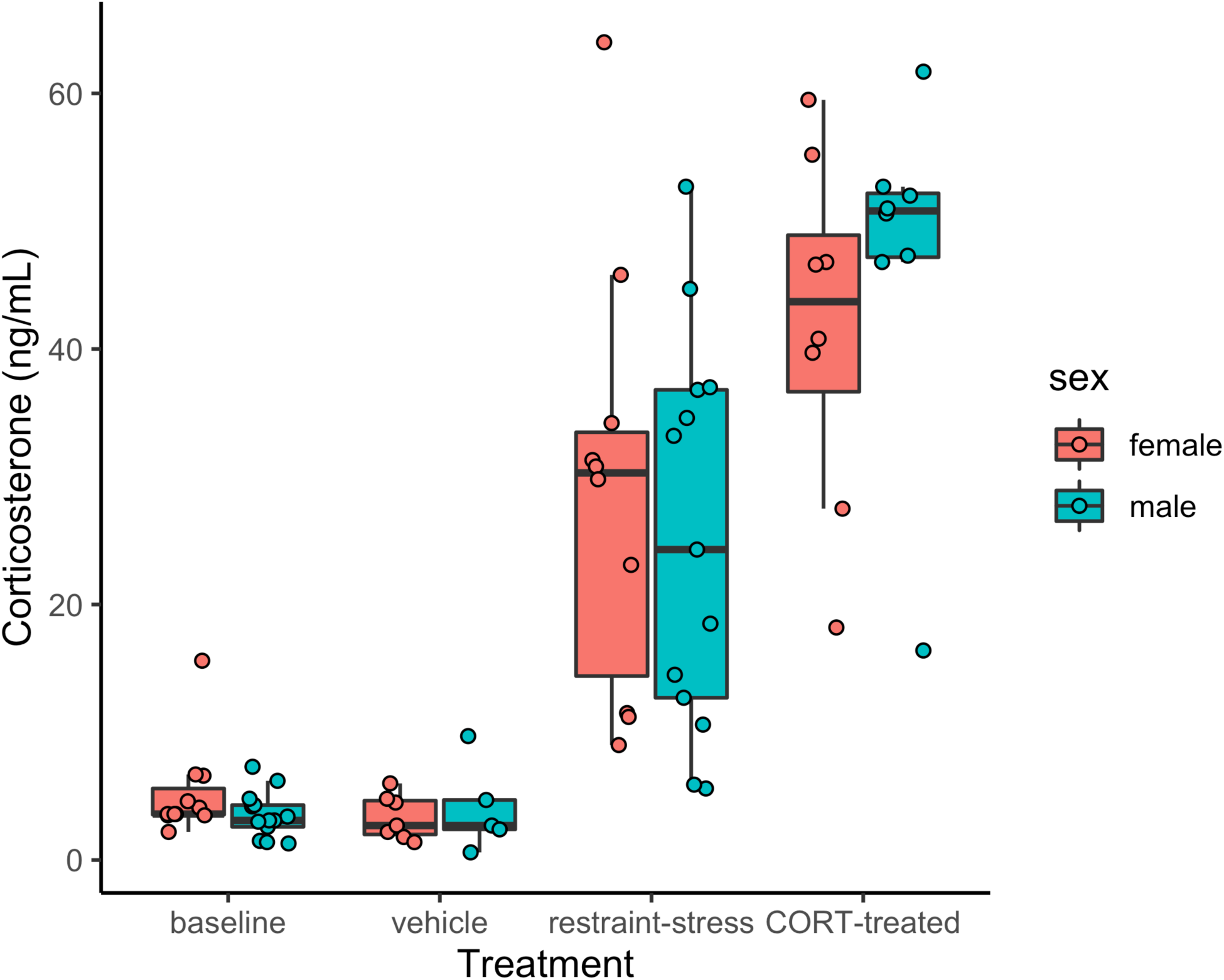
Circulating corticosterone. (ng/mL) of male (aqua) and female (pink) *C. livia* of restraint-stress (baseline control and restraint-stress, left panel) compared to the exogenous treatment (vehicle and CORT-treated, right panel).

### Sequence Read Data and Code Availability

In total, hypothalami, pituitary, and gonads (testes or ovaries) from 11 males and 16 females were sequenced (n=81 samples in total). Each sample was sequenced with between 6.6 million and 23.5 million read pairs. Read data are available using the European Nucleotide Archive project ID PRJEB28645. Code used for the analyses of these data are available at https://github.com/macmanes-lab/RockDove/tree/master/CortStudy.

### Transcriptome Assembly Characterization

The Rock Dove v. 1.1.1 transcriptome contains 97,938 transcripts, of which 5,133 were added as part of this study to the previous version 1.1.0 transcriptome. This newly compiled transcriptome data improves genic contiguity, increasing the number of complete BUSCOs 1.4% to achieve 87.5% relative to the v. 1.1.0 assembly.

### Sequence Read Mapping and Estimation of Gene Expression

Raw sequencing reads corresponding to individual samples of hypothalami, pituitary glands, and gonads were mapped to the *C. livia* reference HPG axis transcriptome (v. 1.1.1) using Salmon, resulting in 80% to 90% read mapping. These data were imported into R and summarized into gene-level counts using tximport, after which, edgeR was used to generate normalized estimates of gene expression. A total of 14,938 genes or their isoforms were expressed in HPG tissues.

### Evaluation of Genomic Expression

Global patterns of gene expression were analyzed using edgeR to assess the HPG transcriptomic response to experimentally elevated circulating CORT. After controlling for > 35,000 multiple comparisons, the count data were normalized using the TMM method (D. J. McCarthy et al., 2012; M. M. McCarthy et al., 2012), which, in brief, uses a set of scaling factors for library sizes to minimize inter-sample log-fold changes for most genes. This analysis revealed a significant transcriptomic response of the HPG axis to CORT treatment, with females experiencing more differential expression as compared to males (Fig. 2). All differentially expressed genes in female and male HPG tissue in response to our 30 min CORT treatment can be found at https://cortstudy.page.link/DEresults.

**Figure 2:**
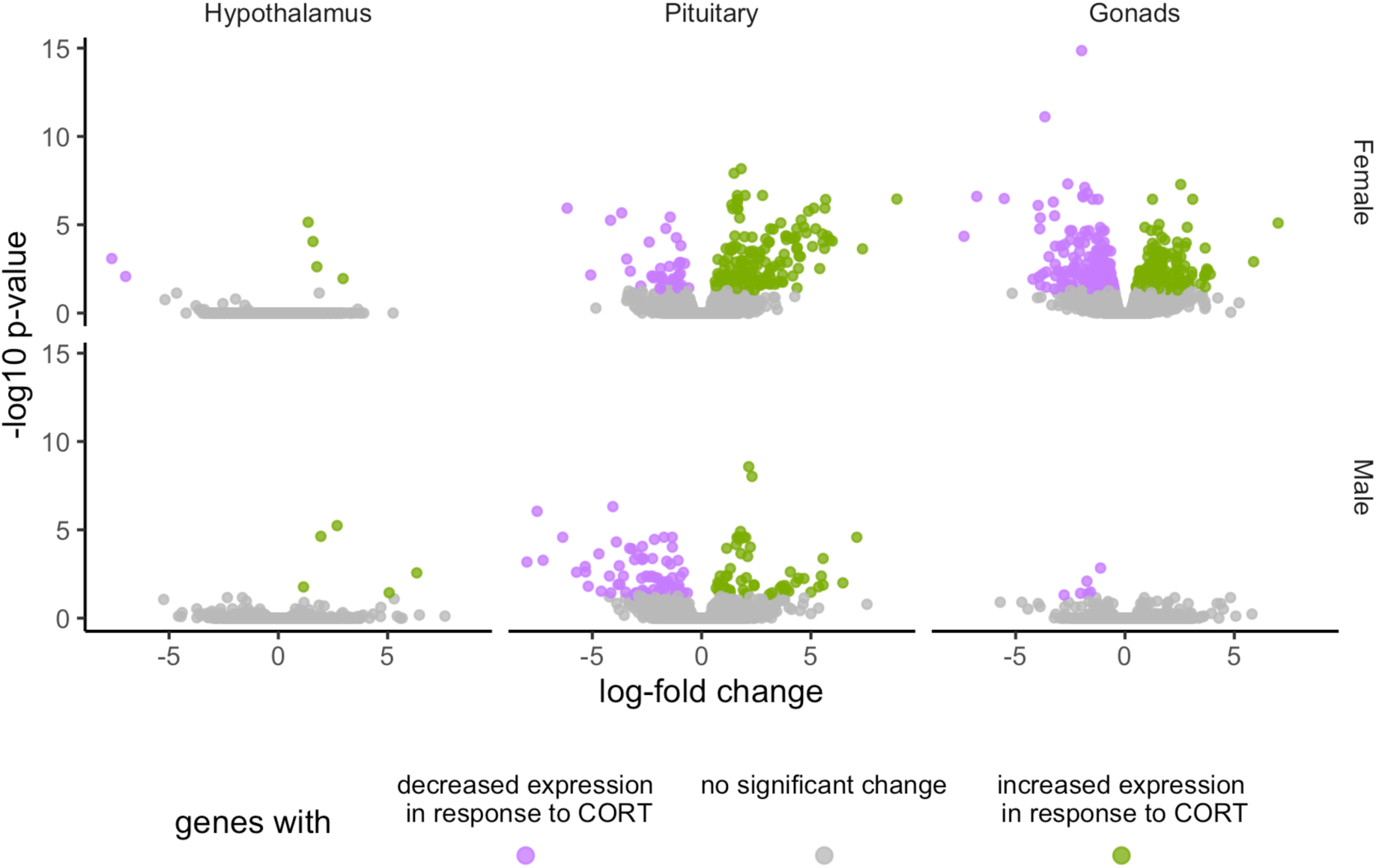
Volcano plots depicting total transcriptional changes of CORT-treatment group as compared to vehicle-control group. Volcano plots represent the number of differentially expressed genes throughout the HPG axis of females (top) and males (bottom). The x-axis represents log-fold change of differentially expressed genes and the y-axis -statistical significance (log10 p-values where FDR <0.01). Each datapoint represents a differentially expressed gene in the HPG axis. Significant differences in gene expression are represented in green (increased expression) or purple (decreased), or gray (no significant difference) in response to corticosterone treatment.

### Unique Differential Gene Expression in Response to Exogenous CORT treatment

We found 304 genes responsive to our exogenous CORT treatment that were not responsive to our restraint stress treatment. Of these genes, 6 exhibited differential expression patterns in the hypothalamus (females: 5, males: 3), 179 in the pituitary (females 136, males 71), and 209 in the gonads (females 208, males 2) (Fig 3; Supplemental Information 1: Tables 1-6). Across all tissues, the differential expression of 38 genes was shared between the sexes in response to exogenous CORT treatment (hypothalamus: 2, pituitary: 28, and gonads: 1; Supplemental Information 1: Table 7-9).

**Table 1.** Differential gene expression (DGE) shared by both sexes during the stress response due to elevated circulating CORT concentrations. Log fold change (logFC), false discovery rate (FDR) and gene function, in brief, are reported. All identified genes increased in expression in response to treatment (indicated by a negative logFC). All DGE occurred in the pituitary

| Gene | Gene Name | Reported Function | CORT-treated |  |  |  | Restraint stress |  |  |  |
| --- | --- | --- | --- | --- | --- | --- | --- | --- | --- | --- |
|  |  |  | Female |  | Male |  | Female |  | Male |  |
|  |  |  | logFC | FDR | logFC | FDR | logFC | FDR | logFC | FDR |
| *KCNJ5* | Potassium voltage-gated channel subfamily J member 5 | *KCNJ5*, also known as *GIRK4*, is a G protein regulated inward-rectifying potassium channel upregulates in response to CORT and restraint stress. *KCNJ5* is under the regulation of CORT signaling via a glucocorticoid response element (GRE) on its promoter region (Muma and Beck, 1999). GIRK channels can influence the resting membrane potential of cells by affecting the flow of potassium into the cell. Cells where *KCNJ5* is upregulated are more difficult to activate. Site-specific information is required to understand specific actions of corticosterone on GIRK signaling (Muma and Beck, 1999). In the hippocampus, neuronal exposure to corticosterone results in selective activation of ligand gated receptors through GIRK1 and 2 pathways (Muma and Beck, 1999). The GIRK pathway can also act as a cellular effector of dopaminergic action on lactotrophs in the anterior pituitary (Christensen et al., 2011; Gregerson et al., 2001). Based on this action, it is feasible that upregulated *KCNJ5* may decrease prolactin, via its action on lactotrophs, following acute exposure to CORT or a stressor. Prolactin is a hormone critical for supporting parental care and immune function in birds (Austin and Word, 2018). Our work did not find that *PRL* was differentially expressed in the pituitary of either treatment group though it upregulates in the male hypothalamus of CORT-treated birds. *KCNJ5* can also influence biochemical processes associated with aldosterone production thereby influencing electrolyte balance (Bollag, 2014). Due to the influence of *KCNJ5* and GIRK pathways in the hypothalamus and prolactin in the pituitary, this gene could be a potential means by which stress affects reproduction. | -1.7 | 4.09E-06 | -1.9 | 2.70E-05 | -1.6 | 4.09E-03 | -1.5 | 7.27E-03 |
| *CISH* | Cytokine-inducible SH2-containing protein | *CISH* is another shared gene that upregulates in response to both restraint stress and CORT treatments. *CISH* is an immediate early gene that is involved in negative feedback of the Jak2-Stat5 pathway (Dif et al., 2001; Matsumoto et al., 1997). The CISH protein competitively binds to phosphorylated prolactin receptors (PRLr), which attenuates Stat5 binding and subsequent downstream pathway signaling (Dif et al., 2001; Radhakrishnan et al., 2012). This action is one way that CISH may inhibit prolactin transcription and downstream events of the Jak-Stat pathway. Evidence also suggests that cytokines are important cell-signaling molecules that play a role in the inflammatory response (Yasukawa et al., 2000). Thus, if increased CISH activity is indicative of decreased cytokine activity or attenuation of PRLr binding, this could be a possible mechanism by which stress could influence both immune function and parental care. | -1.6 | 3.72E-07 | -1.7 | 2.63E-05 | -1.6 | 1.91E-10 | -1.8 | 3.16E-10 |
| *PTGER3* | Prostaglandin E receptor 3 | *PTGER3* increased in expression in response to both restraint stress and CORT-treatments. *PTGER3* encodes a G-protein-coupled receptor, EP3, that binds to its ligand, PGE2. The binding of EP3 to PGE2 results in the activation of an inhibitory G protein that reduces adenylyl cyclase activity and causes a decline in cellular cAMP. Prostaglandins have been linked to inflammation-induced activation of the HPA axis and act as an important immunoregulator (Furuyashiki and Narumiya, 2011; Oka, 2004; Rivest, 2001). PGE<sub>2</sub> binds to EP1 and EP3, which promote an ACTH response (Furuyashiki and Narumiya, 2011). Recall that ACTH is synthesized in the pituitary and travels through the bloodstream where it binds to melanocortin 2 receptors in the adrenals and promotes CORT synthesis. Cytokines may also induce prostaglandin synthesis (PGE<sub>2</sub>) and EP3 activation of the HPA axis (Furuyashiki and Narumiya, 2011). EP3's importance in inflammation-induced HPA activation was shown by a study that used addressed how EP3 knockout mice responded to an induced fever (by injecting pyrogen; Oka, 2004). EP3<sup>-/-</sup> mice did not develop a fever following treatment with pyrogen whereas controls did (Oka, 2004). This result suggests that EP3 activates the HPA axis in response to illness (Oka, 2004). Activation of the HPA via EP3 also includes vasodilation of the skin and increased metabolism of brown fat (Furuyashiki and Narumiya, 2011). *PTGER3* may be another means by which the HPA axis is activated, J7 and critically, *PTGER3* appears to be a means that the HPA axis primes itself for potential injury or infection following exposure to a stressor. | -2 | 2.19E-07 | -2 | 2.70E-05 | -1.5 | 7.50E-03 | -1.8 | 7.91E-04 |
| *CEBPD* | CCAAT enhancer binding protein delta | *CEBPD* upregulates in response to CORT and restraint stress. This gene encodes a CCAAT/enhancer-binding protein that acts as transcription factor and responds to glucocorticoids (Ko et al., 2015). These proteins act in metabolism, adipogenesis, immune and inflammatory responses, and PRL regulation (Ko et al., 2015; Tong et al., 2011). Tong et al. (Tong et al., 2011) suggest that PRL expression and lactotroph cell proliferation in the pituitary are linked: *CEBPD* plays a key role in coordinating their actions. Forced expression of *CEBPD* suppressed PRL expression in PRL-secreting cells and lactotroph proliferation in rats by binding to the PRL promoter (Tong et al., 2011). *CEBPD* may also be activated by a number of inflammatory factors such as PGE2 (Ko et al., 2015). Transcription of *CEBPD* following an inflammatory stimuli can be activated by a number of signaling pathways including JAK (Ko et al., 2015). Thus, upregulation of this gene may increase inflammatory responsiveness and suppress prolactin synthesis in response to a stressor or environmental perturbation. This gene is another potential target for understanding the role of CORT on the inflammatory response and suppression of reproduction. | -1.6 | 1.25E-06 | -1.7 | 3.50E-05 | -1.9 | 1.73E-05 | -2.6 | 3.10E-09 |
| *ZBTB16* | Zinc finger and BTB domain containing 16 | *ZBTB16* upregulates in the pituitary of males and females exposed to restraint stress and CORT. It is a transcription factor that generally acts as a transcriptional repressor, downregulating gene transcription through recruitment of other transcriptional co-repressors and histone deacetylases that modify chromatin (Costoya, 2007; McConnell and Licht, 2007). This protein has been implicated in reducing cell proliferation and increasing cell death via apoptosis in various cancer cell lines and during development (Costoya, 2007). Glucocorticoids have been shown to upregulate *ZBTB16* in the brain of mice and cancer cell lines (Peppi et al., 2011; Wasim et al., 2010). While no known study has examined *ZBTB16* in the pituitary, it may possibly play a role in the negative feedback of CORT on pituitary gene expression and cell proliferation. For example, *ZBTB16* may repress transcription and recruit epigenetic mechanisms, to downregulate genes associated with HPA axis activity. Further, *ZBTB16* may inhibit cell growth and proliferation of corticotrophs in the pituitary. Glucocorticoids have been shown to suppress growth of corticotrophic tumors (Losa et al., 2000) and have | -1.7 | 1.25E-06 | -2.3 | 9.47E-09 | -2.9 | 5.57E-08 | -2.2 | 2.95E-05 |
|  | generally inhibitory effects on DNA synthesis in the pituitary (McNicol and Carbajo-Perez, 1999), possibly through a mechanism including *ZBTB16*. Taken together, these results suggest *ZBTB16* may play a role in negative feedback of the HPA axis by affecting corticotroph transcription and cell proliferation. |  |  |  |  |  |  |  |  |  |

**Figure 3:**
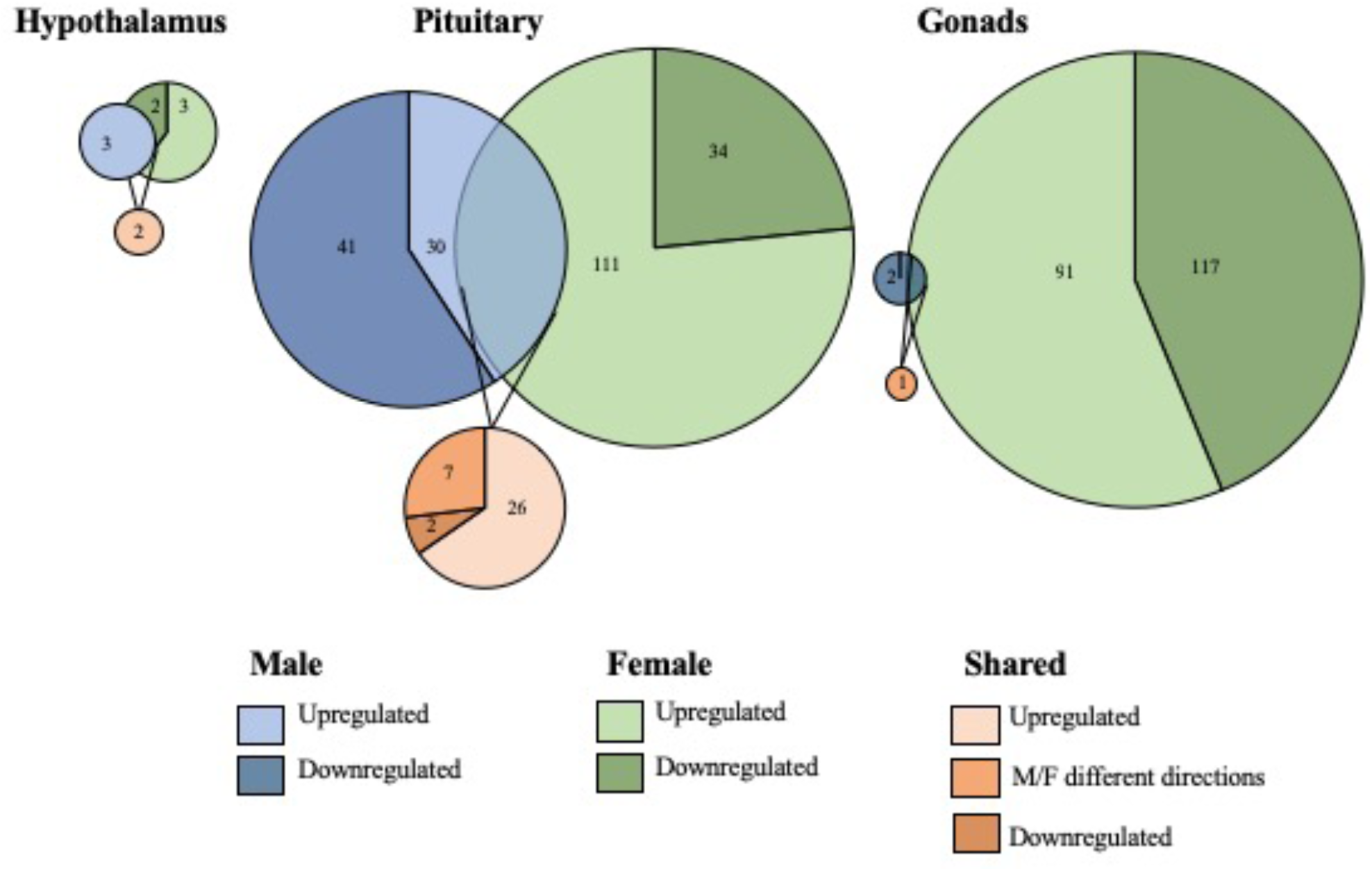
Sex-biased differential genomic expression in response to exogenous CORT treatment. A weighted Venn diagram depicting the overlap of the number of differentially expressed genes between the sexes in the hypothalamus, pituitary, and gonads in response to CORT as compared to vehicle. Genes that upregulated in expression in response to CORT are depicted by a lighter shade; genes that down-regulated in response are depicted by a dark shade. Numbers within shaded areas indicate the number of CORT-responsive genes.

### Unique Differential Gene Expression in Response to Restraint Stress

We found that 1567 genes differentially expressed uniquely in response to restraint stress as compared to those treated with exogenous CORT. Of these, 147 genes were differentially expressed in the hypothalamus (females: 130, males: 17), 562 in the pituitary (females 484, males 78), and 966 in the gonads (females 960, males 6) (Fig. 4). Across all tissues, the differential expression of 38 genes was shared between the sexes in response to restraint stress treatment (hypothalamus: 1, pituitary: 35, gonads: 1). The tables of restraint stress specific genes can be found in Tables 1-9 of the Supplemental Information 2.

**Figure 4:**
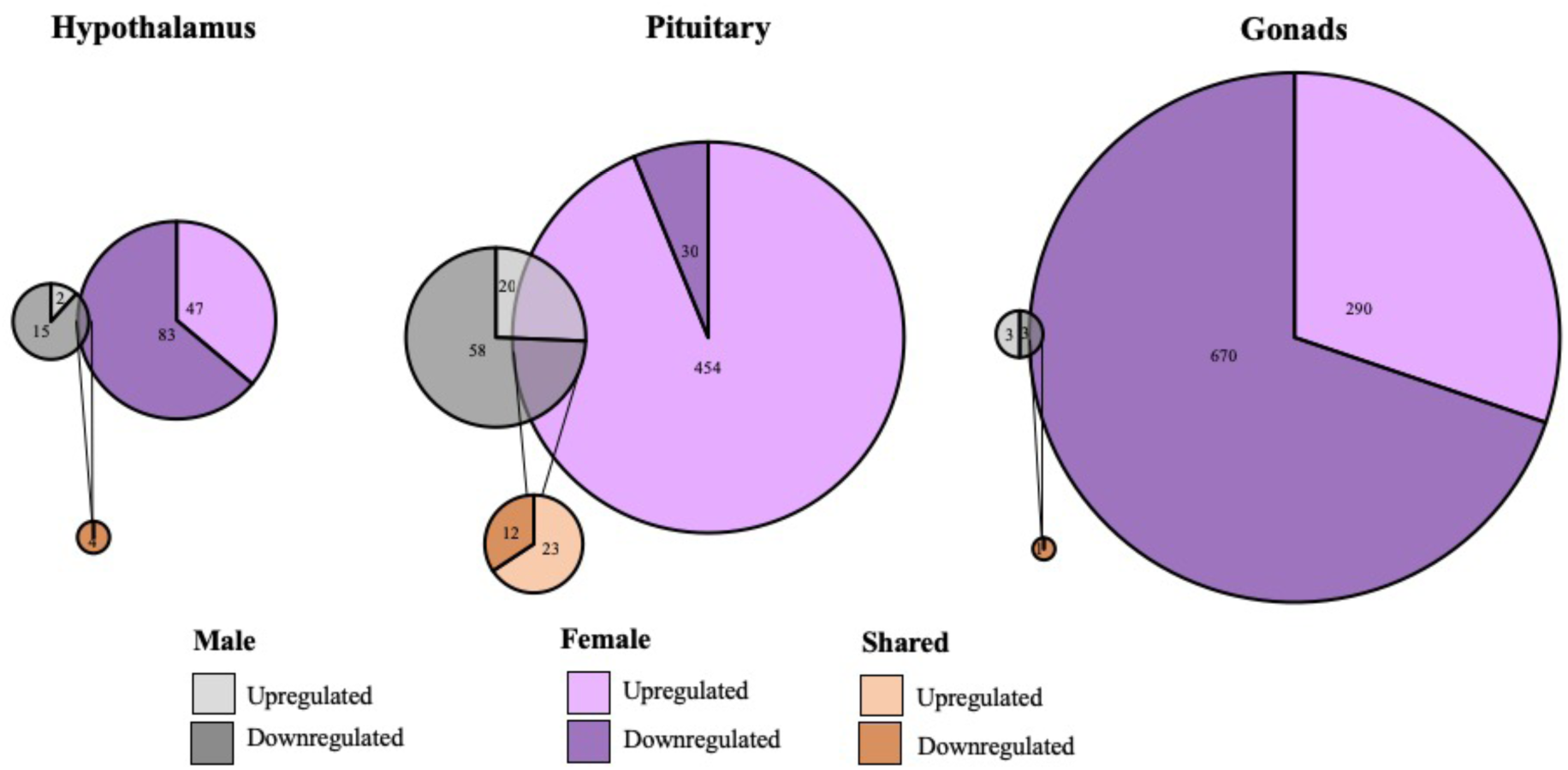
Sex-biased differential genomic expression that uniquely responded to restraint-stress treatment but not CORT treatment. A weighted Venn diagram depicting the overlap of the number of differentially expressed genes between the sexes in the hypothalamus, pituitary, and gonads that uniquely responded to restraint stress and not CORT treatment as compared to controls. Genes that upregulated in expression in response to CORT are depicted by a lighter shade; genes that down-regulated in response are depicted by a dark shade. Numbers within shaded areas indicate the number of restraint stress-responsive genes.

### Isolation of Restraint Stress-Responsive Gene Activity Attributed to Circulating CORT Elevation

#### Changes in Expression Shared by Both Sexes

To isolate the effects of CORT on the HPG transcriptomic stress response observed after 30 min of restraint stress, we identified shared differential expression resulting from both exogenously administered CORT and restraint stress as compared to their respective controls. In both the male and female pituitary, 5 genes differentially expressed in exogenous CORT and restraint-stress treated birds: *KCNJ5, CISH, PTGER3, CEBPD*, and *ZBTB16*. We report the results from our differential analysis [log-fold change (logFC) and false discovery rate (FDR)] with a brief description of known gene functionality in vertebrates in Table 1. There were no shared genes between males and females at the level of the hypothalamus or gonads that responded to both exogenous CORT and restraint-stress treatments.

#### Changes in Expression Unique to Each Sex

Within the HPG axis, we identified 108 genes unique to females (including 5 isoforms) and 8 genes unique to males whose response to 30 min of restraint stress was attributed to circulating CORT elevation. In females, 78 of these genes were isolated in the pituitary (Table 3), and 32 genes in the ovaries (Table 4). *CEPBD* and *TSC22D3* differentially expressed in both the pituitary and ovaries and are therefore included in both tissue gene counts. In males, 8 genes were isolated in pituitary. Only 3 of these genes uniquely differentially expressed in males, *KLF9, SLAINL*, and *PVALB* (Table 2). We did not identify any restraint stress-induced gene activity explained by exogenous CORT elevation within the hypothalamus of either sex or in the testes. A brief summary of gene function for these differentially expressed genes is also provided in tables 2-4.

**Table 2:** Differential gene expression (DGE) in the male pituitary during the stress response due to elevated circulating CORT concentrations. Red indicates upregulation while blue indicates downregulation.

| Entrez ID | Gene | Gene Name | Function | CORT |  | Restraint stress |  |
| --- | --- | --- | --- | --- | --- | --- | --- |
|  |  |  |  | logFC | FDR | logFC | FDR |
| 107052707 | *CEBPD* | CCAAT enhancer binding protein delta | Described in Table 1 | -1.73 | 3.50E-05 | -2.57 | 3.10E-09 |
| 395335 | *CISH* | cytokine inducible SH2 containing protein | Described in Table 1 | -1.70 | 2.63E-05 | -1.76 | 3.16E-10 |
| 395925 | *KCNJ5* | potassium voltage-gated channel subfamily J member 5 | Described in Table 1 | -1.92 | 2.70E-05 | -1.48 | 7.27E-03 |
| 770238 | *KLF9* | Kruppel like factor 9 | Krüppel-like factors are a group of highly conserved genes in vertebrates that act as immediate early genes or hormone mediating transcription factors. They can be direct transcription factors, cofactors/accessory transcription factors, or can mediate nuclear receptor genes to regulate hormone function (Knoedler and Denver, 2014). *KLF9* responds independently or synergistically to thyroid hormones and glucocorticoids in the brains of mice and frogs while in the uterus and ovaries it responds to progesterone and estradiol (Knoedler and Denver, 2014). CORT acts on *KLF9* via a GRE in its promoter region (Knoedler and Denver, 2014). This gene influences the structure and differentiation of neurons and axons (Bonett et al., 2009; Knoedler and Denver, 2014). *KLF9* influenced dendritic spine growth in the hippocampus of mice that were exposed to chronic stress (Besnard et al., 2018). In a study of juvenile frogs, shaking/confinement and intraperitoneal injections of CORT increased *KLF9* mRNA expression in the brain (Bonett et al., 2009). Exposing frogs to a stressor and, subsequently, to a GR antagonist decreased expression of *KLF9* while treatment with a mineralocorticoid antagonist had no effect (Bonett et al., 2009). This work suggests that CORT and GR act to regulate the actions of *KLF9* (Knoedler and Denver, 2014). Little work has been done on this gene in the pituitary, but as an immediate early gene that appears to respond to thyroid hormones, CORT, and other steroid hormones, it may also act to regulate these hormones in the pituitary. | -1.17 | 4.13E-03 | -0.87 | 3.66E-03 |
| 429116 | *PTGER3* | prostaglandin E receptor 3 | Reported in Table 1 | -2.02 | 2.70E-05 | -1.82 | 7.91E-04 |
| 396459 | *PVALB* | parvalbumin | *PVALB* encodes parvalbumin and regulates Ca<sup>2+</sup> and Mg<sup>+</sup> binding activity (Filipovič et al., 2013). Cells containing parvalbumin can act as GABAergic interneurons, which have been found to be associated with the stress response in the hippocampus (Filipovič et al., 2013). In response to chronic high CORT levels, parvalbumin containing cells decreased in the hippocampus of rodents; however, in response to acute stress, Pv-containing cells increased (Filipovič et al., 2013). The response of *PVALB* to glucocorticoids or restraint stress in the pituitary has not been studied. | 2.92 | 2.60E-04 | 3.49 | 9.54E-05 |
| 100859468 | *SLAINL* | SLAIN motif-containing protein-like | Unknown | -1.15 | 1.12E-04 | -1.12 | 4.92E-03 |
| 419759 | *ZBTB16* | zinc finger and BTB domain containing 16 | Described in Table 1 | -2.31 | 9.47E-09 | -2.16 | 2.95E-05 |

**Table 3:** Differential gene expression (DGE) in the pituitary in females during the stress response due to elevated circulating CORT concentrations. Red indicates upregulation while blue indicates downregulation.

| Entrez ID | Gene | Gene Name | Reported function | CORT |  | Restraint stress |  |
| --- | --- | --- | --- | --- | --- | --- | --- |
|  |  |  |  | logFC | FDR | logFC | FDR |
| 431627 | *ACOT12* | acyl-CoA thioesterase 12 | *ACOT12* encodes acyl-coenzyme A thioesterase 12 (also known as StAR-related lipid transfer protein 15). It is involved in a number of functions including acetyl-CoA hydrolase activity (GO:0003986). (O'Leary et al., 2016) | -2.53 | 7.93E-03 | -2.17 | 4.96E-05 |
| 420738 | *ANLN* | anillin actin binding protein | *ANLN* encodes an actin-binding protein that contributes to cytokinesis and cell growth and migration (O'Leary et al., 2016). | -4.89 | 1.71E-06 | -6.33 | 4.03E-11 |
| 419316 | *APCDD1L* | APC down-regulated 1 like | *APCDD1L* encodes a protein of unknown function. | -2.79 | 2.19E-07 | -1.43 | 4.75E-03 |
| 396536 | *APOA1* | apolipoprotein A1 | *APOA1* encodes the protein, apolipoprotein A-I, which is associated with high-density lipoprotein and acts in the transport of cholesterol and phospholipid (Sirniö et al., 2017). APOA1 also functions as an anti-inflammatory and antioxidant (Sirniö et al., 2017). | -2.29 | 6.29E-03 | -4.13 | 3.08E-07 |
| 424893 | *APOD* | apolipoprotein D | *APOD* encodes apolipoprotein D, which can bind to arachidonic acid, progesterone, retinol, cholesterol, among others (Ganfornina et al., 2008). It appears to be a stress responsive gene that acts in mediating oxidative stress (Ganfornina et al., 2008). | -3.58 | 1.22E-03 | -6.76 | 5.12E-10 |
| 417431 | *APOH* | apolipoprotein H | *APOH* encodes apolipoprotein H, which is implicated in a number of physiological processes (e.g., lipoprotein metabolism and coagulation) though its specific function is unknown (O'Leary et al., 2016). | -5.67 | 3.72E-07 | -6.98 | 1.48E-09 |
| 769889 | *APOLD1* | apolipoprotein L domain containing 1 | *APOLD1* is a stress-responsive immediate early gene that appears to be regulated by beta-adrenergic activity (Roszkowski et al., 2016). It functions in angiogenesis (Regard, 2004). | -1.96 | 4.77E-05 | -2.79 | 1.26E-06 |
| 421088 | *AQP4* | aquaporin 4 | *AQP4* encodes an aquaporin protein that functions in water homeostasis (O'Leary et al., 2016). | -2.89 | 2.32E-04 | -4.10 | 2.42E-08 |
| 421369 | *ATF3* | activating transcription factor 3 | *ATF3* encodes a member of the activating transcription factor/cyclic adenosine monophosphate(cAMP)-response element binding (CREB) protein family (Jadhav and Zhang, 2017). It is associated with cell stress and can act as a transcriptional activator or repressor, depending on context (Jadhav and Zhang, 2017). | -1.82 | 2.50E-04 | -2.96 | 2.76E-08 |
| 422966 | *C5H11ORF9* | chromosome 5 open reading frame, human C11orf9 | *C5H11ORF9* or the myelin regulatory factor (*MYRF*) in humans encodes a transcription factor associated with a key regulator of myelination and may directly stimulate gene expression of myelin (Bujalka et al., 2013). | -4.23 | 6.29E-05 | -5.59 | 8.78E-11 |
| 423959 | *C6H10ORF90* | chromosome 6 open reading frame, human C10orf90 | *C6H10ORF90* is a p53 responsive gene that acts in tumor suppression (Zhang et al., 2010). It is unclear what role this gene is playing here. | -2.14 | 8.04E-03 | -4.13 | 2.74E-10 |
| 416927 | *CA15L* | carbonic anhydrase 15-like | *CA15L* encodes a protein of unknown function. | -5.56 | 2.97E-07 | -5.85 | 9.73E-06 |
| 421029 | *CDH19* | cadherin 19 | *CDH19* encodes a calcium dependent cell-cell adhesion protein (O'Leary et al., 2016). | -3.91 | 4.48E-05 | -4.50 | 1.07E-09 |
| 107052707 | *CEBPD* | CCAAT enhancer binding protein delta | Described in Table 1 | -1.63 | 1.25E-06 | -1.89 | 1.73E-05 |
| 395335 | *CISH* | cytokine inducible SH2 containing protein | Described in Table 1 | -1.64 | 3.72E-07 | -1.81 | 1.91E-10 |
| 395399 | *CITED2* | Cbp/p300 interacting transactivator with Glu/Asp rich carboxy-terminal domain 2 | *CITED2* encodes a protein that competitively binds to hypoxia-inducible factor 1-alpha (*HIF1A*) thereby inhibiting expression of *HIF1A*-induced genes (O'Leary et al., 2016). | -1.38 | 7.43E-07 | -0.90 | 7.93E-03 |
| 395921 | *CNP* | 2',3'-cyclic nucleotide 3' phosphodiesterase | *CNP* encodes a protein that encodes a precursor of *NPCC* (C-type natriuretic peptide). *NPCC* is associated with vasodilation and natriuresis. (O'Leary et al., 2016) | -3.15 | 1.69E-05 | -5.09 | 4.87E-12 |
| 428435 | *CREB5* | cAMP responsive element binding protein 5 | *CREB5* encodes a cAMP response element (CRE) binding protein (Yu et al., 2018). *CREB5* regulates cell growth, proliferation, and differentiation, and may be involved in the immune response (Yu et al., 2018). Recent work showed that hypo-methylation and upregulation of *CREB5* in decidua was associated with recurrent pregnancy loss in humans (Yu et al., 2018). | -1.97 | 7.07E-03 | -3.97 | 4.16E-07 |
| 421422 | *DAAM2* | dishevelled associated activator of morphogenesis 2 | *DAAM2* helps regulate the planar cell polarity pathway, a Wnt signaling pathway (Lee et al., 2015). *Daam2* appears to inhibit the differentiation of oligodendrocytes thereby impacting myelination (via the PIP5K-PIP2 axis; Lee et al., 2015). | -1.50 | 7.69E-03 | -2.71 | 1.99E-08 |
| 107053650 | *DDIT4* | DNA damage inducible transcript 4 | *DDIT4* is a stress responsive gene that suppresses mTor (Mechanistic Target of Rapamycin) pathways (Canal et al., 2014; Polman et al., 2012). Upregulation of *DDIT4* occurs following cellular damage associated with a number of stressors (Canal et al., 2014; Polman et al., 2012). *DDIT4* also responds to the presence of glucocorticoids via a GRE-like element on its promoter (Polman et al., 2012). Upregulation of *DDIT4* results in the suppression of the mTOR pathway, which has been associated with reduced cell growth and plasticity in several tissues and an increase in cell apoptosis (Polman et al., 2012). Most of the research on this gene has focused on the hippocampus—its role in the pituitary has not been well-studied. | -1.49 | 1.21E-08 | -2.75 | 9.40E-08 |
| 425241 | *DIRAS2* | DIRAS family GTPase 2 | *DIRAS2* likely acts in GTP binding (GO:0005525) and GTPase activity (GO:0003924). | -1.58 | 1.71E-04 | -1.41 | 4.23E-03 |
| 422551 | *ENPP6* | ectonucleotide pyrophosphatase/phosphodiesterase 6 | *ENPP6* is thought to be involved in extracellular nucleotide metabolism regulation and myelination (O'Leary et al., 2016). | -2.37 | 2.03E-03 | -2.78 | 3.56E-06 |
| 771826 | *ERMN* | ermin | *ERMN* encodes a protein that is associated with myelination and mature nerves (Brockschneider, 2006). | -5.22 | 1.69E-05 | -4.37 | 2.65E-08 |
| 426690 | *ETNK2* | ethanolamine kinase 2 | *ETNK2* is associated with early catalysis of the ethanolamine pathway (O'Leary et al., 2016). | -2.64 | 7.83E-04 | -4.16 | 1.90E-08 |
| 415687 | *FA2H* | fatty acid 2-hydroxylase | *FA2H* encodes the enzyme, fatty acid 2-hydroxylase (O'Leary et al., 2016). It may be involved in fatty acid modification and myelin maintenance (O'Leary et al., 2016). | -4.68 | 1.33E-05 | -5.36 | 1.78E-10 |
| 422413 | *FAM198B* | family with sequence similarity 198 member B | *FAM198B* is predominately expressed in the adrenal and ovary (O'Leary et al., 2016). The function of this gene is currently unknown. | -3.14 | 3.01E-03 | -2.93 | 3.72E-03 |
| 419084 | *GDPD4* | glycerophosphodiester phosphodiesterase domain containing 4 | Little information is known about *GDPD4*, but it is involved in glycerophosphodiester phosphodiesterase activity (GO:0008889), metal ion binding (GO:0046872), and the lipid metabolic process (GO:0006629) (O'Leary et al., 2016). | -2.29 | 4.25E-04 | -2.16 | 2.78E-03 |
| 419969 | *GFAP* | glial fibrillary acidic protein | *GFAP* encodes glial fibrillary acidic protein, which is an intermediate filament protein found in a number of cell types in the CNS (e.g., astrocytes) (O'Leary et al., 2016). | -5.98 | 8.37E-05 | -7.93 | 9.40E-12 |
| 420397 | *GJC2* | gap junction protein gamma 2 | *GJC2* encodes a gap junction protein that is involved in myelination (O'Leary et al., 2016). | -1.65 | 4.42E-03 | -2.93 | 4.70E-08 |
| 427523 | *GLDN* | gliomedin | *GLDN* encodes an olfactomedin-like and collagen-like domain protein that can occur in both transmembrane and secreted forms. It plays a role in the formation of the nodes of Ranvier in the peripheral nervous system. (O'Leary et al., 2016) | -3.10 | 9.43E-05 | -1.43 | 8.56E-03 |
| 396489 | *GLUL* | glutamate-ammonia ligase | *GLUL* encodes a member of the glutamine synthetase family, which plays a role in the catalysis of glutamine synthesis and glutamate detoxification, cell signaling and proliferation, and acid-base homeostasis (O'Leary et al., 2016). | -1.87 | 1.10E-03 | -4.03 | 7.05E-11 |
| 417748 | *GPR37* | G protein-coupled receptor 37 | *GPCR* and *GPCRL1* are associated with the regulation of neuronal and glial physiology and modulate dopaminergic neurotransmission (Meyer et al., 2013). These receptors are bound by prosaposin and prosaptide, which stimulates ERK phosphorylation and protect against cellular stress (Meyer et al., 2013). Prosaposin has neuroprotective and glioprotective action, which is likely mediated by these receptors (Meyer et al., 2013). | -3.32 | 2.28E-03 | -4.80 | 2.76E-08 |
| 421176 | *GPR37LI* | G protein-coupled receptor 37 like 1 | *GPCR* and *GPCRL1* are associated with the regulation of neuronal and glial physiology and modulate dopaminergic neurotransmission (Meyer et al., 2013). These receptors are bound by prosaposin and prosaptide, which stimulates ERK phosphorylation and protect against cellular stress (Meyer et al., 2013). Prosaposin has neuroprotective and glioprotective action, which is likely mediated by these receptors (Meyer et al., 2013). | -3.92 | 1.29E-03 | -5.14 | 2.43E-07 |
| 100859224 | *GPR62* | G protein-coupled receptor 62 | Little is known about *GPCR62*, but this gene encodes a protein that may regulate cAMP production (Muroi et al., 2017). | -5.86 | 6.93E-05 | -6.58 | 1.94E-09 |
| 425975 | *HAPLN2* | hyaluronan and proteoglycan link protein 2 | *HAPLN2* encodes a hyaluronan and proteoglycan binding link protein that is acts in the stabilization and increase of binding of four chondroitin sulfate proteoglycans core proteins and in maintaining the extracellular matrix (Wang et al., 2016). It is expressed in myelinated fiber in the brain and overexpression in rats was associated with cell death of neurons (Wang et al., 2016). | -5.40 | 3.01E-03 | -7.37 | 7.05E-11 |
| 424441 | *HEBP2* | heme binding protein 2 | *HEBP2* encodes a protein that is associated with the collapse of mitochondrial membrane potential in the cytoplasm and enhances mitochondrial membrane permeability during oxidative stress (O'Leary et al., 2016). | -2.54 | 1.95E-03 | -3.30 | 4.98E-10 |
| 428234 | *HEPACAM* | hepatic and glial cell adhesion molecule | *HEPACAM* encodes a protein that functions in cell motility or cell-matrix interaction (O'Leary et al., 2016). | -3.88 | 6.93E-05 | -4.03 | 2.55E-06 |
| 395128 | *HES4* | hes family bHLH transcription factor 4 | *HES4* is associated with the Notch signaling pathway (McManus et al., 2017). | 1.44 | 3.68E-06 | 1.40 | 3.70E-06 |
| 420476 | *JCAD* | junctional cadherin 5 associated | *JCAD* or *KIAA1462* encodes a cell-cell junction protein (O'Leary et al., 2016). | -1.08 | 8.23E-03 | -1.67 | 1.54E-04 |
| 427662 | *KCNJ12* | potassium voltage-gated channel subfamily J member 12 | *KCNJ12* encodes an inwardly rectifying K<sup>+</sup> channel (O'Leary et al., 2016). | -3.09 | 5.26E-03 | -4.24 | 1.43E-06 |
| 395925 | *KCNJ5* | potassium voltage-gated channel subfamily J member 5 | Described in Table 1 | -1.74 | 4.09E-06 | -1.55 | 4.09E-03 |
| 422374 | *KIAA1210* | *KIAA1210* | *KIAA1210* may be a cell junction protein involved with mammalian spermiogenesis in the testes, but little else is known about its function in other tissues (Iwamori et al., 2017). | -1.29 | 2.34E-03 | -1.48 | 1.33E-04 |
| 395705 | *LHX2* | LIM homeobox 2 | *LHX2* functions in transcription regulation (O'Leary et al., 2016). | -2.91 | 1.29E-03 | -2.42 | 3.16E-03 |
| 100859848 | *LOC100859848* | CDC42 small effector protein 2-C-like | *LOC100859848* encodes a protein of unknown function. | -2.20 | 3.28E-04 | -3.26 | 3.32E-09 |
| 107050516 | *LOC107050516* | Schwann cell myelin protein-like | *LOC107050516* encodes a protein of unknown function. | -3.62 | 7.88E-06 | -4.40 | 2.00E-07 |
| 769726 | *LOC769726* | kazal-type serine protease inhibitor domain-containing protein 1-like | *LOC769726* encodes a protein of unknown function. | -5.64 | 1.15E-06 | -6.93 | 1.81E-10 |
| 396217 | *MBP* | myelin basic protein | *MPB* encodes myelin basic protein, which is a critical component of the myelin sheath of oligodendrocytes and Schwann cells in the CNS and in myelination (O'Leary et al., 2016). | -1.81 | 1.77E-03 | -3.92 | 4.37E-12 |
| 417737 | *MLC1* | megalencephalic leukoencephalopathy with subcortical cysts 1 | *MLC1* encodes a product of unknown function (O'Leary et al., 2016). | -2.15 | 9.18E-04 | -3.98 | 2.76E-08 |
| 418151 | *NINJ2* | ninjurin 2 | *NINJ2* encodes a protein of the ninjurin (for nerve injury induced) family, which is a cell surface adhesion molecule that is related to nerve injury and neurite outgrowth (Wang et al., 2017). It has also been suggested as a regulator of multiple proteins associated with inflammation (GDNF, GM-CSF, ICAM-1, IL-1β, M-CSF, TGF-β1, TGF-β3, TNF-α, TNF-β, BLC, CCL28, Fractalkine, GCP-2, I-TAC, IL-8, Lymphotoxin, MIP-3β, MIP-3α and TECK) and may influence the gene expression of several associated genes (*GDNF*, *ICAM-1*, *IL-1β*, *M-CSF*, *TGF-β1*, *TGF-β3*, *BLC*, *CCL28*, *Fractalkine*, *GCP-2*, *I-TAC*, *IL-8*, *Lymphotoxin*, *MIP-3β*, *MIP-3α* and *TECK*), as well (Wang et al., 2017). *NINJ2* and *TLR4* also regulate NF-KB and c-jun pathways (Wang et al., 2017). | -5.13 | 1.15E-06 | -6.08 | 7.05E-11 |
| 421833 | *NT5E* | 5'-nucleotidase ecto | *NT5E* encodes a plasma membrane protein that can act to inhibit immune response via its generation of a adenosine (Kordaß et al., 2018). | -4.37 | 1.93E-04 | -3.29 | 4.39E-06 |
| 428612 | *OLIG2* | oligodendrocyte transcription factor 2 | *OLIG2* encodes a transcription factor that regulates oligodendrocyte progenitor cells, ventral neuroectodermal progenitor cell fate and chromosomal translocation (Bujalka et al., 2013; O'Leary et al., 2016). Oligodendrocytes act in CNS myelination (Bujalka et al., 2013). | -5.52 | 3.28E-05 | -7.78 | 1.80E-11 |
| 395334 | *OPN4-1* | photopigment melanopsin-like | *OPN4-1* is associated with melanopsin biosynthesis and may contribute to circadian entrainment in the tissue of expression (Bailey and Cassone, 2005). | -4.43 | 2.97E-03 | -5.74 | 3.54E-07 |
| 771808 | *PIPOX* | pipecolic acid and sarcosine oxidase | *PIPOX* is associated with L-pipecolate oxidase activity (GO:0050031), sarcosine oxidase activity (GO:0008115), and signaling receptor binding (GO:0005102). | -3.60 | 1.89E-03 | -5.09 | 9.15E-05 |
| 415650 | *PLLP* | plasmolipin | *PLLP* functions in myelin biosynthesis though little more is known about this protein (Yaffe et al., 2015). | -4.55 | 5.36E-06 | -7.10 | 4.88E-11 |
| 396214 | *PLP1* | proteolipid protein 1 | *PLP1* encodes a key component of myelin and may be associated with various functions related to myelin (O'Leary et al., 2016). | -8.93 | 3.52E-07 | -10.69 | 4.37E-12 |
| 420198 | *PMP2* | peripheral myelin protein 2 | *PMP2* encodes a protein that is associated with myelin sheaths in the peripheral nervous system that may act in sheath stabilization (O'Leary et al., 2016). | -7.36 | 2.32E-04 | -4.79 | 3.77E-04 |
| 417327 | *PMP22* | peripheral myelin protein 22 | *PMP22* encodes an integral membrane protein of myelin and the peripheral nervous system (O'Leary et al., 2016). | -2.66 | 6.29E-04 | -2.27 | 2.16E-07 |
| 429116 | *PTGER3* | prostaglandin E receptor 3 | Described in Table 1 | -1.99 | 2.19E-07 | -1.51 | 7.50E-03 |
| 419521 | *RASSF2* | Ras association domain family member 2 | *RASSF2* functions in cell growth inhibition, cell cycle arrest, actin cytoskeleton organization, apoptosis, suppression of transcription of NFkB, and inhibition of *MST2* activity (Volodko et al., 2014). | -3.05 | 1.02E-04 | -4.17 | 9.38E-09 |
| 424038 | *S100B* | S100 calcium binding protein B | *S100B* has been linked to cell survival, proliferation and differentiation, neuromodulation, cell migration, cell morphogenesis and proliferation, and cytoskeletal dynamic, depending on its effector (Dempsey et al., 2016). | -4.31 | 4.90E-05 | -4.45 | 5.98E-08 |
| 424593 | *SEPPIL* | selenoprotein P2 | The function of *SEPIL* is unknown, but it may act as an extracellular antioxidant (O'Leary et al., 2016). | -4.12 | 5.86E-05 | -5.43 | 5.29E-09 |
| 416778 | *SEPT5* | septin 5 | *SEPT5* encodes a nucleotide binding protein that is associated with regulation of cytoskeletal organization (O'Leary et al., 2016). | -1.81 | 1.05E-03 | -3.32 | 7.05E-11 |
| 107049626 | *SFTPC* | SFTPC surfactant protein C | *SFTPC* encodes surfactant protein C, which is a protein required for lung function (O'Leary et al., 2016). It is unclear why it is differentially expressed in the female pituitary as expression is thought to be restricted to lungs (O'Leary et al., 2016). | -4.17 | 7.93E-03 | -6.18 | 8.09E-06 |
| 418731 | *SH3RF3* | SH3 domain containing ring finger 3 | *SH3RF3* encodes a scaffold protein associated with E3 ligase activity, which may be involved in JNK-mediated apoptosis (Kärkkäinen et al., 2010). | -2.12 | 1.90E-03 | -1.86 | 1.38E-03 |
| 396089 | *SHANK3* | SH3 and multiple ankyrin repeat domains 3 | *SHANK3* is involved in synapse function, specifically as a scaffold protein involved in connecting neurotransmitter receptors and ion channels to G protein receptor signaling pathways or actin cytoskeleton (Boeckers et al., 2002; O'Leary et al., 2016). It is also involved in dendritic spine morphology (Boeckers et al., 2002). | -5.62 | 2.21E-04 | -7.31 | 1.24E-10 |
| 395615 | *SHH* | sonic hedgehog | *SHH* encodes a protein involved in the hedgehog pathway. Most notably associated with embryonic development, SHH could play a role in neurogenesis, and the maintenance of homeostasis and repair of neural stem cell progenitors in adult tissues (Petrova and Joyner, 2014). | 1.95 | 9.50E-03 | -2.66 | 7.57E-04 |
| 424576 | *SLC6A9* | solute carrier family 6 member 9 | *SLC6A9* encodes a protein that inhibit glycine signaling, a neurotransmitter inhibitor of the CNS (O'Leary et al., 2016). | -2.93 | 6.29E-05 | -4.37 | 2.29E-09 |
| 432368 | *SNAI2* | snail family transcriptional repressor 2 | *SNAI2* encodes a transcription factor that is associated with embryonic development, but it is also suggested to play a role in maintaining normal function in mature cells as it is found in most tissues (O'Leary et al., 2016). | 1.48 | 2.28E-03 | 1.37 | 1.41E-03 |
| 395573 | *SOX10* | SRY-box 10 | *SOX10* is a transcription factor typically associated with embryonic development. However, in adult cells, Sox10 complexes with Olig10 to activate mbp (myelin basic protein) transcription. (Li et al., 2007) | -4.46 | 5.99E-04 | -6.84 | 6.17E-09 |
| 395483 | *SOX8* | SRY-box 8 | *SOX8* is a SRY-related HMG-box transcription factor, which is associated with embryonic development regulation and cell fate determination (O'Leary et al., 2016). | -2.31 | 5.01E-05 | -3.74 | 1.55E-09 |
| 419414 | *TMEM88B* | transmembrane protein 88B | *TMEM88B* encodes a protein with a currently unknown function. | -5.07 | 1.69E-04 | -5.69 | 1.55E-09 |
| 396032 | *TNNC1* | troponin C1, slow skeletal and cardiac type | *TNNC1* encodes a regulatory protein associated with muscle contraction in the actin filament (O'Leary et al., 2016). | -4.79 | 2.88E-05 | -5.67 | 2.00E-06 |
| 768091 | *TSC22D3* | TSC22 domain family member 3 | *TSC22D3* encodes an anti-inflammatory protein glucocorticoid-induced leucine zipper that is stimulated by glucocorticoids and interleukin-10. It functions in inhibiting inflammation and the immune system. (O'Leary et al., 2016) | -1.02 | 1.22E-03 | -1.65 | 1.16E-06 |
| 428900 | *TSHR* | thyroid stimulating hormone receptor | *TSHR* is a gene that transcribes the thyroid stimulating hormone receptor. The thyroid stimulating hormone receptor responds to the presence of thyroid stimulating hormone, a metabolic hormone that stimulates the thyroid to produce thyroxine (T4) and triiodothyronine (T3). Other studies have shown that glucocorticoids increase expression of *TSHR* in the pituitary (Mariotti and Beck-Peccoz, 2000). While this gene was upregulated, there was high variance, which suggests a trend towards increased expression of *TSHR* (Fig. 6). While pharmaceutical doses of glucocorticoids in rats can suppress circulating TSH, mRNA *TSH*, and thereby, *TSHR*, this study found that *TSHR* was upregulated (Mariotti and Beck-Peccoz, 2000). This result may indicate an increase in metabolism in response to glucocorticoids and restraint stress. | -3.13 | 8.29E-03 | -3.66 | 3.70E-05 |
| 100858879 | *TTYH2* | tweety family member 2 | *TTYH2* encodes a protein that influences calcium (2+)-activated large conductance chloride(-) channels (O'Leary et al., 2016). | -1.79 | 4.79E-03 | -3.86 | 1.72E-09 |
| 374033 | *UGT8* | UDP glycosyltransferase 8 | *UGT8* encodes 2-hydroxyacylsphingosine 1-beta-galactosyltransferase, which is associated with myelin (Bosio et al., 1996). | -2.67 | 3.53E-04 | -3.50 | 2.49E-09 |
| 419759 | *ZBTB16* | zinc finger and BTB domain containing 16 | Described in Table 1 | -1.66 | 1.25E-06 | -2.86 | 5.57E-08 |

**Table 4:**
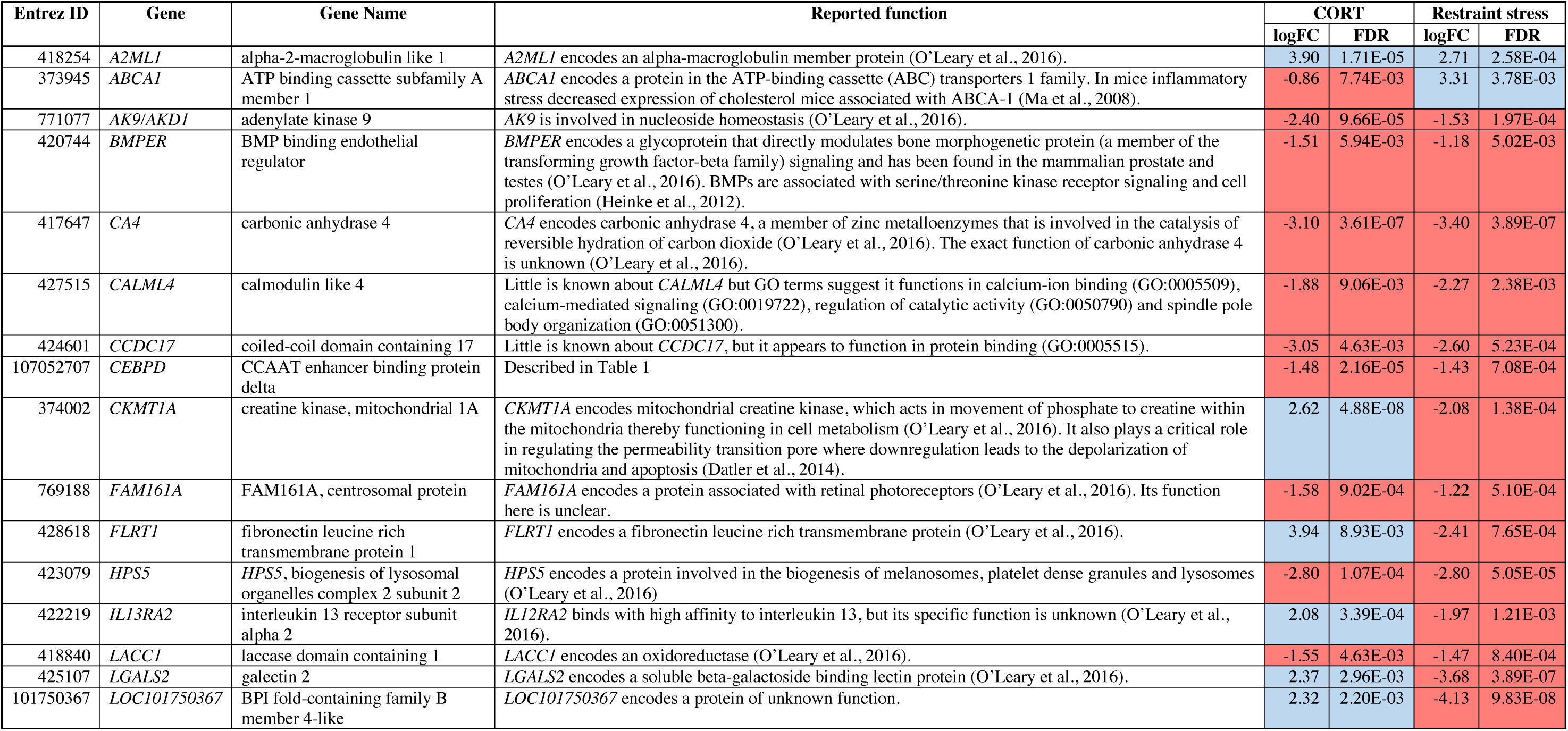
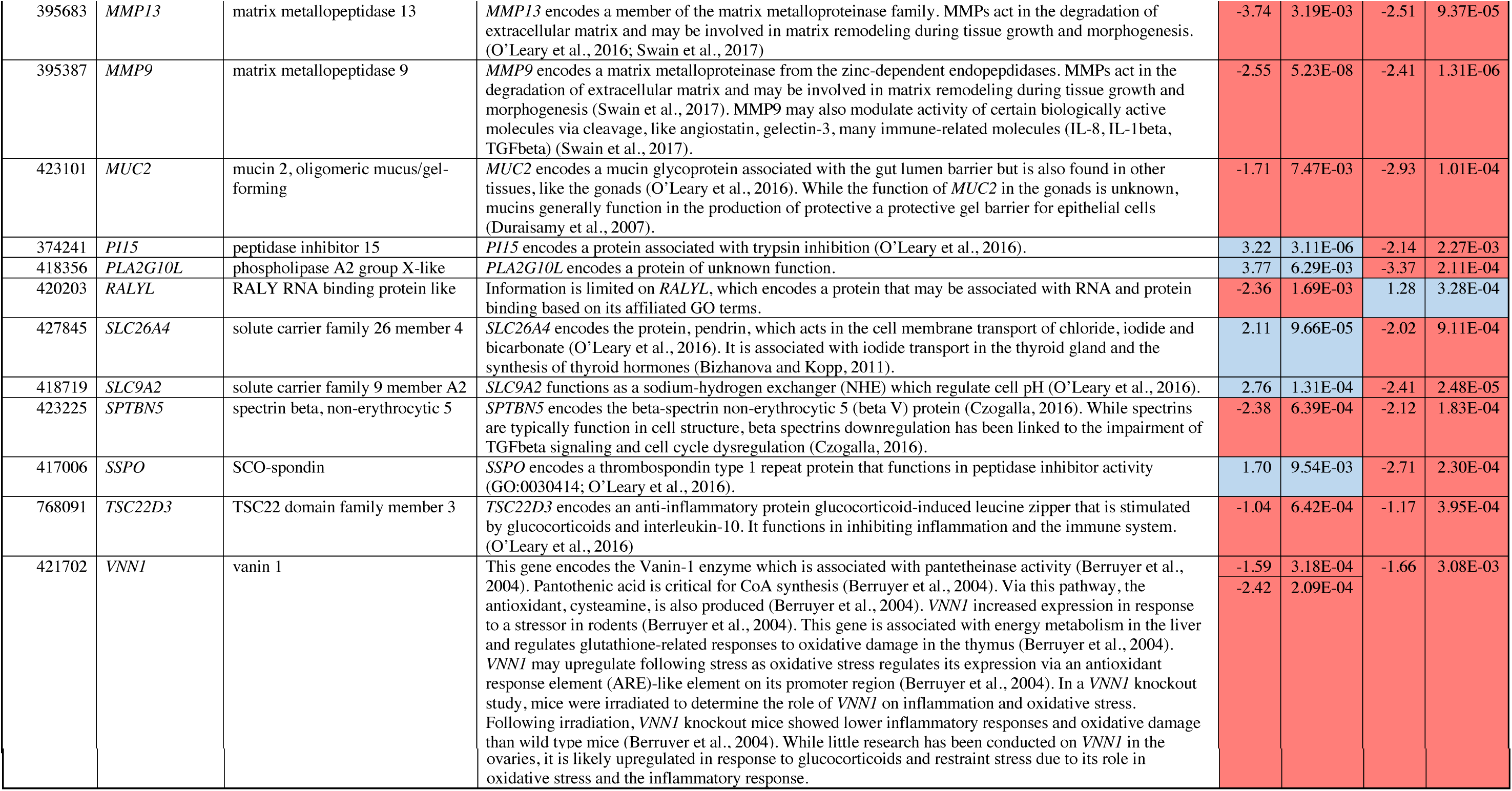
Differential gene expression (DGE) in the ovaries due to elevated circulating CORT concentrations. Red indicates upregulation while blue indicates downregulation.

#### Candidate Gene Expression Evaluation

To detect more subtle changes in gene expression, we conducted a separate analysis to include only a select group of *a priori*-targeted genes (*N* = 44) based on their previously known role in the stress response or reproductive function (Fig. 6; Table 5). We identified two candidate genes, *DIO2* in the female hypothalamus and *PRLR* in the male hypothalamus, for which expression varied in response to 30 min of restraint stress could be explained by elevated CORT.

**Table 5:**
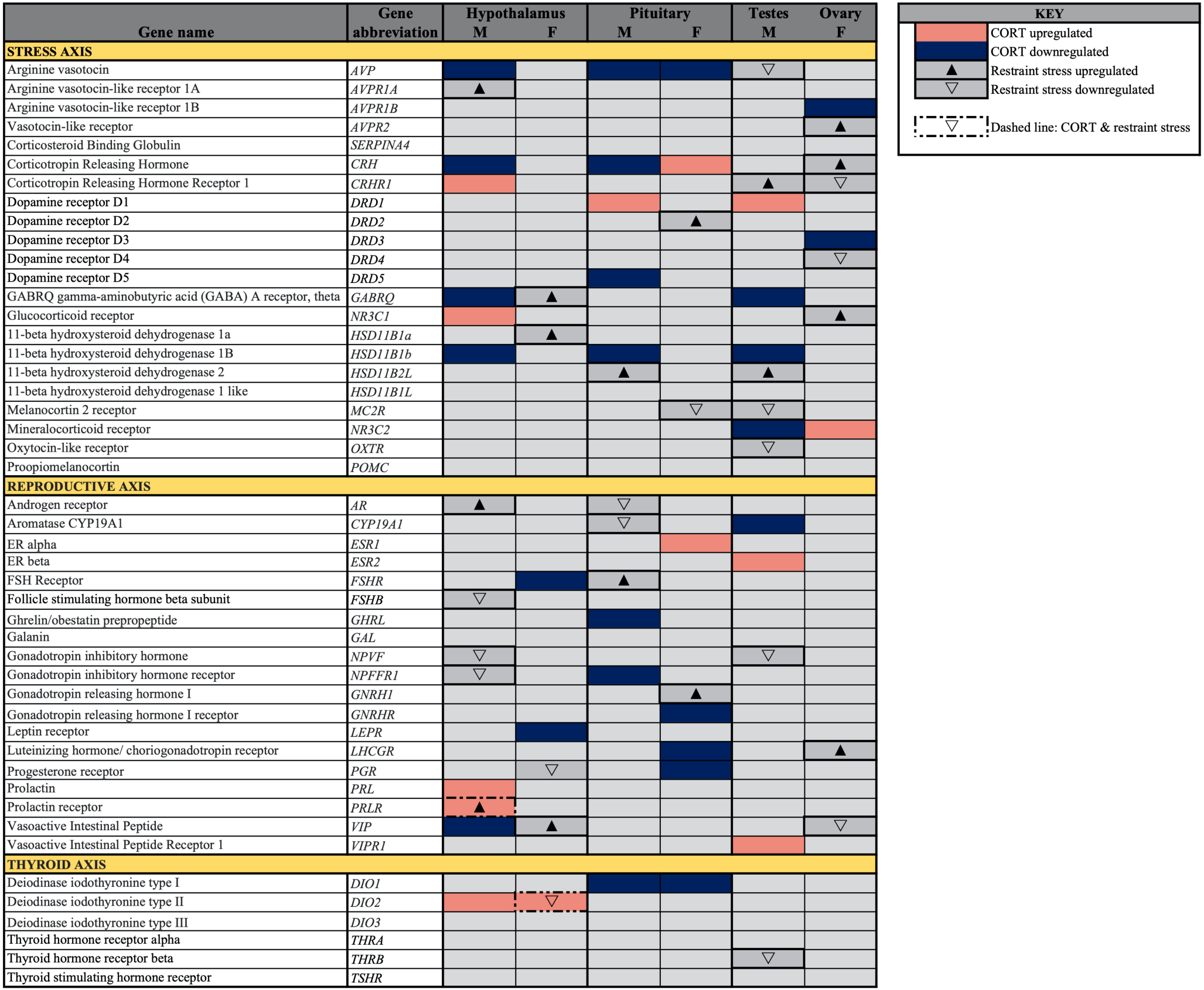
*A priori*-identified stress and reproduction-associated target genes that differentially express in response to CORT and restraint stress treatments (α= 0.05). Pink denotes upregulated activity in CORT-treated birds and dark blue denotes downregulated activity in CORT-treated birds. Solid upward facing triangles denote upregulated in restraint stress birds, and open downward facing triangles denote downregulated in restraint stress birds. Dashed lines denote a significant change in gene expression in both restraint stress and CORT-treated birds.

**Figure 5:**
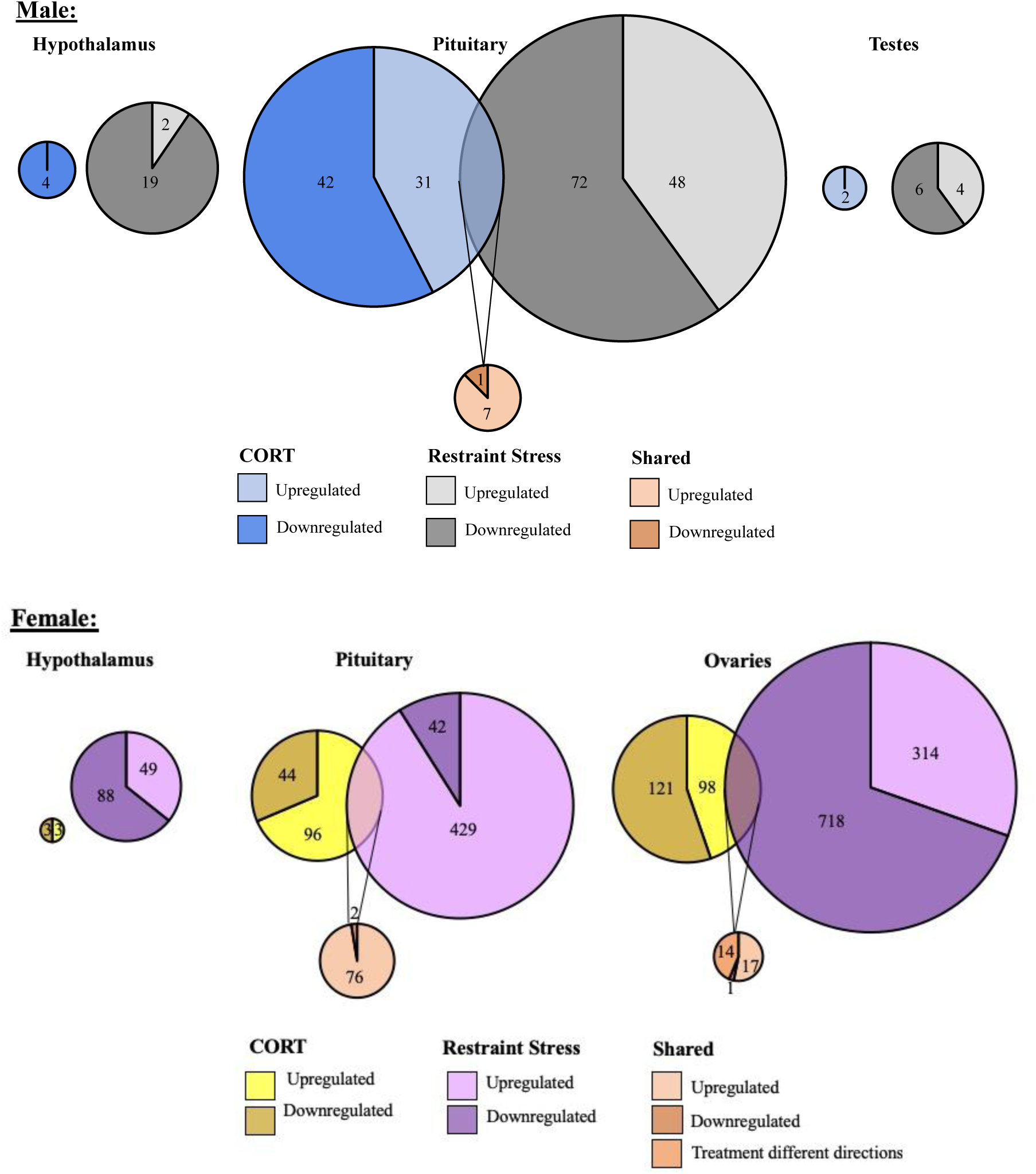
Shared and unique gene expression responses to CORT treatment and restraint-stress. Weighted Venn diagrams of significantly differentially expressed genes for male (top) and female (bottom) of CORT-treated and restraint stress treatments (data from Calisi et. al. 2018) in the hypothalamus, pituitary and gonads. Circles are proportional within males and females, but not between.

**Figure 6:**
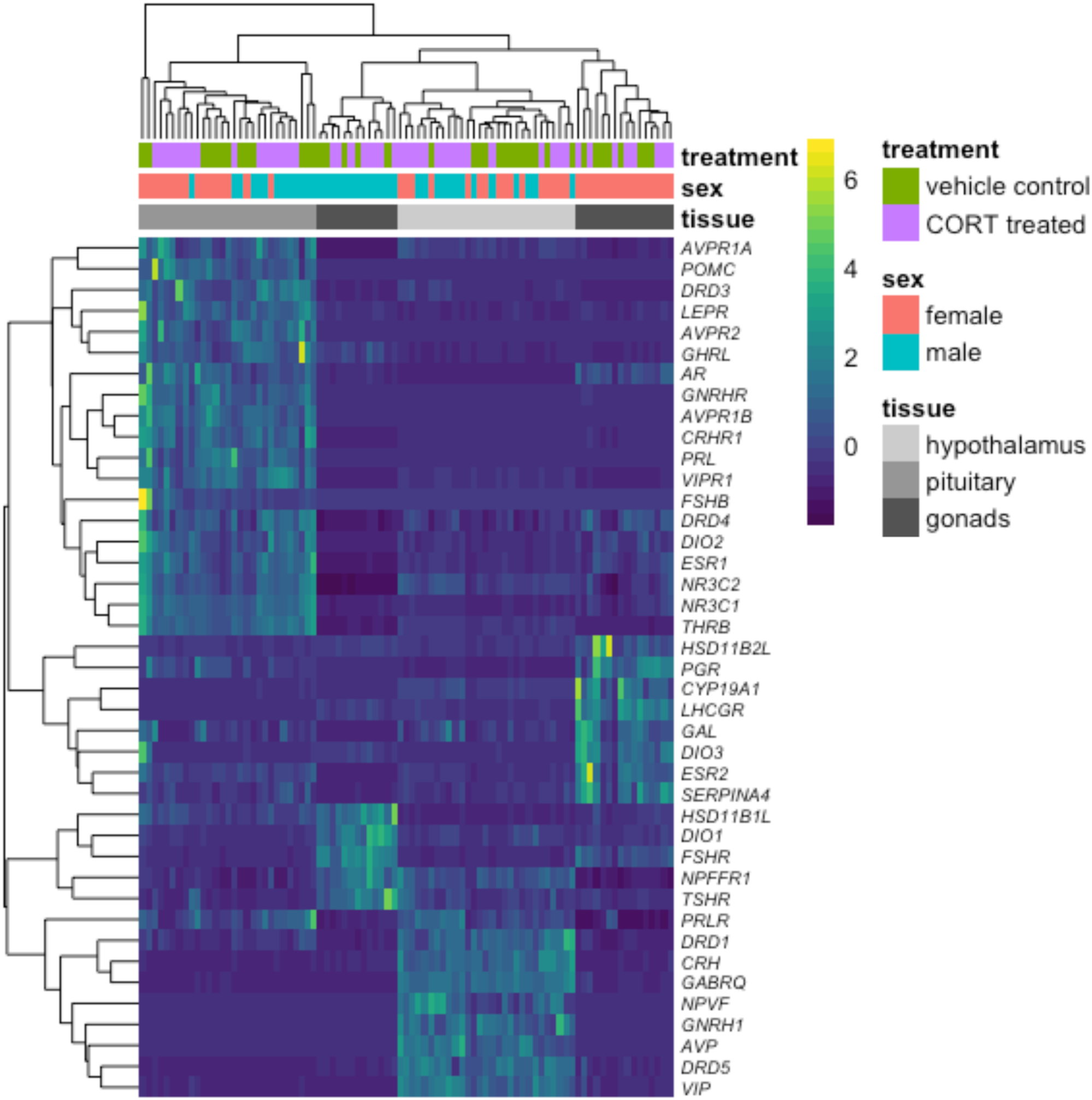
A heatmap depicting individual candidate gene expression values. (normalize gene counts) represented as colors. Blue shades signify low levels of expression; Yellow shades signify relatively higher levels of expression. The *Y*-axis on the right denotes the gene abbreviation; the *Y*-axis on the left is a dendrogram of expression similarity. The *X*-axis denotes the sex, tissue, and treatment group of the animal (CORT-treated or the vehicle control).

**Figure 7.**
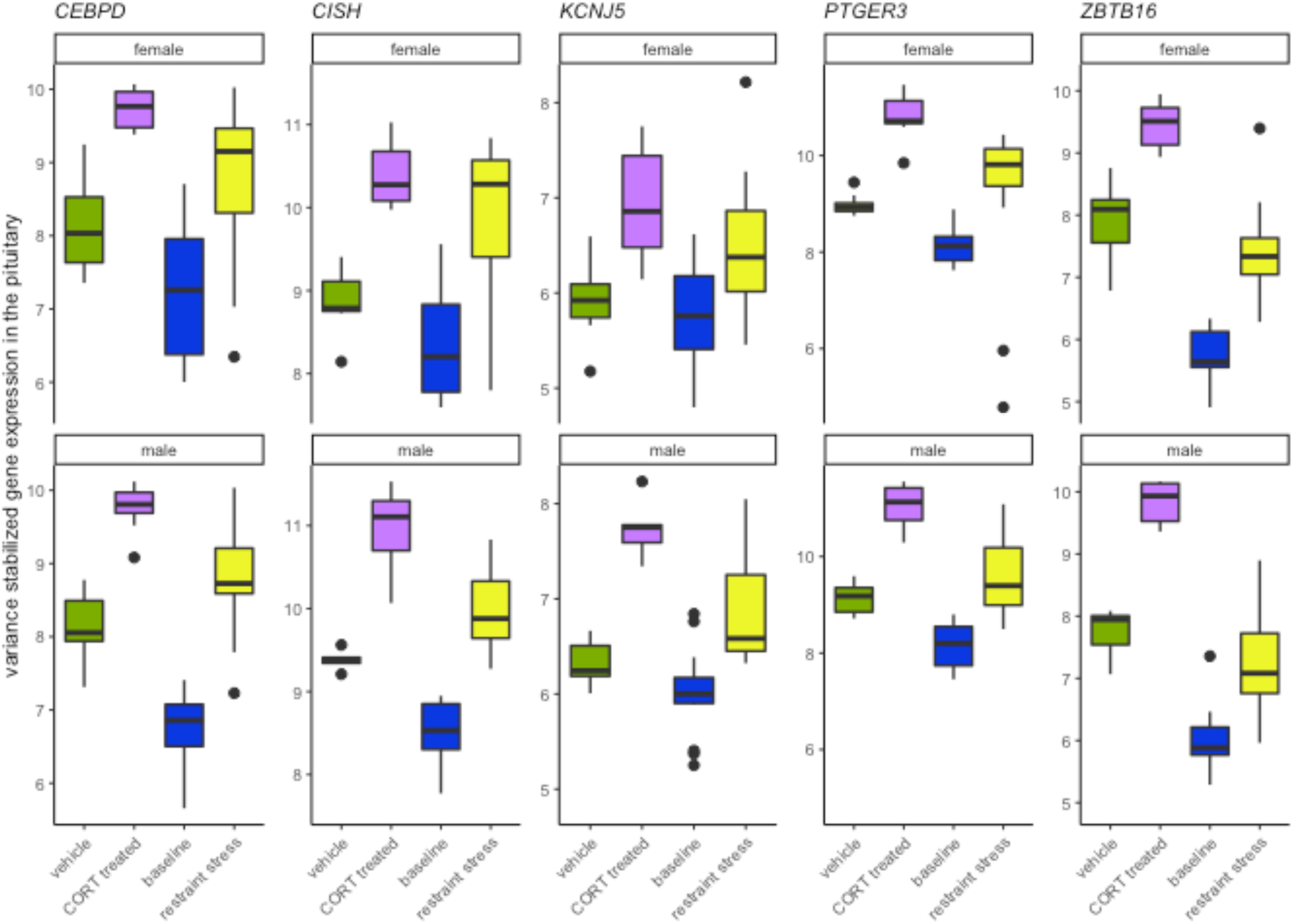
Genes that increase in response to both CORT and stress in both males and females. The elevation of CORT during the stress response resulted in the differential expression in both sexes of *KCNJ5, CISH, PTGER3, CEBPD*, and *ZBTB16* in the pituitary.

Most target genes responded uniquely to either exogenous CORT or restraint stress treatments. As compared to vehicle-injected controls, exogenous CORT treatment resulted in the differential response of 22 candidate genes: 12 in the hypothalamus (male: 10, female: 3), 12 in the pituitary (male: 8, female: 7), and 9 in the gonads (male: 7, female: 3). While restraint stress compared to its control resulted in the differential response of 26 candidate genes: 11 in the hypothalamus (male: 6, female: 5), 7 in the pituitary (male: 4, female: 3), and 12 in the gonads (male: 6, female: 7).

## DISCUSSION

A substantial body of research has been dedicated to understanding the critical role CORT plays via the stress response in promoting survival (Romero and Wingfield, 2015; Sapolsky et al., 2000; Wingfield et al., 1998; Wingfield and Kitaysky, 2002) and its subsequent consequences in other biological processes, like reproductive function (Angelier and Chastel, 2009; Calisi et al., 2018; Geraghty and Kaufer, 2015; Sapolsky et al., 2000). In this study, we experimentally tested the extent to which changes in gene activity in the HPG axis could be explained by an increase in circulating CORT that is characteristic of an acute stress response.

Using a common restraint stress paradigm, we previously reported its effects on HPG axis at the level of gene transcription (Calisi et al., 2018). To isolate the role of elevated CORT in causing these changes, we experimentally manipulated circulating CORT concentrations to mimic those experienced by our subjects during restraint stress. We identified genes that significantly altered their activity in response to both CORT-manipulated and restraint stress-treated groups as compared to their respective controls. We report activity changes in these genes as being indicative of the transcriptomic response to 30 min of restraint stress explained by elevated circulating CORT concentrations.

### Hypothalamus

We did not identify genes in the hypothalamus of either sex whose differential expression patterns as a result of restraint stress was related to elevated circulating CORT. Exogenous CORT administration fails to elicit the typical patterns of sensory integration in the hypothalamus and pituitary that result in the activation of pathways associated with fear, metabolism, anxiety, and stress. Based on this assumption and our understanding of HPA axis function (both activation and negative feedback) and HPG function, we selected *a priori* 47 candidate genes for targeted analyses with relatively small effect sizes. Elevated CORT concentrations during our restraint stress treatment were responsible for a heightened level of prolactin receptor (*PRLR)* transcription in males and a decrease in Type II iodothyronine deiodinase (*DIO2)* transcription in females. Among a myriad of other functions, prolactin also has a known role in immune and reproductive function. Increased expression of *PRLR* in the brain may act as a neurotransmitter or neuromodulator and may influence reproduction, metabolism, and/or nervous system function (Chaiseha et al., 2012). Exposure to short-term stressors may also increase prolactin, though increases in circulating levels occurs predominately in mammals whereas in birds prolactin levels tend to be unaffected or decrease following a stressor during breeding (Angelier et al., 2016; Angelier and Chastel, 2009; Romero and Wingfield, 2015). Administration of exogenous prolactin increased food intake, increased negative feedback of *PRL*, had antigonadotrophic effects, and reduced gonadal mass (Buntin et al., 1999; Buntin and Figge, 1988; Buntin and Tesch, 1985; Fechner and Buntin, 1989; Rozenboim et al., 1993). Prolactin may act to inhibit reproduction by suppressing GnRH in breeding birds. Increased prolactin in birds has been associated with decreased estradiol and ovarian regression in females and reduced gonadal growth in males (Buntin and Tesch, 1985). Because circulating prolactin may rise following a stressor, inhibition of reproduction by prolactin may be another means that birds can adjust to their local environment and potentially reduce the energetic costs of maintaining their reproductive state when environment is unsuitable for reproduction or to avoid an unsuccessful breeding attempt. Evidence in rats suggests that prolactin may inhibit the HPA reactivity to stress where suppression of *PRLR* increased ACTH secretion; it’s not clear if a similar effect occurs in birds (reviewed in Bole-Feysot et al., 1998; Cabrera-Reyes et al., 2017; Torner, 2016). Dio2 converts thyroxine (T4) into triiodothyronine (T3) and stimulates metabolism. In the hypothalamus, Dio2 serves as part of a negative feedback loop on thyroid releasing hormone (Helmreich and Tylee, 2011). Dio2 also plays a role in activating the reproductive axis by stimulating the production of triiodothyronine (T3) and promoting the release of gonadotropin releasing hormone (Pérez et al., 2018).

An alternative, albeit non-exclusive reason for the lack of a hypothalamic transcriptomic response to exogenous CORT may be due to its heterogeneous nature. The avian hypothalamus contains 19 nuclei, each characterized by form and function (Kuenzel and van Tienhoven, 1982). A result of this may be signal dilution, which could obstruct our identification of specific CORT-responsive substrates characteristic of the hypothalamic transcriptomic response. While this is a possibility, it is more likely that CORT treatment simply bypasses the primary action of the hypothalamus in the stress response.

### Pituitary

Elevated CORT during the stress response drives the differential expression of five genes in both sexes, all found in the pituitary: *KCNJ5, CISH, PTGER3, CEBPD*, and *ZBTB16* (Table 1). The rapid action properties of the immediate early gene *CISH*, genes that code for transcription factors, *CEBPD* and *ZBTB16*, or with a glucocorticoid response element (GRE) on its promotor region, *KCNJ5*, suggest their potential role in early responsiveness to the CORT signal of the stress response. *PTGER3* and *CEBPD* also encode proteins that regulate prostaglandin E2 (PGE_2_). PGE_2_ is a vasodilator and immunomodulator, and thus a change in *PTGER3* and *CEBPD* may be acting to suppress immune function in response to CORT. The presence of GREs in these shared genes and several sex-specific genes suggests a mechanism by which glucocorticoids could bind and quickly induce gene transcription.

A common functional theme that emerged was the role that the products of these 5 genes play in the inflammatory immune response (*CEBPD, CISH, PTGER3*) and in the modulation of prolactin (*CEBPD, CISH, KCNJ5*). The role of CORT in immune and inflammatory responses has been relatively well known and are mediated by GR activation (Sapolsky et al., 2000; Webster Marketon and Glaser, 2008). In brief, CORT can both activate and suppress immune function through multiple pathways (reviewed in Gao et al., 2017; Sapolsky et al., 2000). However, much less in known about the role CORT plays in influencing prolactin expression, signaling pathways, or synthesis. While prolactin is a critical component in the initiation and mediation of aspects of reproduction and parental care, it also plays a role in the immune response, as well as the inhibition of the HPA axis (Austin and Word, 2018; Cabrera-Reyes et al., 2017; Torner, 2016). Increases in circulating CORT following a short-term stressor can result in decreased circulating prolactin in breeding birds suggesting a prolactin-stress response (Angelier and Chastel, 2009). Functional tests of these genes and their effect on reproduction, parental care, and immune function during a stress response will help to further elucidate their role.

We also isolated sex-specific effects of elevated CORT in the pituitary during the stress response. We discovered that CORT drives the differential expression of 8 genes in males (Table 2) and 77 genes in females (Table 3). Only three genes, *KLP9, PVALB*, and *SLAINL*, differentially expressed uniquely in the male pituitary. *KLF9* and *PVALB* are associated with cell and endocrine signaling and both have known responses to stress or CORT (see Table 2) while the function of *SLAINL* is currently unknown. In comparison, we observed much more reactivity in female pituitary transcription (Table 3), with a general functional theme of their actions being related to myelination. Increased myelination and oligodendrogenesis has been reported in the hippocampus in response to exogenous glucocorticoids and immobilization stress, where it may modify the function of this tissue by decreasing neurogenesis and changing the hippocampal structure by increasing its white matter (Chetty et al., 2014). It’s unclear if a similar function of increasing oligodendrogenesis and decreasing neurogenesis occurs in the pituitary, perhaps in the pars nervosa, but currently the function of increased myelination in this tissue is unknown.

Another common functional theme that emerged in the pituitary included cell transport and signaling, which may be related to endocrine signaling, secretion, and feedback. Finally, we observed commonalities in gene function regarding the role of their products in immune responsiveness and as an antioxidant or mediator of oxidative stress. Transcription of genes associated with immune function or oxidative damage may, in the face of elevated CORT during a stress response, serve to modulate antioxidant activity (e.g. *APOA1, APOD, HEBP2*) or immune function (e.g., *CEBPD, CREB5, NINJ2, NT5E, TSC22D3*); however, immune actions of these differentially expressing genes in the pituitary is unclear with some acting to increase immune responsiveness (*CEBPD, NINJ2*) and others acting to inhibit it (e.g., *NT5E, TSC22D3*) (Table 3). We can only speculate at this time as to why the female pituitary is more responsive to the CORT signal during the acute stress response. In general, the HPG transcriptomic stress response is more pronounced in females as compared to males (Calisi et al., 2018). This may be because selection has favored increased stress responsiveness in females as compared to males. Females are, at least initially, more energetically invested in reproduction because their lay energy rich eggs and spend more time contact-incubating the clutch than males. A loss of a clutch or brood as a result of a stress-inducing perturbation would result in higher losses for females compared to males. As such, there is a clear selective advantage for females to be responsive to stressful conditions and respond by delaying the onset of breeding.

### Gonads

We identified the extent to which the gonadal transcriptomic response to 30 min of restraint stress could be explained by an increase in circulating CORT. In females, elevated CORT resulted in the differential expression of 28 genes during restraint stress (Table 4), while in males, no changes were observed. Commonalities in functionality of genes that altered their expression within the ovaries included cell signaling and transport, immune response, and cell growth and proliferation. Future research is needed to further uncover the function of these gene products within the ovaries in response to stress.

### Non-CORT mediated stress-responsive genes

Historically, CORT has been a large focus of mechanistic inquiry behind the study of stress-induced reproductive dysfunction. Because of this, research on the influential role of CORT on the HPG axis has predominated much of the stress and reproductive biology literature, potentially overshadowing other influential mediators. We discovered 1567 non-CORT mediated genes in male and female rock doves that differentially expressed in hypothalamic, pituitary, and gonadal tissues in response to restraint stress. Although increased circulation of CORT, and glucocorticoids in general, occur in response to a variety of stressors, their elevation should not be synonymous with “stress”, as they are involved in a variety of other functions and serve as only one component of complex physiological stress responses (MacDougall-Shackleton et al., 2019). Our findings support this notion.

Potential non-CORT drivers of observed changes in gene activity in response to stress include upstream activation of the sympathetic nervous system, the hippocampus, HPA, and HPT (hypothalamus-pituitary-thyroid) axes. An example of one gene of interest that is associated with limbic activation of the stress response is *CCK* (cholecystokinin). CCK is an enteroendocrine hormone associated with hunger and anxiety (Chen et al., 2010; Tang et al., 2012). In female rock doves exposed to restraint stress, *CCK* increases in the pituitary but does not respond to exogenous CORT treatment. Acute and chronic stress in the paraventricular nucleus (PVN) of the hypothalamus has been associated with increased expression of *CCK* (Tang et al., 2012), which may act as a neurotransmitter or neuromodulator in the brain (Hill et al., 1987) and interact with multiple neurotransmitter systems (e.g., dopaminergic, serotonergic, GABAergic; Tang et al., 2012). CCK may also interact with the HPA axis, including corticotropin-releasing factor (CRF), to modulate stress-related physiological responses and behaviors (Tang et al., 2012). Its predominance throughout the brain and limbic system suggests a role in the interaction between neurotransmitter systems, the HPA axis, and the limbic system during stress (Hill et al., 1987; Tang et al., 2012). Thus, the function of *CCK* as a neuromodulator and in the stress response could have wide-ranging physiological and endocrinological impacts.

There also exists the potential for external influence of the HPG axis from other bodily tissues and systems that are responsive to stress. During the stress response, the HPA axis receives signals from the central nervous system and the limbic system. These signals then interact with the HPA and HPT axes to promote individual survival, often at the cost of reproductive function (Romero and Wingfield, 2015). Theoretically, exogenous administration of CORT should bypass these earliest stages of the stress response (Astheimer et al., 1999; Claunch et al., 2017; Romero and Wingfield, 2015). Without input from the limbic system, upstream neuroendocrine and endocrine signaling does not activate the hypothalamic and pituitary stress response. Early CORT-independent responders to stress like catecholamines (e.g., epinephrine, norepinephrine, dopamine) are related to an increase in glucose metabolism, thereby making energy available to tissues during and after exposure to a stressor. These then have their own unique actions and interactions on or with tissues within the HPA, HPT, and HPG axes. Thus, the differential genomic response of the HPG axis to restraint stress as compared to exogenous CORT treatment could be indicative of a lack of input from the limbic system and HPA axis prior to the synthesis of CORT. In addition, CORT-independent endocrine cascades activated by the stress response can also act directly on the HPT axis, influencing its role in regulating metabolism (Helmreich and Tylee, 2011). In turn, metabolic rate can determine resource mobilization with the potential to inhibit the function of the reproductive system.

Finally, due to the dynamic and transient nature of gene transcription, translation, and physiological feedback mechanisms, we emphasize that the data we present represent an informative snapshot of gene activity 30 min post treatment exposure. It is likely that we would observe differential gene transcription at other time points. Future studies of this nature conducted along a temporal gradient will increase the resolution of our understanding of the dynamic genomic transcription and translation landscape.

### Conclusion

We report the causal and sex-typical effects of elevated CORT on the HPG stress response of the rock dove at the level of the transcriptome. We offer an extensive genomic and theoretical foundation on which to innovate the study of stress-induced reproductive dysfunction, offering novel gene targets to spur new lines of investigation and gene therapy development. Our results suggest that elevated circulating CORT concentrations are not responsible for the majority of transcriptional changes observed in the reproductive axis following exposure of 30 minutes of restraint stress. Studies investigating the role of glucocorticoids, like CORT, in the stress response predominate much of stress biology literature. For example, a PubMed search conducted on October 8, 2020 using the search terms “stress” with “corticosterone” or “cortisol” resulted in 13,771 hits and 21,424 hits, respectively. Searches for other stress-responsive genes we have identified in this and previous (Calisi et al., 2018) studies, such as “*KCNJ5”*, “*PTGER3*”, or “*CISH*”, resulted in 7, 12, and 5 hits, respectively, when associated with the search term “stress”. It may be time to shift emphasis from studying the role of this one hormone to investigating the roles of others in order to transcend our comprehension of stress and reproductive system interactions.

## MATERIALS AND METHODS

Subject housing, sampling and analysis procedures used in this study replicated those used in Calisi et al. (2018) unless otherwise noted. These methods are described in brief in the following text.

### Housing

Animals were socially housed at the University of California, Davis, in large semi-enclosed outdoor aviaries (1.5 ⨯1.2 ⨯2.1 meters). Each aviary had 16 nest boxes and approximately 8 sexually reproductive adult pairs and their dependent offspring. Food (Farmer’s Best Turkey/Game Bird Starter Crumbles (27% crude protein, 1.2% lysine, 0.3% methionine, 4% crude fat, 5% crude fiber, 0.8% phosphorous, 1-1.5% calcium, 0.3-0.6% NaCl, 0.25% Na; Farmers Warehouse Company, Keves, CA, USA), Farmer’s Best Re-cleaned Whole Corn (7.5% crude protein, 3.4% crude fat, 3.5% crude fiber, 1.7% ash; Farmers Warehouse Company, Keves, CA, USA), and red grit (33-35% calcium, 0.01% phosphorous, 0.05-0.08% Salt; Volkman Seed Factory, Ceres, CA, USA), water, and nesting material was provided *ad libitum*. Birds were exposed to natural light, which was augmented with artificial fluorescent lights set to a 14L:10D cycle. All birds collected in this study were between 5 months and 2 years old, sexually mature, and were not actively breeding at the time of collection.

### Corticosterone Solution

CORT (0.2mg/ml, Sigma 4-Pregnene-11beta-diol-3,20-dione, C-2505, Lot 092K1255) was dissolved by vortexing it in a peanut oil vehicle (Gam et al., 2011) one day prior to its administration and stored in a 15 mL centrifuge tube at room temperature. The concentration used to simulate circulating levels of endogenous CORT in response to restraint stress was informed by preliminary validation trials.

### Animal and Tissue Collections Methods

Collections occurred between 0900-1200 (PST) following animal care and handling protocols (UC Davis IACUC permit # 20618). Subjects were sexually mature and did not have an active nest. They were randomly assigned to either the vehicle-control group (hereafter, *control*, 8 females, 5 males received the peanut oil vehicle only) or CORT treatment group (8 females, 8 males received CORT mixed with peanut oil). To administer treatments, subjects were captured in ≤1 min of entering their aviary, transported by hand to an adjacent room, and immediately injected with either a control or CORT solutions intramuscularly (pectoralis muscle). Subjects were returned to their aviaries <5 min post initial disturbance and left undisturbed for 30 min, after which they were re-captured and immediately anesthetized using isoflurane (<2 min) prior to decapitation. Trunk blood was immediately collected after decapitation, brains were flash frozen on dry ice, and pituitaries and gonads were submerged in RNALater (Invitrogen, Thermo Fisher Scientific, REF: AM7021) at collection before freezing them on dry ice. All tissues were then transferred to a -80°C freezer and stored until analyses.

In the laboratory of Dr. Rebecca Calisi at the University of California, Davis, brains were sectioned coronally on a Leica CM 1860 cryostat at 100 µm to allow for precise biopsy of hypothalami and lateral septa, replicating our previous collections (Calisi et al., 2018; MacManes et al., 2017). Biopsied hypothalamic tissue was submerged in RNALater and shipped overnight on dry ice to the lab of Dr. Matthew MacManes at the University of New Hampshire for further processing. Trunk blood was centrifuged at 4°C for 10 minutes and plasma was aspirated and stored at -80°C.

### Hormone Assays

Plasma was assayed for circulating CORT concentrations using radioimmunoassay (RIA; see Calisi et al., 2018) using a dilution of 1:20 in a commercially available CORT RIA kit (MP Biomedicals, Orangeburg, NY) to confirm an increase in circulating CORT levels (ng/mL) in response to exogenous CORT and restraint stress treatments. The assay was validated for cross-reactivity to assess the potential for interference from other plasma compounds, and the limit of detection was estimated at 0.0385 ng/mL.

Two-way ANOVAs were conducted to assess differential circulating concentrations between groups (α = 0.05; lme4 v1.1-17, R v3.5.1), and *post-hoc* pairwise comparisons were conducted with a Dunnett adjustment for multiple comparisons (emmeans v.1.2.3). Prior to analysis, CORT values were ln-transformed to normalize the distribution of the data in order to meet model assumptions. Back-transformed estimates (e^estimate^) are presented in the results, which should be interpreted as a magnitude difference between groups.

### Illumina Library Preparation and Sequencing

Tissues frozen in RNALater were thawed on ice in an RNAse-free work environment. Total RNA was extracted using a standard Trizol extraction protocol (Thermo Fisher Scientific, Waltham, MA), and RNA quality was assessed using the Tapestation 2200 Instrument (Agilent, Santa Clara, CA). Illumina sequence libraries were prepared using the TruSeq RNA Stranded LT Kit (Illumina, San Diego, CA), and library quality assessed using the Tapestation 2200 Instrument (Agilent, Santa Clara, CA). Each library was diluted to 2nM with sterile purified commercially available molecular biology-grade water (VWR) and pooled in a multiplexed library sample. The multiplexed library sample was then sent to the Novogene company for 125 base pair paired-end sequencing on a HiSeq 4000 platform.

### Transcriptome assembly evaluation and improvement

The previously constructed Rock Dove transcriptome version 1.0.4 assembly (Calisi et al., 2018) was evaluated to ensure that transcripts expressed uniquely in the stress condition were included. To accomplish this, reads from the pituitary, hypothalamus, and gonads from one stressed male and one stressed female were assembled following the Oyster River Protocol (MacManes, 2018). Unique transcripts contained in this assembly relative to the previously described assembly were identified via a BLAST procedure. Novel transcripts, presumably expressed uniquely in the stress condition were added to this existing assembly, thereby creating the Rock Dove v. 1.1.1 transcriptome (available at https://s3.amazonaws.com/reference_assemblies/Rockdove/transcriptome/RockDove.HPG.v1.1.1.fasta). This new assembly was evaluated for genic content via comparison with the BUSCO version 2.0 Aves database (Simão et al., 2015).

### Mapping and Global Analysis of Differential Gene Expression

Reads were quality and adapter trimmed to a Phred score =2 using the software package Trimmomatic (Bolger et al., 2014). Reads were then quasimapped to the Rock Dove transcriptome (v. 1.1.1) after an index was prepared using Salmon 0.9.0 (Patro et al., 2017). Rock dove transcript IDs were mapped to genes from the *Gallus gallus* genome (v. 5), using BLAST (Camacho et al., 2009). All data were then imported into the R statistical package (v. 3.3.0) (RStudio Team, 2015) using tximport (Soneson et al., 2016) for gene level evaluation of gene expression, which was calculated using contrasts that separately compared treatment group (control vs. CORT treatment) for each tissue (hypothalamus, pituitary, gonads) and sex (male or female) using edgeR (v. 3.5.0) (Robinson et al., 2010) following TMM normalization and correction for multiple hypothesis tests by setting the false discovery rate (FDR) to 1%. Differential expression values resulting from 30 min of restraint stress were taken directly from Calisi et al. (2018). Briefly, contrasts comparing treatment groups (baseline control vs. restraint stress treatment) were calculated as above for each tissue and sex (see (Calisi et al., 2018) for more details).

### Candidate Gene Expression Evaluation

A set of 44 genes that target specific research questions was selected *a priori* for evaluation based on their known involvement in reproduction and in association with the stress response or CORT feedback (Table 5). Normalized values of gene expression were compared by treatment group. Differences in gene expression of candidate genes in the hypothalamic, pituitary, and gonadal tissues were compared in response to CORT treatment as compared to controls (expression ∼ treatment) or involved a re-analysis of restraint-stress and baseline controls from Calisi et al. (2018) using a robust regression model framework (alpha=0.05; rlm, Package MASS v7.3-50; robtest, Package sfsmisc v1.1-2; R v3.5.2; Venables et al., 2002). These analyses were conducted separately for each sex.

In Calisi et al. (2018), a similar candidate gene analysis was performed using generalized linear models; however, due to the high variance of the *y*-variable (i.e., normalized gene expression) for a number of candidate genes, and the overall smaller sample size in the current study, we deemed robust regression a more appropriate analytical approach. Briefly, robust regression is a type of analysis that both incorporates and reduces the influence of outliers on results without discounting their contributions. This approach is a more conservative approach, and thus, can increase the chance of false negatives, particularly when effect sizes are small. To make the previous work on restraint-stress directly comparable with the results from current analyses, and to remain internally consistent within the current study, we chose to re-analyze gene expression of the target genes from the restraint-stress dataset using the methodology we deemed most appropriate for the current study. We also expanded on the original candidate gene list from Calisi et al. (2018) to include a number of genes associated with CORT activation or de-activation, which were of particular interest in the current study. Because we used a different, more conservative analytical to analyze the CORT dataset that then required us to re-analyze the restraint stress dataset, there were some differences in statistical significance in the current study than were originally reported in Calisi et al. (2018). Any data we report that differ between those reported in Calisi et al. (2018) and this re-analysis can be attributed to the use of a different analytical approach.

### Isolating the role of CORT in the HPG genomic stress response

We compared genes that differentially expressed in response to CORT treatment (as compared to vehicle-injected controls) to those that differentially expressed in response to restraint-stress treatment (as compared to non-stressed controls; the latter results we reported in Calisi et al. (2018). Genes identified as differentially expressed in response to both experimental manipulations (CORT and restraint-stress treatments) as compared to their respective controls (vehicle control and unstressed treatments) were considered the genes within the HPG axis whose response to the stress stimuli was most likely due to elevated circulating concentrations of CORT.

## Supporting information

Supporting Information: Exogenous CORT

Supporting Information: Restraint Stress

## ACKNOWLEDGEMENTS

We thank the many undergraduate researchers of the Calisi Lab, especially Olivia Calisi, Tiffany Chen, and Tanner Feustel for their assistance on this project, including their involvement in avian husbandry, conducting the experiment, and tissue collection. We particularly thank Luke Remage Healey for his input during early stages of this work, the labs of Matthew MacManes and Titus Brown for computational support, as well as Paulina Gonzalez-Gomez, Tom Hahn, Isaac Ligocki, Gabrielle Names, Jonathan Perez, Marilyn Ramenofsky, John Wingfield, and Karen Word for intellectual contributions and discussions. This work was funded by NSF IOS 1455960 (to RMC and MM).

## Author contributions

RMC conceived the idea and developed the experimental design. JSK and MM provided additional logistical advice. AMB validated the CORT dosage. VSF and AMB conducted the data collection, SHA and AMB processed the tissues and biopsied hypothalami, and ASL completed the RNA extraction and library preparation prior to sequencing. ASL, SHA, TAH, RMH, and MM conducted data analyses. SHA drafted the manuscript, RMC edited the manuscript, and all authors contributed input to the document.

