## Supporting Information: Exogenous CORT for "Isolating the role of corticosterone in the hypothalamic-pituitary-gonadal genomic stress response"

**Supplemental Information 1: CORT**

CORT specific, differentially expressed genes by tissue and sex. Each table contains differentially expressed genes that were unique to CORT treatment, which are either sex-specific or overlapping across sex. We report log fold change (logFC) and false discovery rate (FDR<0.01). Upregulated genes are highlighted in red and downregulated genes are blue.

**Table 1:** Sex-specific, CORT responsive genes that were differentially expressed in the female hypothalamus

| Entrez ID | Gene Name | logFC | FDR |
|---|---|---|---|
| 395384 | *C4BPA* | 7.62191024 | 0.00081864 |
| 107053650 | *DDIT4* | -1.3641926 | 0.00000722 |
| 100858474 | *LOC100858474* | 6.98742125 | 0.00846125 |

**Table 2:** Sex-specific, CORT responsive genes that were differentially expressed in the male hypothalamus.

| Entrez ID | Gene Name | logFC | FDR |
|---|---|---|---|
| 420592 | *LRRC72* | -6.3325476 | 0.00273415 |

**Table 3:** Sex-specific, CORT responsive genes that were differentially expressed in the female pituitary.

| Entrez ID | Gene Name | logFC | FDR |
|---|---|---|---|
| 424490 | *ABCA4* | 3.42839287 | 0.00087459 |
| 769237 | *ABI3BP* | -1.8203605 | 0.00700656 |
| 431627 | *ACOT12* | -2.5289569 | 0.00792938 |
| 418479 | *ADAMTS1* | -1.350909 | 0.00024624 |
| 421297 | *ALK* | 3.65729005 | 0.00000214 |
| 420738 | *ANLN* | -4.8887466 | 0.00000171 |
| 396536 | *APOA1* | -2.287156 | 0.00628945 |
| 424893 | *APOD* | -3.5765603 | 0.00122136 |
| 769889 | *APOLD1* | -1.9641033 | 0.0000477 |
| 421088 | *AQP4* | -2.8884783 | 0.00023196 |
| 421369 | *ATF3* | -1.8172924 | 0.00025 |
| 107052744 | *C2H8orf34* | -2.9389569 | 0.00889358 |
| 422966 | *C5H11ORF9* | -4.2336243 | 0.0000629 |
| 770209 | *C5H11ORF96* | -1.1061049 | 0.00774226 |
| 423959 | *C6H10ORF90* | -2.1371616 | 0.00803811 |
| 421029 | *CDH19* | -3.9098923 | 0.0000448 |
| 395921 | *CNP* | -3.1456022 | 0.0000169 |
| 427597 | *CNTN6* | -3.7792797 | 0.00046519 |
| 428435 | *CREB5* | -1.9740611 | 0.00707024 |
| 421422 | *DAAM2* | -1.4990997 | 0.00768802 |
| 776499 | *DCLK3* | -1.1192708 | 0.00022132 |
| 107053650 | *DDIT4* | -1.4852373 | 1.21E-08 |
| 395939 | *DIO3* | 4.16775226 | 0.00000557 |
| 425241 | *DIRAS2* | -1.5768922 | 0.00017086 |
| 421884 | *DST* | -4.3504175 | 0.00023196 |
| 422551 | *ENPP6* | -2.3741719 | 0.00202937 |
| 771826 | *ERMN* | -5.2206005 | 0.0000169 |
| 426690 | *ETNK2* | -2.6377886 | 0.00078273 |
| 415687 | *FA2H* | -4.6788107 | 0.0000133 |
| 422413 | *FAM198B* | -3.1443697 | 0.00300551 |
| 419084 | *GDPD4* | -2.2916137 | 0.00042517 |
| 419969 | *GFAP* | -5.9757928 | 0.0000837 |
| 420397 | *GJC2* | -1.6502815 | 0.00442165 |
| 396489 | *GLUL* | -1.869448 | 0.00110485 |
| 431418 | *GPR139* | 0.97276845 | 0.00136411 |
| 417748 | *GPR37* | -3.3203086 | 0.00227741 |
| 421176 | *GPR37L1* | -3.9247638 | 0.00129104 |
| 427575 | *GRM2* | -2.3590185 | 0.00030702 |
| 416728 | *HAO1* | -1.2621179 | 0.00867023 |
| 425975 | *HAPLN2* | -5.4044026 | 0.00301302 |
| 424441 | *HEBP2* | -2.5446672 | 0.00195309 |
| 428234 | *HEPACAM* | -3.8809357 | 0.0000693 |
| 395128 | *HES4* | 1.43850911 | 0.00000368 |

| | | | |
|---|---|---|---|
| 426516 | *HNRPK* | 3.26377291 | 0.0042366 |
| 419771 | *IL10RA* | 1.63727996 | 0.0000163 |
| 427662 | *KCNJ12* | -3.0905596 | 0.00525551 |
| 422374 | *KIAA1210* | -1.2923415 | 0.00233732 |
| 420476 | *KIAA1462* | -1.0813841 | 0.0082338 |
| 421934 | *KLF11* | 1.05197595 | 0.00745418 |
| 416732 | *LAMP5* | -1.3715911 | 0.00026828 |
| 107049625 | *LGI3* | -4.348784 | 0.0000167 |
| 418889 | *LHFP* | -1.3575629 | 0.00026828 |
| 395705 | *LHX2* | -2.9103875 | 0.00129104 |
| 100859224 | *LOC100859224* | -5.8575025 | 0.0000693 |
| 100859848 | *LOC100859848* | -2.1970407 | 0.00032798 |
| 107050516 | *LOC107050516* | -3.6208916 | 0.00000788 |
| 107055436 | *LOC107055436* | 2.18207245 | 0.00834063 |
| 416927 | *LOC416927* | -5.5556961 | 0.0000943 |
| 769039 | *LOC769039* | 1.01210942 | 0.00141439 |
| 374011 | *MAB21L2* | 1.82144289 | 0.0088409 |
| 396217 | *MBP* | -1.8132315 | 0.00176756 |
| 107055352 | *MIDN* | 0.94959316 | 0.00015121 |
| 417737 | *MLC1* | -2.1486007 | 0.00091839 |
| 419010 | *NAALAD2* | -2.0113394 | 0.00424508 |
| 418151 | *NINJ2* | -5.1322959 | 0.00000115 |
| 421833 | *NT5E* | -4.373281 | 0.00019306 |
| 395334 | *OPN4-1* | -4.4283406 | 0.00296667 |
| 422007 | *OTOF* | -3.8480519 | 0.0000429 |
| 422030 | *OvoDA1* | 5.0744276 | 0.00700656 |
| 100858468 | *PDYN* | 2.40007783 | 0.0000954 |
| 771808 | *PIPOX* | -3.5980662 | 0.0018916 |
| 415650 | *PLLP* | -4.5541969 | 0.00000536 |
| 396214 | *PLP1* | -8.9306286 | 3.52E-07 |
| 425328 | *PPP1R3C* | -1.5194554 | 0.0000429 |
| 374187 | *PTCH2* | 1.0457712 | 0.00792938 |
| 416507 | *RASD1* | -1.4417544 | 0.00023196 |
| 422756 | *RASL11B* | -1.0400228 | 0.00708453 |
| 419521 | *RASSF2* | -3.0547377 | 0.0001018 |
| 424409 | *RGS16* | 2.27790308 | 0.00792938 |
| 424409 | *RGS16* | 2.27790308 | 0.00792938 |
| 395359 | *SALL3* | 0.78394827 | 0.00155515 |
| 374006 | *SAT1* | -1.4298953 | 0.00000125 |
| 418228 | *SCUBE1* | -1.0781472 | 0.00700656 |
| 424593 | *SEPP1L* | -4.1167333 | 0.0000586 |
| 416778 | *SEPT5* | -1.8086218 | 0.00104754 |
| 107049626 | *SFTPC* | -4.1712098 | 0.00792938 |
| 418731 | *SH3RF3* | -2.1217571 | 0.0018969 |
| 396089 | *SHANK3* | -5.6229119 | 0.00022132 |
| 395615 | *SHH* | 1.95114021 | 0.00950115 |
| 427459 | *SLC28A3* | 1.87914802 | 0.00296098 |
| 424576 | *SLC6A9* | -2.9307868 | 0.0000629 |

| | | | |
|---|---|---|---|
| 432368 | *SNAI2* | 1.47526135 | 0.00227741 |
| 395573 | *SOX10* | -4.4550521 | 0.00059949 |
| 395483 | *SOX8* | -2.3092929 | 0.0000501 |
| 416208 | *STC2* | -2.2605966 | 0.00576626 |
| 395592 | *TH* | 6.15460068 | 0.00000115 |
| 419401 | *TMEM52* | -3.2746687 | 0.000018 |
| 419414 | *TMEM88B* | -5.0726254 | 0.00016927 |
| 396032 | *TNNC1* | -4.7851431 | 0.0000288 |
| 768091 | *TSC22D3* | -1.0163052 | 0.00122136 |
| 428900 | *TSHR* | -3.1297683 | 0.00828514 |
| 419889 | *TSPAN2* | -1.4330295 | 0.00016927 |
| 100858879 | *TTYH2* | -1.7900637 | 0.004794 |
| 107051021 | *TXNIP* | -1.0489175 | 0.00304851 |
| 374033 | *UGT8* | -2.672702 | 0.00035335 |
| 428995 | *VWC2L* | -0.8910228 | 0.00792938 |
| 416887 | *WSCD2* | 1.08909738 | 0.00202714 |
| 419618 | *ZC3H12A* | -0.7386901 | 0.00587455 |

**Table 4:** Sex-specific, CORT responsive genes that were differentially expressed in the male pituitary.

| Entrez ID | Gene Name | logFC | FDR |
|---|---|---|---|
| 423846 | *ARHGAP19* | 1.78534547 | 0.00987046 |
| 101750689 | *BHLHE41* | 0.98090895 | 0.00439612 |
| 771055 | *C1H2ORF40* | 2.24634841 | 0.0004201 |
| 770534 | *C5H15ORF62* | -1.8005857 | 0.00022038 |
| 396256 | *CALCA* | 4.21679034 | 0.00413295 |
| 416155 | *CCNJL* | -2.0430218 | 0.00509869 |
| 395527 | *CD9* | 2.66785803 | 0.0004201 |
| 395386 | *CLEC3B* | 1.33483153 | 0.00509869 |
| 428448 | *COL6A6* | 2.16089158 | 0.000035 |
| 771797 | *FAM101A* | 1.59623628 | 0.00060137 |
| 396119 | *FST* | 3.90119714 | 0.0000487 |
| 428628 | *GRIK2* | -2.1045585 | 0.00032328 |
| 419110 | *ITIH2* | 2.72026805 | 0.00048391 |
| 428101 | *KCNA1* | 2.27275094 | 0.00413295 |
| 425504 | *KCTD12* | 1.07818415 | 0.00054115 |
| 770238 | *KLF9* | -1.1719745 | 0.00413295 |
| 100858447 | *LOC100858447* | 5.72337106 | 0.00251315 |
| 427408 | *LOC101747901* | 2.71309605 | 0.0004201 |
| 395944 | *LOC395944* | 7.99361432 | 0.00066177 |
| 404752 | *MID1IP1* | 0.83233673 | 0.00255214 |
| 421889 | *MLIP* | 6.35614627 | 0.0000263 |
| 769245 | *MZT2B* | 5.3058003 | 0.00240232 |
| 422078 | *NPTX1* | 2.1128186 | 0.00509869 |
| 693264 | *NPY6R* | -7.0988646 | 0.0000263 |
| 424117 | *PPP1R1C* | 2.00472083 | 0.00550383 |
| 417183 | *PRDM12* | 3.76858544 | 0.00107929 |
| 424548 | *PTGFR* | -1.814985 | 0.00753864 |
| 396459 | *PVALB* | 2.9174758 | 0.00026039 |
| 424312 | *RND3* | 1.72048443 | 0.0000263 |
| 420251 | *RNF19A* | -0.9587109 | 0.00930352 |
| 107053895 | *RPRM* | 3.06692678 | 0.00049207 |
| 421715 | *RSPO3* | 2.70938016 | 0.0000867 |
| 395715 | *SERPINB10* | 2.74747729 | 0.00550383 |
| 417616 | *SLC47A2* | 5.32725411 | 0.00119519 |
| 395393 | *SNCA* | 2.53594265 | 0.00413295 |
| 107051027 | *SOX21* | -4.0598474 | 0.00240232 |
| 420073 | *SSPO* | -0.8494481 | 0.00409517 |
| 420573 | *TAC1* | 3.19991435 | 0.00011946 |
| 395810 | *TEAD3* | -4.3106756 | 0.00944429 |
| 427825 | *TLCD2* | 2.39842736 | 0.0046092 |
| 417905 | *TMCC3* | -1.3145099 | 0.00155544 |
| 423491 | *TMEM179* | 1.42215295 | 0.00087152 |
| 423477 | *TNFAIP2* | 1.55703179 | 0.00823426 |

**Table 5:** Sex-specific, CORT responsive genes that were differentially expressed in the female gonads

| Entrez ID | Gene Name | logFC | FDR |
|---|---|---|---|
| 418254 | *A2ML1* | 3.89950284 | 0.0000171 |
| 373945 | *ABCA1* | -0.8640436 | 0.00774306 |
| 776390 | *ACPP* | 1.82354474 | 0.00339185 |
| 100858277 | *ADAMTS18* | 1.37171569 | 0.00291717 |
| 426545 | *ADPRHL* | 1.9432785 | 0.00888002 |
| 771077 | *AKD1* | -2.3964956 | 0.0000966 |
| 415402 | *AQP9* | -1.2561301 | 0.00011139 |
| 769191 | *ASIC4* | 2.1852466 | 0.00124521 |
| 424349 | *ATP6V1G3* | 1.56758017 | 0.00396435 |
| 421809 | *BACH2* | -1.0371513 | 0.00393492 |
| 420744 | *BMPER* | -1.5135845 | 0.00593651 |
| 770363 | *BSG* | 0.95723565 | 0.0000966 |
| 417179 | *C9ORF58* | 1.37426866 | 0.00139229 |
| 417647 | *CA4* | -3.0968458 | 3.61E-07 |
| 421139 | *CA8* | 3.27931835 | 5.16E-07 |
| 417151 | *CACFD1* | 0.90162358 | 0.00367487 |
| 415996 | *CACNA2D3* | -1.7080744 | 0.00174309 |
| 396519 | *CALB1* | 2.43168658 | 0.0000216 |
| 423990 | *CALCRL* | -0.6870457 | 0.00396435 |
| 427515 | *CALML4* | -1.8826522 | 0.00906168 |
| 424601 | *CCDC17* | -3.0541205 | 0.00462991 |
| 415315 | *CD276* | 1.71955626 | 1.57E-07 |
| 107052707 | *CEBPD* | -1.4845859 | 0.0000216 |
| 415476 | *CEMIP* | -2.1134273 | 0.00042534 |
| 415507 | *CHD2* | -2.7333219 | 0.00210892 |
| 396120 | *CHRNA8* | -3.053195 | 0.00744057 |
| 428866 | *CHST1* | 1.63775732 | 0.00065183 |
| 395399 | *CITED2* | -1.1996508 | 0.0000211 |
| 374002 | *CKMT1A* | 2.61658061 | 4.88E-08 |
| 424768 | *CLDN15* | -1.7043716 | 0.00356272 |
| 100858979 | *COL10A1* | 6.78061851 | 2.49E-07 |
| 396503 | *COL17A1* | 1.74199079 | 0.0000404 |
| 807639 | *COX1* | -1.4285244 | 0.00396435 |
| 807635 | *COX2* | -1.6448941 | 0.0016697 |
| 807637 | *COX3* | -1.3992481 | 0.00480154 |
| 101747832 | *CPNE9* | 3.47709786 | 0.00063875 |
| 768680 | *CREG1* | -0.9083675 | 0.00829127 |
| 417735 | *CRELD2* | 0.90843884 | 0.00774306 |
| 396499 | *CRYBA1* | 2.91055692 | 0.0054706 |
| 408252 | *CTDSPL* | -0.6689948 | 0.00727034 |
| 424152 | *CYBRD1* | -1.1392711 | 0.00210892 |
| 427802 | *CYGB* | -1.5605939 | 0.00210892 |

| | | | |
|---|---|---|---|
| 807641 | *CYTB* | -1.3743486 | 0.00396435 |
| 107053650 | *DDIT4* | -0.830587 | 0.00953958 |
| 425542 | *DDOST* | 0.71084418 | 0.00888002 |
| 770448 | *DESI1* | 0.65934636 | 0.0003117 |
| 395271 | *DMBX1* | -2.7365372 | 0.00017366 |
| 427004 | *DNAH3* | -2.3586282 | 0.0011016 |
| 420571 | *DYNC1I1* | 1.49516771 | 0.00367487 |
| 395689 | *ENO2* | -1.8681352 | 0.00049186 |
| 421703 | *EPB41L2* | -1.8790429 | 0.00424831 |
| 421703 | *EPB41L2* | -0.8653211 | 0.00926777 |
| 421509 | *ERO1LB* | 0.79927548 | 0.0002771 |
| 769188 | *FAM161A* | -1.5780313 | 0.00090247 |
| 415505 | *FAM174B* | 0.81170839 | 0.0002771 |
| 771833 | *FAM194A* | -2.432687 | 0.00657155 |
| 424427 | *FAM20B* | 0.69560334 | 0.00396435 |
| 416889 | *FICD* | 1.00312345 | 0.00093458 |
| 428618 | *FLRT1* | 3.93716024 | 0.00893298 |
| 421365 | *FLVCR1* | 1.08548101 | 0.00013067 |
| 386747 | *FMN1* | -1.2951485 | 0.00367487 |
| 771133 | *FUT9* | -3.6679303 | 0.00020872 |
| 419770 | *FXYD2* | 3.18708386 | 0.00016312 |
| 424171 | *G6PC2* | -1.7674145 | 0.00037283 |
| 417006 | *GAL3ST1* | 1.70145792 | 0.00953958 |
| 429501 | *GALNT6* | 0.8560541 | 0.00280289 |
| 378781 | *GH* | -2.84771 | 0.0000139 |
| 428397 | *GHRH-LR* | 3.97213572 | 8.06E-07 |
| 396533 | *GOT2* | 0.6756896 | 0.0016697 |
| 431418 | *GPR139* | -1.484179 | 0.00103737 |
| 100857454 | *GPX2* | 2.80083524 | 0.00037283 |
| 422407 | *GUCY1A3* | 1.3132422 | 0.0000634 |
| 107055138 | *HAX1* | -1.8224772 | 0.0000715 |
| 395145 | *HMGCR* | 0.73169209 | 0.00154313 |
| 423079 | *HPS5* | -2.7970592 | 0.00010663 |
| 100858361 | *HSD11B2L* | 1.85096469 | 0.00010447 |
| 395785 | *HSD17B4* | 0.68506514 | 0.00461263 |
| 428251 | *HYOU1* | 0.82269448 | 0.00454652 |
| 396362 | *IAPP* | -3.7874244 | 0.00367487 |
| 422219 | *IL13RA2* | 2.08037281 | 0.00033948 |
| 107053356 | *INAFM2* | 1.15517408 | 0.00505892 |
| 419347 | *KCNG1* | 1.30521608 | 0.00027795 |
| 427917 | *KIAA1147* | 0.64824719 | 0.00893298 |
| 418645 | *KIAA1324L* | 1.83865557 | 7.64E-08 |
| 427588 | *KLF15* | -0.9539071 | 0.00396435 |
| 419829 | *KLHDC8A* | 1.18600396 | 0.00057293 |
| 424851 | *KLHL30* | 0.90227776 | 0.00676046 |
| 395772 | *KRT7* | 1.04916276 | 0.00437386 |
| 418840 | *LACC1* | -1.5526333 | 0.00462991 |
| 426779 | *LETM2* | 0.78147233 | 0.00896754 |

| | | | |
|---|---|---|---|
| 425107 | *LGALS2* | 2.37391673 | 0.00296288 |
| 423789 | *LIPA* | -0.977248 | 0.00296288 |
| 426849 | *LMAN1* | 0.74421571 | 0.00918287 |
| 100857341 | *LOC100857341* | -2.5874059 | 0.00359958 |
| 100859804 | *LOC100859804* | -2.5562857 | 0.00063875 |
| 101749151 | *LOC101749151* | 2.93828317 | 0.00016312 |
| 101750367 | *LOC101750367* | 2.32214996 | 0.00220046 |
| 107049904 | *LOC107049904* | 7.36511003 | 0.000045 |
| 107050309 | *LOC107050309* | -3.9016967 | 0.00628612 |
| 107050569 | *LOC107050569* | 2.15686607 | 0.00420313 |
| 107050717 | *LOC107050717* | 1.05400409 | 0.0000216 |
| 107050736 | *LOC107050736* | 1.25268381 | 0.0003117 |
| 107053409 | *LOC107053409* | -7.0048116 | 0.00000801 |
| 107056154 | *LOC107056154* | -1.2099324 | 0.00351022 |
| 417800 | *LOC417800* | 2.46484862 | 0.00462991 |
| 418356 | *LOC418356* | 3.76747091 | 0.00628612 |
| 422179 | *LOC422179* | -1.5387874 | 0.0019543 |
| 422895 | *LOC422895* | 1.98337799 | 1.38E-15 |
| 428451 | *LOC428451* | 2.71377612 | 0.00424831 |
| 428971 | *LOC428971* | 2.10673556 | 0.00011139 |
| 429955 | *LOC429955* | 1.44884673 | 0.00063875 |
| 419533 | *LOXL2* | 3.66285514 | 7.7E-12 |
| 424469 | *LRRC39* | -2.333139 | 0.00381132 |
| 101749866 | *LRTM2* | 2.50705725 | 0.0000715 |
| 693248 | *MAF* | -1.0821154 | 0.00687367 |
| 419173 | *MAFB* | -1.2095523 | 0.00291717 |
| 421047 | *MC5R* | -1.1187812 | 0.0016697 |
| 420437 | *MLL3* | -0.6682672 | 0.00436886 |
| 420437 | *MLL3* | -0.6682672 | 0.00436886 |
| 395683 | *MMP13* | -3.7433912 | 0.0031882 |
| 395387 | *MMP9* | -2.5485055 | 5.23E-08 |
| 423101 | *MUC2* | -1.7108933 | 0.00746836 |
| 414878 | *MUC6* | -5.8831043 | 0.00124294 |
| 421948 | *MYCN* | 1.29181577 | 0.00094369 |
| 417598 | *NCOR1* | -3.6541352 | 0.00888002 |
| 807636 | *ND1* | -1.6545808 | 0.0006331 |
| 430210 | *NDUFV1* | 0.83758957 | 0.00424831 |
| 395736 | *NEDD9* | -1.2588805 | 3.61E-07 |
| 396093 | *NFKBIA* | -0.7241225 | 0.00953958 |
| 419697 | *NIPAL3* | 0.63982001 | 0.00557854 |
| 422748 | *NMU* | 3.59947575 | 0.00468661 |
| 422535 | *NPNT* | -1.0780087 | 0.0002771 |
| 396464 | *NPY* | 2.38207228 | 0.00528824 |
| 421746 | *NT5DC1* | 1.24861588 | 0.00005 |
| 427914 | *NT5DC3* | -1.3970822 | 0.00763554 |
| 428789 | *NWD2* | 2.33924957 | 0.00070335 |
| 422525 | *OSTC* | 0.78243341 | 0.00213222 |
| 374091 | *P4HB* | 0.95277875 | 0.0000216 |

| | | | |
|---|---|---|---|
| 428065 | *PCDH8* | 1.72058962 | 0.00154313 |
| 107050625 | *PCOLCE* | 0.98943619 | 0.00291717 |
| 429078 | *PDC* | 3.17098684 | 0.00789292 |
| 421940 | *PDIA6* | 0.76593851 | 0.00469647 |
| 374241 | *PI15* | 3.22012014 | 0.00000311 |
| 423705 | *PLA2G12B* | 1.90953988 | 2.65E-07 |
| 418183 | *PLEKHA5* | -1.5609971 | 0.0000095 |
| 428164 | *PLTP* | -1.3820884 | 0.00057293 |
| 417989 | *PMM1* | 0.6991518 | 0.00896754 |
| 422019 | *PNOC* | 3.87524835 | 0.00000407 |
| 427605 | *PSD2* | 1.12349848 | 0.00172704 |
| 429116 | *PTGER3* | 1.26713061 | 0.00415405 |
| 771985 | *PTPRR* | 1.82197161 | 0.0000238 |
| 373914 | *QSOX1* | 1.21146443 | 0.00608505 |
| 430750 | *RAB31* | -1.0216729 | 0.00589831 |
| 420203 | *RALYL* | -2.3610435 | 0.00169396 |
| 395209 | *RARRES1* | -1.0426794 | 0.00426344 |
| 416507 | *RASD1* | 1.92966589 | 2.29E-07 |
| 396107 | *RET* | 1.27881441 | 0.00229095 |
| 374105 | *RIMBP2* | 2.02082994 | 0.00005 |
| 770339 | *RP11-26J3.4* | 5.52485415 | 0.00000033 |
| 396090 | *RP11-295K3.1* | -1.2580618 | 0.00022682 |
| 373919 | *RUNX2* | -2.1708331 | 0.00628612 |
| 419434 | *SAMD11* | 1.04337564 | 0.00033948 |
| 426369 | *SEC61A2* | 0.68376549 | 0.00571702 |
| 420895 | *SERPINB6* | 1.23660884 | 0.0080927 |
| 418651 | *SHROOM2* | -0.9145476 | 0.00896754 |
| 769612 | *SLC1A4* | -1.3967103 | 0.00016466 |
| 427845 | *SLC26A4* | 2.10730019 | 0.0000966 |
| 425029 | *SLC33A1* | 1.15865888 | 0.0003178 |
| 395185 | *SLC35B1* | 1.1279135 | 0.0000142 |
| 107052162 | *SLC38A7* | -1.3452675 | 0.00063875 |
| 107052162 | *SLC38A7* | -0.8653211 | 0.00926777 |
| 430411 | *SLC39A8* | 2.15744714 | 0.00665573 |
| 100861584 | *SLC6A8* | 1.2343538 | 3.61E-07 |
| 418719 | *SLC9A2* | 2.75708953 | 0.00013067 |
| 416550 | *SLC9A3R2* | 0.86671721 | 0.00356272 |
| 424862 | *SLCO2A1* | -1.6100793 | 0.0016697 |
| 419051 | *SLCO2B1* | 1.54035864 | 0.00763554 |
| 396105 | *SOX2* | -1.5319176 | 0.0004654 |
| 429033 | *SPEG* | 0.96717445 | 0.00367487 |
| 423206 | *SPINT1* | 1.48255557 | 3.61E-07 |
| 423225 | *SPTBN5* | -2.3843542 | 0.00063875 |
| 420871 | *SSR1* | 0.92584722 | 0.00030165 |
| 425656 | *SSR2* | 0.80422596 | 0.00293201 |
| 425027 | *SSR3* | 0.72481355 | 0.00405625 |
| 107049822 | *SSR4* | 0.91298825 | 0.0013767 |
| 771648 | *SSTR1* | 1.81140495 | 0.00576975 |

| | | | |
|---|---|---|---|
| 396241 | *TF* | 3.19249305 | 0.00174309 |
| 417575 | *TLCD1* | 1.08308207 | 0.0013767 |
| 416821 | *TMED2* | 0.70931446 | 0.00292529 |
| 424483 | *TMEM56* | 1.08237378 | 0.00313371 |
| 395883 | *TMOD1* | -1.9474436 | 0.00066919 |
| 415724 | *TOX3* | -1.0783869 | 0.00426344 |
| 424835 | *TRPM2* | 2.51575982 | 0.0000216 |
| 768091 | *TSC22D3* | -1.043332 | 0.00064229 |
| 426419 | *UNC5A* | 1.64916428 | 0.0000216 |
| 395582 | *VAX1* | -3.6431957 | 0.00699024 |
| 421702 | *VNN1* | -2.423167 | 0.00020872 |
| 421702 | *VNN1* | -1.5921092 | 0.0003178 |
| 414795 | *VSIG1* | 3.06906773 | 0.00396435 |
| 431270 | *VWA5A* | 1.97692897 | 0.00687367 |
| 416918 | *XBP1* | 1.01818103 | 0.00057071 |
| 771005 | *XKRX* | 1.11122284 | 0.0013767 |
| 419759 | *ZBTB16* | -1.1932609 | 0.00336961 |
| 421763 | *ZBTB24* | -1.1212923 | 0.00547041 |
| 107052930 | *ZFP36L2* | -0.9092372 | 0.0000139 |
| 419152 | *ZNF341* | -1.2979584 | 0.00321418 |
| 422017 | *ZNF395* | -1.0349004 | 0.00279947 |

**Table 6:** Sex-specific, CORT responsive genes that were differentially expressed in the male gonads

| Entrez ID | Gene Name | logFC | FDR |
|---|---|---|---|
| 419751 | GRIK4 | 1.12000808 | 0.00145882 |

**Table 7:** Differentially expressed genes found in the hypothalamus of both sexes for CORT.

| Entrez ID | Gene Name | Female logFC | Female FDR | Male logFC | Male FDR |
|---|---|---|---|---|---|
| 420570 | *PDK4* | -1.762850788 | 0.0023829 | -2.688725 | 0.00000577 |
| 378912 | *RGS2* | -1.584808442 | 0.0000891 | -1.9444537 | 0.0000232 |

**Table 8:** Differentially expressed genes found in the pituitary of both sexes for CORT.

| Entrez ID | Gene Name | Female logFC | Female FDR | Male logFC | Male FDR |
|---|---|---|---|---|---|
| 419316 | *APCDD1L* | -2.786556117 | 0.000000219 | 1.758956886 | 0.009303519 |
| 417431 | *APOH* | -5.673546962 | 0.000000372 | -4.433937462 | 0.005486274 |
| 107052707 | *CEBPD* | -1.631670454 | 0.00000125 | -1.727844466 | 0.000035 |
| 395335 | *CISH* | -1.639177862 | 0.000000372 | -1.696539484 | 0.0000263 |
| 395399 | *CITED2* | -1.381745829 | 0.000000743 | -1.103405016 | 0.004004957 |
| 429929 | *GADD45B* | -1.634815968 | 0.000000219 | -2.1563618 | 2.67E-09 |
| 417931 | *GAS2L3* | -3.701004482 | 0.006813782 | 4.700929248 | 0.000228888 |
| 427523 | *GLDN* | -3.097751226 | 0.0000943 | 3.293259117 | 0.000110543 |
| 395925 | *KCNJ5* | -1.737522346 | 0.00000409 | -1.921886101 | 0.000027 |
| 100859468 | *LOC100859468* | -0.800572283 | 0.008487232 | -1.146377438 | 0.000111517 |
| 100859605 | *LOC100859605* | -5.750417794 | 0.0000429 | 7.526791921 | 0.00000089 |
| 769726 | *LOC769726* | -5.639373015 | 0.00000115 | -5.558226781 | 0.000420103 |
| 770169 | *LRRN3* | 0.971597164 | 0.005818832 | 1.338240942 | 0.0000949 |
| 428612 | *OLIG2* | -5.520505816 | 0.0000328 | -5.474443141 | 0.004132952 |
| 420570 | *PDK4* | -1.577439264 | 0.001286217 | -2.227524929 | 0.0000949 |
| 420198 | *PMP2* | -7.358868517 | 0.000231962 | 7.255651243 | 0.000528957 |
| 417327 | *PMP22* | -2.65744883 | 0.000628851 | 4.062040853 | 0.000000485 |
| 429116 | *PTGER3* | -1.988902181 | 0.000000219 | -2.016560884 | 0.000027 |
| 378912 | *RGS2* | -1.809462366 | 6.74E-09 | -1.617696468 | 0.000027 |
| 424038 | *S100B* | -4.309449522 | 0.000049 | 3.578834533 | 0.004132952 |
| 421766 | *SESN1* | -0.746193676 | 0.000918391 | -0.743982949 | 0.009499457 |
| 420514 | *SLC39A12* | -4.518647242 | 0.000620787 | -4.682943196 | 0.005715574 |
| 420514 | *SLC39A12* | -2.662794323 | 0.026980015 | -3.222835361 | 0.036829277 |
| 422998 | *SYT12* | -1.296634838 | 0.000231962 | -1.708983675 | 0.000027 |
| 428270 | *SYT6* | -1.198674464 | 0.000781579 | -1.582449835 | 0.0000689 |
| 771074 | *TTPA* | -1.636283666 | 0.000000538 | -1.779655548 | 0.0000124 |
| 420951 | *VSTM2A* | 1.155412738 | 0.0000541 | 1.346062256 | 0.0000263 |
| 419759 | *ZBTB16* | -1.659919857 | 0.00000125 | -2.30520479 | 9.47E-09 |

**Table 9:** Differentially expressed genes found in the gonads of both sexes for CORT.

| Entrez ID | GeneName | Female logFC | Female FDR | Male logFC | Male FDR |
|---|---|---|---|---|---|
| 427387 | SEMA6A | -1.686828569 | 0.00005 | 1.74357079 | 0.00810488 |
