## Supporting Information: Restraint Stress for "Isolating the role of corticosterone in the hypothalamic-pituitary-gonadal genomic stress response"

**Supplemental Information 2:** Restraint Stress

Restraint stress specific, differentially expressed genes by tissue and sex. Each table contains differentially expressed genes that were unique to restraint stress, which are either sex-specific or overlapping across sex. We report log fold change (logFC) and false discovery rate (FDR<0.01). Upregulated genes are highlighted in red and downregulated genes are blue.

**Table 1:** Sex-specific, restraint stress responsive genes that were differentially expressed in the female hypothalamus

| Entrez ID | Gene Name | logFC | FDR |
|---|---|---|---|
| 404536 | *ADIPOQ* | -3.7134836 | 0.000416322 |
| 418741 | *ADPRHL1* | -9.230272235 | 0.006639986 |
| 776127 | *ADSSL1* | -1.815859373 | 0.003680397 |
| 422652 | *AFP* | 1.708382347 | 0.002313855 |
| 396197 | *ALB* | 1.288828989 | 0.007339737 |
| 107049270 | *ANKRD13D* | 1.804373495 | 0.009822831 |
| 423841 | *ANKRD2* | -7.000605739 | 0.001867118 |
| 420342 | *ANXA13* | -8.076495004 | 0.005713728 |
| 424072 | *AOX2* | -4.899222927 | 0.001505985 |
| 429337 | *APLNR* | -4.303861096 | 0.00019071 |
| 769661 | *ATF7* | 1.961307191 | 0.000136202 |
| 395707 | *ATP2A3* | -2.129477967 | 0.000473 |
| 101749042 | *ATP5L* | -2.381765882 | 0.0000369 |
| 417421 | *C17ORF58* | 0.970878777 | 0.009790557 |
| 420348 | *C2H8ORF76* | 1.329957953 | 0.003095424 |
| 422853 | *C4H4ORF50* | 2.3898953 | 0.002420027 |
| 101747380 | *CAMK2N2* | 2.596297143 | 0.001582653 |
| 100359387 | *CARNS1* | 3.309399155 | 0.000445475 |
| 417701 | *CBLL1* | 1.511556882 | 0.00378503 |
| 424726 | *CCDC101* | 1.516341106 | 0.007181269 |
| 417318 | *CCDC42* | 1.178856731 | 0.00383452 |
| 101748852 | *CCDC97* | 2.716421337 | 0.006393893 |
| 395468 | *CCL4* | 2.804342008 | 0.007364241 |
| 100859215 | *CCNT1* | 1.087141606 | 0.00432082 |
| 395430 | *CDC37* | -8.376106117 | 0.000000143 |
| 416746 | *CDC42BPA* | -1.62097865 | 0.000512887 |
| 418546 | *CRYAA* | -6.715109906 | 5.98E-11 |
| 422979 | *CSRP3* | -10.71879925 | 0.006834958 |
| 426804 | *DCAF10* | -0.95563386 | 0.009790557 |
| 426247 | *DHRS4* | 0.997183864 | 0.005842423 |
| 771938 | *ESRRB* | -4.988550717 | 0.00000373 |
| 100857953 | *FAM162B* | 1.939209152 | 0.008233874 |
| 416256 | *FAXDC2* | -3.131548137 | 0.002396987 |
| 427013 | *FKBP10* | 1.985474605 | 0.00851363 |

| | | | |
|---|---|---|---|
| 420385 | *GHRHR* | 2.958454409 | 0.000661722 |
| 107050739 | *GPKOW* | 3.24189608 | 5.38E-08 |
| 427671 | *GRID2IP* | 3.236537723 | 0.00199439 |
| 424518 | *GTF2B* | 1.413056142 | 0.003101523 |
| 395813 | *HAND2* | -5.512368892 | 0.000304248 |
| 424589 | *HECTD3* | -3.866210706 | 0.002120827 |
| 429879 | *HPS1* | 1.365493518 | 0.003479737 |
| 403121 | *IFI27L2* | -7.505404732 | 0.00550626 |
| 422992 | *IFITM5* | -11.09773289 | 3.03E-10 |
| 425454 | *IGSF9* | 2.251926663 | 0.000108917 |
| 101747801 | *IKZF4* | 2.371962178 | 0.000421254 |
| 770867 | *JPH2* | -3.264068847 | 0.000683856 |
| 428314 | *KRT12* | -6.213482214 | 0.000299442 |
| 100858693 | *LOC100858693* | 2.807602482 | 0.00425458 |
| 100859084 | *LOC100859084* | 2.980918872 | 0.00240706 |
| 100859449 | *LOC100859449* | 6.709945881 | 0.003479737 |
| 101749247 | *LOC101749247* | 1.97549291 | 0.007874088 |
| 101749377 | *LOC101749377* | 0.973887737 | 0.002989826 |
| 101749825 | *LOC101749825* | 1.110955496 | 0.007774098 |
| 101750794 | *LOC101750794* | 2.756179192 | 0.009258455 |
| 101751286 | *LOC101751286* | 2.177553698 | 0.00378503 |
| 101751319 | *LOC101751319* | 4.386270965 | 0.002195021 |
| 107049137 | *LOC107049137* | 3.707967688 | 4.75E-09 |
| 107049263 | *LOC107049263* | -6.617237534 | 0.00000469 |
| 107049800 | *LOC107049800* | 6.294971508 | 0.000139847 |
| 107050461 | *LOC107050461* | 3.882664368 | 0.000299442 |
| 107050474 | *LOC107050474* | -6.647705449 | 0.001117331 |
| 107050556 | *LOC107050556* | 2.218866313 | 0.005171315 |
| 107050585 | *LOC107050585* | 1.737426115 | 0.001488346 |
| 107050614 | *LOC107050614* | 1.418754973 | 0.00028139 |
| 107050687 | *LOC107050687* | 4.464486606 | 0.000232691 |
| 107050965 | *LOC107050965* | -5.257501922 | 0.001922164 |
| 107051173 | *LOC107051173* | 2.757797576 | 0.000332186 |
| 107051182 | *LOC107051182* | 2.377920869 | 0.000726957 |
| 107051192 | *LOC107051192* | 1.45017959 | 0.005323203 |
| 107051288 | *LOC107051288* | 2.211478551 | 0.003348046 |
| 107051321 | *LOC107051321* | 2.163857996 | 0.003907505 |
| 107051325 | *LOC107051325* | 1.749374323 | 0.002313855 |
| 107051455 | *LOC107051455* | 1.683861046 | 0.002590294 |
| 107051647 | *LOC107051647* | -4.558739577 | 0.00028789 |
| 107053388 | *LOC107053388* | 1.507910741 | 0.008814146 |
| 107054855 | *LOC107054855* | 2.117402724 | 0.008026419 |
| 107055016 | *LOC107055016* | -3.930800666 | 0.001241845 |
| 107056413 | *LOC107056413* | -3.877269355 | 0.000249711 |

| | | | |
|---|---|---|---|
| 107057566 | *LOC107057566* | 4.305548415 | 0.00329965 |
| 426202 | *LRRC2* | -6.728723341 | 0.0000212 |
| 417971 | *LUC7L2* | 1.874395802 | 0.000000112 |
| 107052766 | *MATN2* | -11.99276739 | 4.5E-11 |
| 418056 | *MB* | -13.23076105 | 0.000000373 |
| 374124 | *MIP* | 2.50530927 | 0.004285187 |
| 420509 | *MLLT10* | 1.212600601 | 0.001777872 |
| 768566 | *MYH1F* | -9.542789192 | 2.25E-10 |
| 395279 | *MYH7B* | -4.657344343 | 0.0000079 |
| 771218 | *MYNN* | 0.901821828 | 0.003680397 |
| 422682 | *MYOZ2* | -5.686207005 | 0.00000184 |
| 769997 | *NAF1* | 1.063110721 | 0.00851363 |
| 386585 | *NR2F2* | 3.655120937 | 0.000798417 |
| 420996 | *NR4A3* | -1.541571637 | 0.003907505 |
| 424198 | *OBSL1* | -4.575613634 | 0.0000649 |
| 396525 | *OPN2SW* | 2.400663484 | 0.005713728 |
| 419818 | *PI16* | -2.395424196 | 0.00383452 |
| 395862 | *PITX2* | 2.057243522 | 0.003479737 |
| 423705 | *PLA2G12B* | -5.240854371 | 0.00019071 |
| 417990 | *POLR3H* | 0.789056853 | 0.003680397 |
| 396453 | *PRL* | 2.547250071 | 0.000249711 |
| 107050765 | *PRPF31* | 5.54405538 | 0.003680397 |
| 418191 | *PYROXD1* | 2.514412279 | 0.00000264 |
| 395209 | *RARRES1* | 2.078700766 | 0.000380602 |
| 396280 | *RPL27* | 2.11112904 | 0.000478923 |
| 419904 | *RPS10* | 1.393089091 | 0.003907505 |
| 429557 | *RPS6KB2* | 1.336348761 | 0.00738611 |
| 100857976 | *S1PR2* | 2.449906439 | 0.004481285 |
| 771297 | *SCN4A* | -7.282172626 | 0.005342475 |
| 415774 | *SLC7A10* | 1.196290007 | 0.000250613 |
| 107051019 | *SMG9* | 1.292691213 | 0.002867755 |
| 771780 | *SMPX* | -9.018247978 | 0.000365545 |
| 771344 | *SNRPG* | 1.818409136 | 0.004789207 |
| 422586 | *SPARCL1* | -3.518185645 | 0.000108917 |
| 419116 | *SRSF6* | 1.435218327 | 0.001241845 |
| 417321 | *STX8* | 1.512590577 | 0.006979728 |
| 422662 | *SULT1E1* | -4.44283172 | 0.00019071 |
| 426493 | *SUPT5H* | -6.701578378 | 3.91E-09 |
| 424869 | *TBCCD1* | 1.426891494 | 0.000260323 |
| 395472 | *TCIRG1* | 1.53537683 | 0.000191145 |
| 422618 | *TECRL* | -4.410856264 | 0.0000212 |
| 425829 | *TET3* | 1.926344175 | 0.002120827 |
| 419922 | *TFEB* | 1.570807686 | 0.007364241 |
| 107050764 | *TFPT* | 1.754467281 | 0.003879865 |

| | | | |
|---|---|---|---|
| 414743 | *TGFA* | 1.797828336 | 0.001057198 |
| 396106 | *TN* | 2.953496808 | 0.000228845 |
| 396433 | *TNNT2* | -13.70556682 | 0.0000212 |
| 107057622 | *UBL5* | -1.897279144 | 0.002313855 |
| 420154 | *UHRF1* | 1.23129014 | 0.007382259 |
| 417527 | *UNC45B* | -8.550969782 | 9.01E-12 |
| 423023 | *VPS37C* | 2.843241994 | 0.0000329 |
| 421674 | *VTA1* | 1.449069161 | 0.001270592 |

**Table 2:** Sex-specific, restraint stress responsive genes that were differentially expressed in the male hypothalamus.

| Entrez ID | Gene Name | logFC | FDR |
|---|---|---|---|
| 421264 | *APLF* | 0.95339217 | 0.00470254 |
| 420764 | *C2H7ORF36* | 0.85784852 | 0.00583442 |
| 417900 | *CRADD* | 1.07993306 | 0.00331244 |
| 395689 | *ENO2* | 1.37300435 | 0.00219722 |
| 770459 | *FBXW4* | 0.97703393 | 0.00331244 |
| 418589 | *GK* | -2.2639711 | 0.00470254 |
| 418328 | *GTF2E1* | 1.21840677 | 0.00219722 |
| 771273 | *MRPS36* | 1.21576339 | 0.00484729 |
| 421822 | *ORC3* | 1.14550735 | 0.00605776 |
| 415881 | *PHLPP2* | -3.6695858 | 2.41E-09 |
| 418091 | *PMCH* | 3.80947773 | 0.0000253 |
| 426311 | *RBM34* | 1.65997383 | 0.0000525 |
| 421919 | *RPS7* | 1.74623113 | 0.00047475 |
| 424250 | *SEC22A* | 1.46852196 | 0.00215916 |
| 416152 | *TTC1* | 1.23047692 | 0.0003345 |
| 421804 | *UFL1* | 1.221758 | 0.0069145 |
| 420705 | *ZDHHC3* | 0.76444263 | 0.00878401 |

**Table 3:** Sex-specific, restraint stress responsive genes of differentially expressed in the female pituitary.

| Entrez ID | Gene Name | logFC | FDR |
|---|---|---|---|
| 418623 | *ACE2* | -3.1297461 | 0.00049072 |
| 431627 | *ACOT12* | -2.1706851 | 4.96E-05 |
| 428282 | *ADAM11* | -2.0126165 | 0.00718515 |
| 416451 | *ADAP1* | -1.0768923 | 0.00466886 |
| 420386 | *ADCYAP1R1* | -1.8483194 | 0.0021005 |
| 422882 | *ADD1* | -1.0433061 | 0.00108432 |
| 107054985 | *ADD2* | -1.3800308 | 0.00026535 |
| 418155 | *ADIPOR2* | -0.7179608 | 0.00557253 |
| 423908 | *AFAP1L2* | -1.0378091 | 0.00855331 |
| 769745 | *AGBL3* | 1.01609512 | 0.00191412 |
| 421543 | *AGT* | -2.8841691 | 0.0000225 |
| 424956 | *AHSG* | -2.7426995 | 0.004088 |
| 416999 | *AIFM3* | -2.2187165 | 0.00360266 |
| 428898 | *ALKBH1* | 1.18142352 | 0.00045362 |
| 770703 | *ALKBH5* | -0.4490316 | 0.00787258 |
| 415697 | *AMFR* | -0.6475222 | 0.00686695 |
| 396311 | *ANK1* | -1.7086199 | 0.00035226 |
| 420738 | *ANLN* | -6.3295952 | 4.03E-11 |
| 419316 | *APCDD1L* | -1.4315197 | 0.00474944 |
| 396536 | *APOA1* | -4.127136 | 3.08E-07 |
| 424893 | *APOD* | -6.7632308 | 5.12E-10 |
| 417431 | *APOH* | -6.9782307 | 1.48E-09 |
| 426894 | *AQP3* | -1.3587336 | 0.00835717 |
| 421088 | *AQP4* | -4.0969943 | 2.42E-08 |
| 423846 | *ARHGAP19* | -1.7320797 | 0.00010179 |
| 429163 | *ARHGEF26* | -0.4684037 | 0.00917262 |
| 416149 | *ARHGEF37* | -2.2972096 | 0.00000261 |
| 100859297 | *ARID3B* | 0.56905707 | 0.00863815 |
| 424026 | *ARL4C* | -1.1904639 | 0.00401932 |
| 771165 | *ARRDC2* | -2.1694345 | 0.0003692 |
| 768599 | *ASAP2* | 0.85344955 | 0.00227285 |
| 417609 | *ASPA* | -1.9901558 | 0.00030443 |
| 415954 | *ASPN* | -2.2887862 | 0.00512318 |
| 422365 | *ATP1B4* | -2.6747321 | 0.00147599 |
| 414340 | *AvBD5* | -6.773035 | 9.31E-08 |
| 422077 | *BAIAP2* | -1.6064945 | 0.00431745 |
| 425976 | *BCAN* | -5.0343518 | 2.3E-09 |
| 419338 | *BCAS1* | -3.68056 | 2.76E-08 |
| 419239 | *BIRC7* | -2.1584898 | 0.0010334 |
| 395177 | *BMP10* | 4.13642243 | 0.00033568 |

| | | | |
|---|---|---|---|
| 378779 | *BMP2* | 1.24548485 | 0.00190319 |
| 771624 | *C12ORF75* | -1.6167076 | 0.00015987 |
| 101748677 | *C1H12ORF40* | 2.75287373 | 0.00031744 |
| 416156 | *C1QTNF2* | -2.5140002 | 0.0000781 |
| 419400 | *C21H1ORF222* | -2.1768382 | 0.0000106 |
| 422966 | *C5H11ORF9* | -5.58617 | 8.78E-11 |
| 423959 | *C6H10ORF90* | -4.1300399 | 2.74E-10 |
| 417179 | *C9ORF58* | -1.1019734 | 0.00628448 |
| 396257 | *CA2* | -1.5805653 | 0.00013979 |
| 373924 | *CACNG4* | -3.6389744 | 0.0000344 |
| 417429 | *CACNG5* | -2.8469814 | 0.00169175 |
| 428425 | *CALCR* | -1.8287949 | 0.00801432 |
| 101752216 | *CALII* | -2.0953096 | 0.00639807 |
| 427267 | *CAMK4L* | -1.6524279 | 0.00041966 |
| 770854 | *CAMKV* | -3.4787834 | 0.0000478 |
| 100359387 | *CARNS1* | -3.1120586 | 0.00387743 |
| 100858488 | *CARTPT* | -3.0148027 | 0.00030679 |
| 373996 | *CAV1* | -1.3323442 | 0.00409292 |
| 421019 | *CBLN2* | -1.4740426 | 0.00180642 |
| 414884 | *CCK* | -5.85894 | 1.05E-07 |
| 416155 | *CCNJL* | -1.4214383 | 0.00155063 |
| 423105 | *CD151* | -0.7151989 | 0.00650114 |
| 422827 | *CD38* | -2.4728927 | 0.0000956 |
| 419852 | *CD55* | -3.0152756 | 0.00429445 |
| 416746 | *CDC42BPA* | -1.3538875 | 0.00274061 |
| 771509 | *CDC42EP4* | -1.4782483 | 0.00030496 |
| 408049 | *CDH10* | -2.7816473 | 0.0000554 |
| 421029 | *CDH19* | -4.4961837 | 1.07E-09 |
| 419222 | *CDH4* | -0.9893025 | 0.00491522 |
| 428487 | *CDH9* | -3.0147666 | 0.00010197 |
| 416223 | *CDHR2* | -3.4409915 | 0.00758671 |
| 378914 | *CDKN1A* | -1.4923094 | 0.00305005 |
| 415969 | *CECR5L* | -1.1161519 | 0.00778298 |
| 428437 | *CHN2* | -2.2664885 | 2.58E-08 |
| 395828 | *CHRD* | -2.3169188 | 0.0095621 |
| 395399 | *CITED2* | -0.8984846 | 0.00793074 |
| 768830 | *CKS2* | -1.7416712 | 0.00060696 |
| 424990 | *CLDN11* | -3.4130519 | 4.22E-07 |
| 419595 | *CLIC4* | -0.7090392 | 0.00611141 |
| 395722 | *CLU* | -2.9767996 | 1.27E-07 |
| 421323 | *CNIH3* | -1.2114158 | 0.00179669 |
| 396522 | *CNN1* | -4.3158293 | 0.00059856 |
| 395921 | *CNP* | -5.0877837 | 4.87E-12 |
| 419825 | *CNTN2* | -2.217628 | 0.00033432 |

| | | | |
|---|---|---|---|
| 395779 | *COCH* | -1.7698884 | 0.00627443 |
| 420576 | *COL28A1* | -1.2759485 | 0.00836055 |
| 427584 | *COL7A1* | 2.91190416 | 1.34E-07 |
| 396524 | *COL9A2* | -3.7010711 | 0.00000369 |
| 421061 | *COLEC12* | -2.5866908 | 0.00000409 |
| 416783 | *COMT* | -1.1816922 | 0.00022112 |
| 428435 | *CREB5* | -3.9746793 | 4.16E-07 |
| 374218 | *CRHR1* | 1.59484269 | 0.00045227 |
| 427008 | *CRYM* | -1.8662739 | 0.00065345 |
| 425524 | *CSPG4* | -1.8730889 | 0.00758671 |
| 395466 | *CSPG5* | -2.9395403 | 0.00155063 |
| 396176 | *CSRP1* | -2.2434942 | 0.00017915 |
| 772072 | *CTD-2510F5.6* | 1.34407512 | 0.00022068 |
| 374135 | *CTGF* | -2.0453262 | 0.0000627 |
| 417124 | *CUTA* | -1.9205205 | 0.000036 |
| 422058 | *CYP39A1* | -2.6863992 | 0.00033002 |
| 421422 | *DAAM2* | -2.7104147 | 1.99E-08 |
| 419180 | *DBNDD2* | -1.9573277 | 0.00000656 |
| 107053650 | *DDIT4* | -2.7488128 | 9.40E-08 |
| 395906 | *DES* | -2.3715637 | 0.00360985 |
| 425241 | *DIRAS2* | -1.4094971 | 0.00422776 |
| 422740 | *DLC1* | -0.735029 | 0.00436062 |
| 107054041 | *DMRTA2* | 1.65703371 | 0.00173647 |
| 421340 | *DUSP10* | -0.8802786 | 0.00238833 |
| 431336 | *DUSP16* | -0.8486377 | 0.00181133 |
| 425353 | *DYSF* | 1.81918851 | 6.11E-07 |
| 427326 | *EDIL3* | -1.4388315 | 0.00110876 |
| 416619 | *EEF2K* | -0.7456012 | 0.00304048 |
| 408035 | *EGF* | -2.3473544 | 0.00000809 |
| 416093 | *EIF4E3* | -3.1583191 | 5.5E-08 |
| 427906 | *ELFN2* | -3.4084986 | 9.42E-08 |
| 417604 | *EMC6* | -1.4168988 | 0.00780016 |
| 396017 | *ENO1* | -1.3329856 | 0.00374572 |
| 420361 | *ENPP2* | -1.4319124 | 0.0000251 |
| 422551 | *ENPP6* | -2.7818022 | 3.56E-06 |
| 395797 | *ENTPD2* | -1.7246718 | 0.00012641 |
| 395671 | *ERBB4* | -3.0700231 | 6.11E-07 |
| 771826 | *ERMN* | -4.3689238 | 2.65E-08 |
| 426690 | *ETNK2* | -4.1570752 | 1.90E-08 |
| 429084 | *F3* | -1.1922879 | 0.00800761 |
| 415687 | *FA2H* | -5.3629261 | -5.3629261 |
| 396246 | *FABP7* | -1.8899298 | 0.00470526 |
| 771797 | *FAM101A* | -1.3144505 | 0.00209368 |
| 418939 | *FAM123A* | -1.79612 | 7.21E-07 |

| | | | |
|---|---|---|---|
| 770395 | *FAM124A* | -2.3831021 | 0.0000781 |
| 424451 | *FAM129A* | -1.053501 | 0.003706 |
| 424644 | *FAM159A* | -2.6756803 | 0.00091508 |
| 101751376 | *FAM181B* | -1.9592269 | 0.0000002 |
| 422413 | *FAM198B* | -2.93007 | 0.00371963 |
| 419043 | *FAR2* | -1.6183396 | 0.00000421 |
| 395167 | *FAT3* | -1.5431702 | 0.00152694 |
| 373979 | *FBLN1* | -1.2656528 | 0.00554261 |
| 423413 | *FBLN5* | -1.5501848 | 0.00787258 |
| 420343 | *FBXO32* | -1.0475075 | 0.00000483 |
| 769464 | *FCHO2* | 1.86841427 | 0.0000102 |
| 374020 | *FECH* | -0.9613865 | 0.00360266 |
| 417908 | *FGD6* | -1.5125347 | 0.0000741 |
| 421860 | *FILIP1* | -1.5725231 | 0.00198874 |
| 771739 | *FMNL2* | -1.3406381 | 0.0000407 |
| 395814 | *FMOD* | -3.2503528 | 0.00013316 |
| 417335 | *FN3K* | -1.412922 | 0.00010379 |
| 100857683 | *FNTB* | -1.6484446 | 0.00078958 |
| 415576 | *FRMD5* | -1.4037334 | 0.00012527 |
| 770406 | *FXYD6* | -2.2287055 | 0.00000625 |
| 374060 | *FZD7* | -1.1746203 | 0.00534579 |
| 422456 | *GAB1* | -1.2112775 | 0.00031115 |
| 770511 | *GABRA2* | -2.5347862 | 0.00000288 |
| 422289 | *GABRA3* | -2.6104935 | 0.0000193 |
| 422770 | *GABRA4* | -1.1207767 | 0.00413008 |
| 770888 | *GAREML* | -4.3590487 | 0.00013022 |
| 415404 | *GCOM1* | -0.7300382 | 0.00501581 |
| 419084 | *GDPD4* | -2.1601186 | 0.00278465 |
| 404771 | *GEM* | -1.2127712 | 0.004088 |
| 419969 | *GFAP* | -7.929595 | 9.40E-12 |
| 416945 | *GGT1* | -3.0849358 | 4.69E-07 |
| 419178 | *GHRH* | -3.6548799 | 9.31E-08 |
| 395278 | *GJA1* | -1.7750978 | 0.0000103 |
| 378797 | *GJB1* | -5.8250316 | 0.0000552 |
| 420397 | *GJC2* | -2.9251607 | 4.70E-08 |
| 427523 | *GLDN* | -1.4304639 | 0.00856022 |
| 772019 | *GLRA2* | -3.6102582 | 0.0000187 |
| 769706 | *GLRA4* | -2.8590234 | 0.0000157 |
| 396489 | *GLUL* | -4.0305311 | 7.05E-11 |
| 100857704 | *GMNC* | -3.1792237 | 0.00093742 |
| 107051576 | *GP1BB* | -1.948306 | 0.0000555 |
| 422234 | *GPC4* | -1.9106967 | 2.03E-07 |
| 418632 | *GPM6B* | -1.4035089 | 0.00024018 |
| 428932 | *GPR137C* | -1.4993713 | 0.0003488 |

| | | | |
|---|---|---|---|
| 769024 | *GPR17* | -2.9853884 | 0.00025042 |
| 417748 | *GPR37* | -4.8029829 | 2.76E-08 |
| 421176 | *GPR37L1* | -5.1350853 | 2.43E-07 |
| 101747453 | *GPRC5B* | -2.0295322 | 0.00000105 |
| 415746 | *GPT2* | -0.7203764 | 0.00512499 |
| 373933 | *GRM5* | -3.0566117 | 0.00000466 |
| 423988 | *GULP1* | -1.7438912 | 0.0026923 |
| 693250 | *H1F0* | -1.9352823 | 0.00036754 |
| 423721 | *H2AFY2* | 1.17093984 | 0.00143889 |
| 418311 | *HAO2* | -6.2461551 | 0.0000297 |
| 425975 | *HAPLN2* | -7.3679893 | 7.05E-11 |
| 424441 | *HEBP2* | -3.297287 | 4.98E-10 |
| 427378 | *HEMGN* | -2.4743976 | 0.00017915 |
| 428234 | *HEPACAM* | -4.0277005 | 2.55E-06 |
| 771101 | *HEYL* | -2.1341499 | 0.0000317 |
| 396287 | *HMOX1* | -1.8795098 | 0.00021142 |
| 395863 | *HPGDS* | -3.0941895 | 0.00297719 |
| 100858928 | *HRH1* | 0.95441728 | 0.00859669 |
| 428310 | *HSP25* | -2.4090404 | 0.00127938 |
| 395853 | *HSPA8* | -1.1853252 | 0.0000201 |
| 431581 | *HTR1A* | -4.4782456 | 0.00012984 |
| 374150 | *ID4* | -1.4085235 | 0.00470752 |
| 396315 | *IGFBP2* | -2.5385596 | 0.00000356 |
| 424220 | *IGFBP5* | -1.496201 | 0.00022537 |
| 423492 | *INF2* | -1.4855444 | 0.00148022 |
| 429941 | *ISLR2* | -3.2004859 | 0.00041966 |
| 395470 | *ITGB8* | -1.0853299 | 0.00413222 |
| 419110 | *ITIH2* | -4.1552788 | 0.0000002 |
| 395694 | *ITPKA* | -1.264422 | 0.00110049 |
| 417355 | *JMJD6* | -0.8453023 | 0.00012049 |
| 396300 | *KBP* | -3.5993082 | 3.75E-09 |
| 100858067 | *KCNA6* | -2.1602699 | 0.00250193 |
| 100857799 | *KCNJ10* | -4.0683626 | 0.00000369 |
| 427662 | *KCNJ12* | -4.2359915 | 1.43E-06 |
| 421424 | *KCNK5* | -1.9325955 | 0.00324347 |
| 395301 | *KCNMB1* | -3.1046803 | 0.00030133 |
| 395248 | *KCNT1* | -1.2643616 | 0.00695854 |
| 424434 | *KIAA0040* | -2.0127429 | 0.00000817 |
| 418854 | *KIAA0226L* | -1.3378601 | 0.0001687 |
| 422374 | *KIAA1210* | -1.4805042 | 0.00013316 |
| 420476 | *KIAA1462* | -1.6742373 | 0.00015415 |
| 768701 | *KIAA1671* | -0.5693561 | 0.00826564 |
| 419968 | *KIF18B* | -1.939094 | 0.00959591 |
| 421801 | *KLHL32* | 1.18221783 | 0.0000817 |

| | | | |
|---|---|---|---|
| 396221 | *LDHA* | -0.8816742 | 0.00315261 |
| 395705 | *LHX2* | -2.41938 | 0.00316411 |
| 100858027 | *LLGL1* | -1.4689814 | 0.00106089 |
| 416113 | *LMCD1* | -2.6107241 | 0.00048879 |
| 100857927 | *LOC100857927* | -2.4794081 | 0.0000133 |
| 100858941 | *LOC100858941* | -2.4603547 | 0.00448207 |
| 100859224 | *LOC100859224* | -6.5802979 | 1.94E-09 |
| 100859848 | *LOC100859848* | -3.2593545 | 3.32E-09 |
| 100859906 | *LOC100859906* | -4.3137274 | 1.05E-07 |
| 101747844 | *LOC101747844* | -3.0905492 | 0.00044219 |
| 427408 | *LOC101747901* | -3.3798149 | 0.0005672 |
| 101748788 | *LOC101748788* | -4.9677128 | 0.00032715 |
| 101748987 | *LOC101748987* | -2.3876973 | 0.00020293 |
| 101750794 | *LOC101750794* | 2.79017741 | 0.00367527 |
| 101751203 | *LOC101751203* | -1.685082 | 0.00469958 |
| 101752135 | *LOC101752135* | -3.4527743 | 0.00155063 |
| 107050516 | *LOC107050516* | -4.4047463 | 2.00E-07 |
| 107050828 | *LOC107050828* | -3.4095629 | 0.0000477 |
| 107050945 | *LOC107050945* | -1.8681064 | 0.0000744 |
| 107051100 | *LOC107051100* | 1.16484004 | 0.00885896 |
| 107051857 | *LOC107051857* | 0.73242457 | 0.00243979 |
| 107052456 | *LOC107052456* | -3.9597286 | 0.00064488 |
| 107056412 | *LOC107056412* | -3.2534036 | 0.0000291 |
| 396098 | *LOC396098* | -2.4010436 | 0.00117598 |
| 416927 | *LOC416927* | -5.8510276 | 9.73E-06 |
| 417962 | *LOC417962* | -1.9069947 | 0.00040964 |
| 418109 | *LOC418109* | -3.0530045 | 0.0000193 |
| 420030 | *LOC420030* | -3.3883832 | 0.0000181 |
| 420209 | *LOC420209* | 3.00424686 | 0.00548213 |
| 421298 | *LOC421298* | -3.7338635 | 0.00084076 |
| 421584 | *LOC421584* | -4.1681292 | 0.00010516 |
| 422305 | *LOC422305* | -2.8862162 | 0.0000341 |
| 422321 | *LOC422321* | -2.8235973 | 0.0000315 |
| 422323 | *LOC422323* | -1.3051574 | 0.00707106 |
| 425001 | *LOC425001* | -1.988993 | 0.0000575 |
| 427201 | *LOC427201* | -1.0569485 | 0.0085929 |
| 427400 | *LOC427400* | -1.9921765 | 0.00628811 |
| 428754 | *LOC428754* | -1.0046045 | 0.00155063 |
| 769726 | *LOC769726* | -6.9301756 | 1.81E-10 |
| 770718 | *LOC770718* | -2.3885342 | 0.000016 |
| 771456 | *LOC771456* | -2.770959 | 0.0000228 |
| 427686 | *LRRN4* | -1.7984747 | 0.00232551 |
| 770619 | *LTC4S* | -3.5242913 | 0.00207691 |
| 424192 | *LY75* | -2.3105733 | 2.76E-08 |

| | | | |
|---|---|---|---|
| 771649 | *LYPD6* | -2.6729829 | 0.0000611 |
| 396097 | *MAP4* | -1.2925258 | 0.00596172 |
| 423770 | *MARCH8* | -1.3914879 | 0.00000162 |
| 396217 | *MBP* | -3.92078 | 4.37E-12 |
| 427126 | *MEGF10* | -0.9962598 | 0.00474944 |
| 374137 | *MEOX2* | -2.0204962 | 0.00422776 |
| 426871 | *METTL7A* | -1.9825318 | 0.00000356 |
| 415494 | *MFGE8* | -1.3104356 | 0.00330048 |
| 771284 | *MKX* | -2.6342822 | 0.0000295 |
| 417737 | *MLC1* | -3.9822063 | 2.76E-08 |
| 395847 | *MLLT11* | -1.5565072 | 0.00376189 |
| 416480 | *MMD2* | -1.8456714 | 0.0000132 |
| 418627 | *MOSPD2* | -0.8312987 | 0.0059203 |
| 395333 | *MOXD1* | -1.5563025 | 0.00835717 |
| 420006 | *MPP2* | -4.1825098 | 1.17E-08 |
| 100858795 | *MRPL40* | -0.8041952 | 0.00278938 |
| 396484 | *MSX1* | -4.6887598 | 0.00000247 |
| 396211 | *MYH11* | -3.9302377 | 0.00000417 |
| 418751 | *MYO16* | -2.9737718 | 0.0000274 |
| 417341 | *NARF* | -0.6866989 | 0.00346869 |
| 373952 | *NBL1* | -1.4046463 | 0.00110876 |
| 395493 | *NCAN* | -5.091642 | 8.46E-07 |
| 416597 | *NDE1* | -1.2767299 | 0.00285819 |
| 422672 | *NDNF* | -1.4010932 | 0.00078573 |
| 418560 | *NDP* | -3.5203485 | 0.000017 |
| 420321 | *NDRG1* | -1.2074996 | 0.0000972 |
| 420219 | *NECAB1* | -3.7023341 | 0.0000139 |
| 396206 | *NEFM* | -3.0823817 | 0.0000266 |
| 423977 | *NEMP2* | 0.66883622 | 0.00174195 |
| 395890 | *NES* | -2.9709781 | 0.00079094 |
| 416693 | *NET1* | -1.1553311 | 0.00015558 |
| 418151 | *NINJ2* | -6.0772711 | 7.05E-11 |
| 421720 | *NKAIN2* | -2.8222921 | 0.0000108 |
| 415729 | *NKD1* | -1.3476143 | 0.00012797 |
| 395591 | *NKX6-2* | -2.4947811 | 0.0000271 |
| 416438 | *NPTX2* | -1.9486747 | 0.0021005 |
| 396464 | *NPY* | -2.8266281 | 0.0010334 |
| 422405 | *NPY2R* | -1.8118087 | 0.00296869 |
| 693264 | *NPY6R* | -8.4578559 | 5.78E-10 |
| 396082 | *NR2E1* | -1.7501466 | 0.0067973 |
| 421833 | *NT5E* | -3.2903079 | 4.39E-06 |
| 417914 | *NTN4* | -1.3605587 | 0.00094382 |
| 396157 | *NTRK2* | -1.0780971 | 0.00025042 |
| 417883 | *NTS* | -1.5015832 | 0.00594519 |

| | | | |
|---|---|---|---|
| 428612 | *OLIG2* | -7.7767413 | 1.80E-11 |
| 101751011 | *OMG* | -3.5548374 | 2.52E-08 |
| 396486 | *OPN1MSW* | 2.65883344 | 0.00099129 |
| 395334 | *OPN4-1* | -5.7387218 | 3.54E-07 |
| 423974 | *OPNVA* | -2.7468583 | 0.00027486 |
| 429951 | *OPRD1* | -2.301814 | 0.0098975 |
| 415706 | *OSGIN1* | -0.9442717 | 0.00272934 |
| 554220 | *OTP* | -3.6390992 | 0.00937928 |
| 420502 | *OTUD1* | -0.8939909 | 0.00852694 |
| 426041 | *OTUD7B* | -1.3334993 | 0.00081426 |
| 396275 | *P2RY1* | -1.7411615 | 0.00010116 |
| 395910 | *PADI3* | -3.7309815 | 7.06E-08 |
| 425505 | *PAQR7* | -1.0201674 | 0.0021204 |
| 422042 | *PAQR8* | -1.330784 | 0.00022068 |
| 374127 | *PAX3* | -4.1663916 | 0.00331046 |
| 428065 | *PCDH8* | -3.6329915 | 0.00147599 |
| 107057452 | *PCP4L1* | -3.6040333 | 0.00391354 |
| 421131 | *PENK* | -4.0211142 | 0.0000271 |
| 418247 | *PFKFB3* | 0.67614186 | 0.00285819 |
| 427215 | *PGM5* | -1.5704311 | 0.00198874 |
| 424381 | *PHGDH* | -2.9571061 | 0.0007764 |
| 420904 | *PHLPP1* | -1.1656561 | 0.0000236 |
| 418181 | *PIK3C2G* | -1.2291935 | 0.00938514 |
| 771808 | *PIPOX* | -5.0864391 | 9.15E-05 |
| 420645 | *PLCL2* | -2.1649576 | 0.00000424 |
| 415650 | *PLLP* | -7.0956819 | 4.88E-11 |
| 428164 | *PLTP* | -1.8054032 | 0.00013507 |
| 420198 | *PMP2* | -4.794516 | 0.000377 |
| 417327 | *PMP22* | -2.2673437 | 2.16E-07 |
| 415809 | *PNAT3* | -1.6994693 | 0.00333513 |
| 422019 | *PNOC* | -4.2723341 | 0.00000312 |
| 107053103 | *POU3F2* | -3.7652877 | 0.00684632 |
| 107051987 | *POU3F3* | -5.4061833 | 2.93E-08 |
| 420753 | *PPP1R17* | 1.50258427 | 0.00716526 |
| 416350 | *PPP2R2B* | -0.8846606 | 0.00995558 |
| 419681 | *PPT1* | -0.8096993 | 0.00501581 |
| 417183 | *PRDM12* | -3.9575753 | 0.00254648 |
| 426800 | *PRDM6* | -4.6737927 | 0.0000817 |
| 419933 | *PRELP* | -1.5618838 | 0.00096762 |
| 421405 | *PREPL* | 0.75220259 | 0.00270855 |
| 772317 | *PRIMA1* | -2.6252603 | 0.00399585 |
| 374110 | *PTGDS* | -3.4909675 | 0.00000963 |
| 421174 | *PTPRVP* | -2.0850796 | 0.00088029 |
| 548626 | *PTX3* | -3.9428753 | 0.00091676 |

| | | | |
|---|---|---|---|
| 374204 | *QKI* | -1.7723936 | 4.76E-07 |
| 419020 | *RAB30* | -1.1327022 | 0.00918585 |
| 100857462 | *RAPGEF5* | -0.6405184 | 0.00179669 |
| 421460 | *RASGRP3* | -1.8580502 | 0.0000552 |
| 771592 | *RASL10B* | -1.998694 | 0.0007565 |
| 415530 | *RASL12* | -2.6783823 | 0.00051661 |
| 419521 | *RASSF2* | -4.1664749 | 9.38E-09 |
| 768866 | *RBM38* | -1.3213028 | 0.00061543 |
| 395678 | *RBPMS2* | -3.3118437 | 0.0000915 |
| 422057 | *RCAN2* | -1.1593896 | 0.00224478 |
| 427850 | *RELN* | -1.6003625 | 0.00477515 |
| 418070 | *RFX4* | -2.8113387 | 0.00015244 |
| 426557 | *RGS8* | -2.7170983 | 0.0000412 |
| 416647 | *RHBDF1* | -0.9171363 | 0.00280928 |
| 395734 | *RHOB* | -1.2326409 | 0.0000916 |
| 420251 | *RNF19A* | -0.8852591 | 0.00089088 |
| 420046 | *RP5-1028K7.3* | -2.1522758 | 0.00051661 |
| 415790 | *RRAD* | -1.8289886 | 0.00071031 |
| 424038 | *S100B* | -4.4545983 | 5.98E-08 |
| 417146 | *SARDH* | -2.620262 | 0.00065049 |
| 395706 | *SCD* | -1.148659 | 0.00096762 |
| 425016 | *SCHIP1* | -0.7928191 | 0.00926346 |
| 419774 | *SCN4B* | -3.9846254 | 4.88E-07 |
| 422563 | *SCRG1* | -3.7529461 | 0.0000727 |
| 374102 | *SDC2* | -1.0580788 | 0.00610647 |
| 419184 | *SDC4* | -0.9290366 | 0.00390259 |
| 415435 | *SECISBP2L* | -0.6530168 | 0.00475428 |
| 396332 | *SEMA3D* | -2.5404374 | 2.75E-07 |
| 424593 | *SEPP1L* | -5.4251148 | 5.29E-09 |
| 416778 | *SEPT5* | -3.3187374 | 7.05E-11 |
| 395877 | *SERPIND1* | -3.9974434 | 0.00000229 |
| 107049626 | *SFTPC* | -6.18283 | 8.09E-06 |
| 422954 | *SFXN5* | -1.093801 | 0.00922745 |
| 395133 | *SGK1* | -0.9226814 | 0.0038318 |
| 422632 | *SGK223* | -1.3211948 | 0.00501581 |
| 418731 | *SH3RF3* | -1.8609763 | 0.00138055 |
| 396089 | *SHANK3* | -7.3111408 | 1.24E-10 |
| 395615 | *SHH* | -2.6627442 | 0.0007565 |
| 418807 | *SLAIN1* | -1.7498453 | 0.00018084 |
| 427123 | *SLC12A2* | -1.0414258 | 0.00550602 |
| 770495 | *SLC13A3* | -6.1348152 | 0.00000013 |
| 417678 | *SLC13A5* | -1.3544811 | 0.00166073 |
| 423156 | *SLC1A2* | -1.1084656 | 0.0060913 |
| 395443 | *SLC1A3* | -1.6096097 | 0.00087772 |

| | | | |
|---|---|---|---|
| 419530 | *SLC25A37* | -1.2925585 | 0.00051439 |
| 426008 | *SLC27A1* | -2.0738376 | 0.00000652 |
| 396130 | *SLC2A1* | -1.3745597 | 0.0000323 |
| 419167 | *SLC32A1* | -5.2142884 | 4.3E-09 |
| 417809 | *SLC38A4* | -4.4588815 | 0.00000384 |
| 417557 | *SLC43A2* | -0.8300134 | 0.00010799 |
| 429117 | *SLC44A5* | -0.9669969 | 0.00013022 |
| 771751 | *SLC45A3* | -1.9628529 | 0.00248455 |
| 417616 | *SLC47A2* | -4.2022344 | 0.00452713 |
| 422649 | *SLC4A4* | -0.9685743 | 0.0057525 |
| 416562 | *SLC5A11* | -3.7353525 | 0.00000776 |
| 414870 | *SLC5A7* | -2.8593443 | 0.004088 |
| 418147 | *SLC6A12* | -2.2832907 | 0.00000523 |
| 416277 | *SLC6A7* | -4.2476164 | 0.0000238 |
| 100861584 | *SLC6A8* | -1.2633427 | 0.00069786 |
| 424576 | *SLC6A9* | -4.3683153 | 2.29E-09 |
| 395759 | *SLC8A3* | -1.3943517 | 0.00029075 |
| 419356 | *SLC9A8* | 0.58965308 | 0.00374572 |
| 418187 | *SLCO1B1* | -3.1239085 | 0.00010116 |
| 395293 | *SLIT1* | -2.9804896 | 2.43E-07 |
| 395949 | *SMO* | -1.075377 | 0.00309867 |
| 422934 | *SMOX* | -1.1136601 | 0.00028299 |
| 100858894 | *SMTN* | -1.9331897 | 0.00011741 |
| 416487 | *SMURF1* | -1.5811804 | 0.00000279 |
| 395393 | *SNCA* | -3.7444804 | 7.06E-08 |
| 395392 | *SNCG* | -1.4638679 | 0.00955029 |
| 395573 | *SOX10* | -6.8385771 | 6.17E-09 |
| 107051027 | *SOX21* | -4.7439823 | 0.00000862 |
| 395483 | *SOX8* | -3.7436611 | 1.55E-09 |
| 417595 | *SPECC1* | -0.9867755 | 0.00155063 |
| 418902 | *SPG20* | -0.828427 | 0.00926346 |
| 395657 | *SPON1* | -2.4388 | 5.44E-07 |
| 424741 | *SSPO* | -1.7815033 | 0.00000161 |
| 395421 | *STAR* | 2.05857487 | 0.0002509 |
| 423238 | *STARD9* | -1.1398665 | 0.00643277 |
| 422637 | *STBD1* | -2.0524491 | 0.0000173 |
| 420775 | *STK17A* | -0.7824739 | 0.00999605 |
| 396057 | *STMN1* | -1.2166284 | 0.00112794 |
| 422010 | *STMN4* | -2.5926782 | 3.04E-08 |
| 423390 | *STON2* | -0.7553553 | 0.00640385 |
| 415988 | *SUSD3* | -1.526683 | 0.00181133 |
| 423044 | *SWAP70* | -1.486767 | 0.00001 |
| 428768 | *SYNPO2* | -2.8165188 | 0.00000553 |
| 420573 | *TAC1* | -3.1656164 | 0.00011615 |

| | | | |
|---|---|---|---|
| 396490 | *TAGLN* | -4.9306523 | 0.00000685 |
| 396298 | *TAL1* | -1.5623012 | 0.00425039 |
| 395104 | *TBX18* | -1.8609251 | 0.00548703 |
| 373895 | *TBX22* | -2.4791014 | 0.00574606 |
| 771113 | *TESC* | -1.9400221 | 0.00172723 |
| 395982 | *TFAP2A* | 1.99294344 | 0.00182966 |
| 768711 | *TIMP4* | -2.9518804 | 9.38E-09 |
| 395751 | *TJP2* | -0.8872209 | 0.00231215 |
| 771874 | *TMBIM4* | -1.2252653 | 0.00049334 |
| 416892 | *TMEM119* | -3.0247458 | 0.00311741 |
| 424560 | *TMEM125* | -2.4749628 | 0.0000187 |
| 771698 | *TMEM81* | -1.9106726 | 0.00060622 |
| 419414 | *TMEM88B* | -5.692252 | 1.55E-09 |
| 427988 | *TMPRSS3* | 1.6383776 | 0.00825729 |
| 427499 | *TNFAIP8L3* | -2.482272 | 0.00042493 |
| 396032 | *TNNC1* | -5.6713539 | 2.00E-06 |
| 420800 | *TPPP* | -1.3134142 | 0.00033002 |
| 423471 | *TRAF3* | -1.391274 | 0.00274106 |
| 425131 | *TRIM3* | -1.2723085 | 0.00897966 |
| 428900 | *TSHR* | -3.6637132 | 3.70E-05 |
| 424543 | *TTLL7* | -1.1413286 | 0.00155339 |
| 100858879 | *TTYH2* | -3.8611993 | 1.72E-09 |
| 396427 | *TUBB1* | -0.9319689 | 0.00581465 |
| 416678 | *UBE2H* | -0.4710867 | 0.00566643 |
| 374033 | *UGT8* | -3.4963169 | 2.49E-09 |
| 417221 | *URM1* | -1.6655933 | 0.00460309 |
| 424467 | *VCAM1* | -2.1012374 | 0.00013507 |
| 395565 | *VCAN* | -2.1674619 | 0.00026608 |
| 420519 | *VIM* | -2.4306511 | 0.00000132 |
| 396323 | *VIP* | -4.3381882 | 0.00000252 |
| 417635 | *VMP1* | -1.0010543 | 0.00027602 |
| 423782 | *VSTM4* | -1.6827131 | 0.00058649 |
| 424547 | *VTG1* | 3.75398864 | 0.00024532 |
| 419413 | *VWA1* | -2.254335 | 4.16E-07 |
| 420944 | *VWC2* | -2.4045732 | 0.00825729 |
| 418829 | *WBP4* | -0.6222235 | 0.00511432 |
| 427497 | *WDR72* | -3.9939251 | 0.000083 |
| 417831 | *WIF1* | -1.8637682 | 0.00547929 |
| 395562 | *WNT11* | -1.4145576 | 0.00525833 |
| 395235 | *WNT6* | -5.30917 | 0.0000225 |
| 427937 | *WNT7B* | -3.8599505 | 0.00581618 |
| 100857831 | *YJEFN3* | -5.0368982 | 0.0000817 |
| 424306 | *ZEB2* | -1.5929106 | 0.00021054 |
| 374103 | *ZIC1* | -3.3459738 | 0.000054 |

| | | | |
|---|---|---|---|
| 428021 | *ZIC2* | -6.9046595 | 2.3E-09 |
| 422251 | *ZIC3* | -4.3096862 | 0.0000443 |
| 424885 | *ZIC4* | -4.354597 | 0.0000324 |
| 420651 | *ZNF385D* | -1.0546122 | 0.00265575 |

**Table 4:** Sex-specific, restraint stress responsive genes of differentially expressed in the male pituitary.

| Entrez ID | Gene Name | logFC | FDR |
|---|---|---|---|
| 396024 | *ACTN4* | 1.60317412 | 0.00817479 |
| 418479 | *ADAMTS1* | -2.0643209 | 0.0000121 |
| 423548 | *AP5M1* | 2.00435349 | 0.00865146 |
| 386573 | *ASCL1* | 1.38414663 | 0.00619581 |
| 417481 | *AUTS2* | 1.66995437 | 0.00450389 |
| 421779 | *C3H6ORF203* | 0.9887876 | 0.0013641 |
| 422853 | *C4H4ORF50* | 2.34723784 | 0.00188576 |
| 107051972 | *CCDC138* | 1.68116704 | 0.00910379 |
| 416044 | *CCDC174* | 0.97876197 | 0.00128007 |
| 416613 | *CCP110* | 0.68424101 | 0.00948447 |
| 417730 | *CD36* | 1.60685176 | 0.0044556 |
| 418461 | *CHMP2B* | 0.82526661 | 0.00826393 |
| 424095 | *CPO* | -3.0664577 | 0.00207013 |
| 417117 | *DAB2IP* | -1.1668448 | 0.00483383 |
| 424187 | *DPP4* | -2.4297595 | 0.00270677 |
| 423733 | *DUPD1* | 3.04837727 | 0.00116191 |
| 374192 | *DUSP1* | -1.1483094 | 0.00938244 |
| 423890 | *DUSP5* | -1.8823196 | 0.0000623 |
| 431579 | *ELOVL7* | -1.52569 | 0.00079554 |
| 422608 | *FAM175A* | 1.13188071 | 0.00089404 |
| 770787 | *FHL1* | 2.92652911 | 0.00053047 |
| 419860 | *G0S2* | 3.12591438 | 0.00202266 |
| 378911 | *GHSR* | 3.89426529 | 0.00053144 |
| 395273 | *GJD2* | -2.221682 | 0.0000149 |
| 422772 | *GNPDA2* | 1.57255292 | 0.00142952 |
| 771199 | *GPR34* | 2.78644469 | 0.00657757 |
| 427887 | *H3F3C* | 0.91435056 | 0.00860256 |
| 396227 | *HSPB1* | 2.76211269 | 0.00826393 |
| 101747801 | *IKZF4* | 1.74827243 | 0.00797275 |
| 770238 | *KLF9* | -0.8690342 | 0.0036568 |
| 423824 | *KNDC1* | 1.96404241 | 0.00059636 |
| 100857380 | *LOC100857380* | -1.4335824 | 0.0013641 |
| 100858647 | *LOC100858647* | -3.1461843 | 0.00415232 |
| 100859468 | *LOC100859468* | -1.1201188 | 0.004921 |
| 101749060 | *LOC101749060* | -1.1122871 | 0.00120214 |
| 101750533 | *LOC101750533* | 1.62230155 | 0.00042636 |
| 107049255 | *LOC107049255* | 2.91962133 | 0.00082958 |
| 107049800 | *LOC107049800* | 5.47500389 | 0.0000149 |
| 107050569 | *LOC107050569* | 1.81314457 | 0.00580654 |
| 107050718 | *LOC107050718* | 1.60242387 | 0.00430068 |

| | | | |
|---|---|---|---|
| 107054798 | *LOC107054798* | -2.1821401 | 0.0027726 |
| 417873 | *MYF6* | 5.54715704 | 0.0000665 |
| 417506 | *MYL10* | 2.72184828 | 0.00370237 |
| 420893 | *MYLK4* | 3.00824604 | 0.00031589 |
| 429272 | *MYO7L2* | 3.07396689 | 0.00690999 |
| 395805 | *MYOM1* | 3.43774423 | 0.0000322 |
| 423744 | *MYOZ1* | 3.55450943 | 0.00020771 |
| 416659 | *NAA60* | 0.60144236 | 0.0054821 |
| 374027 | *NEB* | 2.23585373 | 0.00059636 |
| 386585 | *NR2F2* | 3.20490335 | 0.00255123 |
| 420582 | *NXPH1* | 1.40045497 | 0.00657757 |
| 396525 | *OPN2SW* | 2.4682132 | 0.00127084 |
| 428607 | *PCMT1* | 1.35829484 | 0.00620683 |
| 769230 | *PCSK1* | -1.0197284 | 0.00650208 |
| 416912 | *PITPNB* | 0.88083529 | 0.00103383 |
| 423324 | *PPP2R3C* | 1.20755191 | 0.00017692 |
| 396459 | *PVALB* | 3.4852614 | 9.54E-05 |
| 422756 | *RASL11B* | -1.0654673 | 0.00291405 |
| 107050161 | *RENBP* | 2.260811 | 0.00254265 |
| 419669 | *RNF19B* | -0.9608657 | 0.00080023 |
| 427281 | *RNF38* | 1.95479739 | 0.00127084 |
| 404773 | *RP5-966M1.6* | 3.20061649 | 0.00361116 |
| 374006 | *SAT1* | -1.0024146 | 0.0016801 |
| 395946 | *SCN9A* | 1.14688352 | 0.0061185 |
| 777244 | *SHOX2* | 1.9543518 | 0.00603903 |
| 395583 | *SPRY1* | -0.9538911 | 0.00127084 |
| 424475 | *SSPO* | 1.73150732 | 0.00681807 |
| 396327 | *SYT2* | 1.21353445 | 0.00993951 |
| 418721 | *TMEM182* | 3.61608458 | 0.0000446 |
| 395883 | *TMOD1* | 1.63755948 | 0.00103383 |
| 396106 | *TN* | 2.46866482 | 0.00109198 |
| 396386 | *TNNI2* | 3.50924225 | 0.00437694 |
| 421721 | *TRDN* | 5.2194338 | 0.0000663 |
| 395766 | *USP2* | -0.9564947 | 0.00657757 |
| 428580 | *ZBTB18* | 0.80535379 | 0.0000993 |
| 422847 | *ZBTB49* | 1.32491754 | 0.00436677 |
| 422465 | *ZNF827* | 0.75780671 | 0.00718952 |
| 770670 | *ZSWIM6* | 1.49050803 | 0.00265528 |

**Table 5:** Sex-specific, restraint stress responsive genes that were differentially expressed in the female gonads.

| Entrez ID | Gene Name | logFC | FDR |
|---|---|---|---|
| 418254 | *A2ML1* | 2.71248959 | 0.00025841 |
| 416811 | *AACS* | 0.67751373 | 0.00466832 |
| 418975 | *AASDHPPT* | 0.66794428 | 0.00366279 |
| 373945 | *ABCA1* | 3.30570908 | 0.0037845 |
| 420673 | *ABHD5* | -0.6848438 | 0.00107753 |
| 420489 | *ABI1* | 0.90084263 | 0.00366657 |
| 417850 | *AC025263.3* | 1.58817547 | 0.00000239 |
| 421317 | *ACBD3* | 0.55934801 | 0.00449219 |
| 769222 | *ADAMTS8* | 0.89636176 | 0.00933488 |
| 420328 | *ADCY8* | -1.9230066 | 0.00045043 |
| 421618 | *ADGB* | -1.5689045 | 0.00086759 |
| 101750527 | *ADGRF5* | -1.2541622 | 0.00054172 |
| 421036 | *AFG3L2* | 0.67192747 | 0.00019814 |
| 426630 | *AGBL2* | -2.3659319 | 0.00016309 |
| 421194 | *AHSA2* | -0.8945527 | 0.0000684 |
| 421783 | *AIM1* | 1.88807823 | 0.00000547 |
| 430522 | *AK8* | -0.8979394 | 0.00680178 |
| 100858093 | *AKAP7L* | 1.51746234 | 0.00544149 |
| 395695 | *AKAP9* | -0.6392695 | 0.00520395 |
| 771077 | *AKD1* | -1.5288568 | 0.00019742 |
| 395844 | *ALDH1A2* | 1.13062212 | 0.00775069 |
| 421755 | *AMD1* | 0.94494826 | 0.00815639 |
| 417810 | *AMIGO2* | 1.12023472 | 0.00293935 |
| 416143 | *ANKHD1* | 0.89485988 | 0.00180886 |
| 422651 | *ANKRD17* | 0.55820484 | 0.00836437 |
| 415766 | *ANKRD27* | 0.68039112 | 0.0010282 |
| 420640 | *ANKRD28* | -0.5203204 | 0.00280223 |
| 422968 | *ANO3* | 1.01768693 | 0.00346714 |
| 423637 | *ANXA11* | -0.5101722 | 0.00850092 |
| 420149 | *AP1M1* | 0.58380678 | 0.00425282 |
| 417645 | *APPBP2* | 0.85429417 | 5.53E-07 |
| 428752 | *AREG* | 1.50009492 | 0.00101966 |
| 423289 | *ARHGAP11A* | 0.91276974 | 0.00109821 |
| 429163 | *ARHGEF26* | -0.4686067 | 0.00358763 |
| 428744 | *ARHGEF38* | 1.88002842 | 0.00914776 |
| 418761 | *ARHGEF7* | 0.76554991 | 0.00016953 |
| 426495 | *ARL2BP* | 0.68771217 | 0.00520395 |
| 428459 | *ARPP21* | 0.94030944 | 0.0033604 |
| 427772 | *ARRDC1* | 0.7652075 | 0.00514231 |
| 395119 | *ARVCF* | 1.19549759 | 0.00298888 |

| | | | |
|---|---|---|---|
| 422572 | *ASB5* | 0.62432081 | 0.00891337 |
| 424354 | *ASPM* | 1.0758644 | 0.00281199 |
| 417185 | *ASS1* | -1.8263922 | 0.0000917 |
| 418213 | *ASUN* | -0.7306758 | 0.00091092 |
| 421993 | *ASXL2* | -0.7999453 | 0.00329193 |
| 417966 | *ATF7IP* | 0.89565131 | 0.00136414 |
| 422254 | *ATP11C* | -0.8151709 | 0.0005731 |
| 419866 | *ATP5F1* | 0.81214452 | 0.00366657 |
| 395821 | *ATP6V1A* | 0.5521733 | 0.00254896 |
| 422776 | *ATP8A1* | 0.7019133 | 0.00178132 |
| 415949 | *ATRIP* | 0.94188418 | 0.00018894 |
| 416078 | *ATXN7* | 1.09917356 | 0.00046782 |
| 428167 | *AURKA* | 1.04459211 | 0.00354439 |
| 420750 | *AVL9* | 0.54970438 | 0.00437512 |
| 396121 | *B4GALT1* | -0.8245629 | 0.00584057 |
| 421875 | *BAI3* | 0.99251609 | 0.00132658 |
| 107053959 | *BARHL2* | 2.28522753 | 0.00263348 |
| 427819 | *BAZ1B* | 1.02914785 | 0.00032031 |
| 420745 | *BBS9* | 0.77990924 | 0.00282392 |
| 396056 | *BFSP1* | 1.3430809 | 0.00084734 |
| 395165 | *BHLHE23* | 1.8727893 | 0.00136645 |
| 416371 | *BHMT* | 1.203604 | 0.00343677 |
| 415577 | *BLM* | 0.76124263 | 0.00670302 |
| 395996 | *BLMH* | 0.56210658 | 0.00548396 |
| 420744 | *BMPER* | -1.1817173 | 0.00502134 |
| 427504 | *BNC1* | 1.48954399 | 0.00042566 |
| 415379 | *BNIP2* | 0.7530432 | 0.00112613 |
| 424061 | *BOLL* | 1.25769549 | 0.00204206 |
| 395882 | *BPIFB2* | 3.34895207 | 0.0037328 |
| 373983 | *BRCA1* | 0.81841834 | 0.00933488 |
| 374139 | *BRCA2* | 1.36100827 | 0.00085239 |
| 424506 | *BRDT* | 1.69626229 | 0.00021406 |
| 417642 | *BRIP1* | 0.72795837 | 0.00720197 |
| 423098 | *BRSK2* | 1.05120183 | 0.00054172 |
| 423029 | *BTBD10* | 1.19599505 | 0.0000178 |
| 101752037 | *BTBD6* | 1.00268226 | 0.00025841 |
| 428248 | *BUD13* | 1.09036443 | 0.00032031 |
| 101747767 | *C10H15ORF61* | 1.16906942 | 0.00166004 |
| 416292 | *C10ORF10* | 0.71342721 | 0.00997681 |
| 428970 | *C10ORF12* | 1.15551674 | 0.00366657 |
| 423053 | *C11ORF16* | -1.5893116 | 0.00095481 |
| 423113 | *C11ORF24* | 1.4093318 | 0.00018799 |
| 419090 | *C11ORF30* | 0.66256994 | 0.00420064 |
| 416070 | *C12H3ORF67* | 1.12432919 | 0.00017112 |

| | | | |
|---|---|---|---|
| 423524 | *C14ORF39* | 2.08560642 | 0.00017428 |
| 772069 | *C17H9ORF9* | -1.3989959 | 0.00070041 |
| 417154 | *C17H9ORF96* | 1.76400086 | 0.00478254 |
| 772176 | *C17ORF64* | 3.57888373 | 0.00047172 |
| 418068 | *C1H12ORF23* | 0.99907989 | 0.0000337 |
| 419041 | *C1H12ORF4* | 0.7786006 | 0.00082039 |
| 101748677 | *C1H12ORF40* | 2.24860432 | 0.00275124 |
| 417921 | *C1H12ORF63* | -1.2006948 | 0.0065254 |
| 395489 | *C1H21ORF91* | 0.82703325 | 0.00088566 |
| 770460 | *C1ORF146* | 1.97917989 | 0.00042078 |
| 395744 | *C1ORF158* | 3.95422438 | 0.0000271 |
| 419427 | *C21H1ORF159* | 0.92733719 | 0.00115926 |
| 101750708 | *C26H6ORF132* | 1.64983542 | 0.00032584 |
| 420625 | *C2H7ORF31* | 1.9964536 | 0.00103592 |
| 428440 | *C3ORF48* | -1.9196453 | 0.00010682 |
| 422431 | *C4H4ORF27* | 1.35981562 | 0.00020175 |
| 428893 | *C5H14ORF166B* | 2.43735845 | 0.00385558 |
| 423498 | *C5H14ORF79* | 0.96302195 | 0.00642402 |
| 430674 | *C6H10ORF137* | 0.6239831 | 0.00091092 |
| 421712 | *C6ORF58* | 1.67359104 | 0.00477535 |
| 424040 | *C7H21ORF58* | -1.8695492 | 0.0063964 |
| 416658 | *C7ORF26* | 0.89595138 | 0.00026804 |
| 100857655 | *C7ORF72* | 3.33451181 | 0.00039543 |
| 424399 | *C8H1ORF112* | 0.96604338 | 0.00042988 |
| 420220 | *C8ORF88* | 1.39511362 | 0.00127854 |
| 427402 | *C9ORF24* | 1.63717299 | 0.00035438 |
| 417647 | *CA4* | -3.4048422 | 3.89E-07 |
| 770178 | *CABP1* | 1.20236878 | 0.00366657 |
| 416526 | *CACNA1H* | 1.06250731 | 0.00509724 |
| 427900 | *CACNA1I* | 2.17786658 | 0.0000074 |
| 428425 | *CALCR* | 1.5763007 | 0.00738208 |
| 427515 | *CALML4* | -2.2683024 | 0.00237845 |
| 723973 | *CAMK4* | 1.35108909 | 0.00016309 |
| 417127 | *CAMSAP1* | 0.67360003 | 0.00696369 |
| 422341 | *CAPN6* | 0.77862732 | 0.00187433 |
| 416476 | *CARD11* | -0.935615 | 0.00575582 |
| 418757 | *CARS2* | 1.09589586 | 0.00644019 |
| 374038 | *CBL* | 0.73216223 | 0.00432627 |
| 424206 | *CCDC108* | 5.24827712 | 0.00011311 |
| 416917 | *CCDC117* | -0.4732529 | 0.0080679 |
| 417723 | *CCDC146* | -1.931957 | 0.00000668 |
| 424601 | *CCDC17* | -2.6043593 | 0.00052257 |
| 421609 | *CCDC34* | 1.25767123 | 0.00028492 |
| 427589 | *CCDC37* | -1.828804 | 0.00986237 |

| | | | |
|---|---|---|---|
| 419958 | *CCDC47* | 0.75707281 | 0.00298888 |
| 423666 | *CCDC6* | 0.80418267 | 0.00079769 |
| 416873 | *CCDC63* | 2.17732271 | 0.00402357 |
| 769634 | *CCDC78* | -1.2057707 | 0.00881307 |
| 424277 | *CCDC93* | 0.67038996 | 0.00610169 |
| 416161 | *CCNG1* | 0.78986453 | 0.00288063 |
| 423821 | *CCNJ* | 0.92567907 | 0.00042988 |
| 423446 | *CCNK* | 1.11162174 | 0.00666272 |
| 423606 | *CCSER2* | 0.63137339 | 0.00102152 |
| 419406 | *CDC2L1* | 0.46009976 | 0.00534907 |
| 416746 | *CDC42BPA* | -1.1476343 | 0.0064124 |
| 424658 | *CDCP2* | 1.39257877 | 0.00032031 |
| 420912 | *CDH12* | 1.29634625 | 0.00218874 |
| 416223 | *CDHR2* | -3.4706078 | 0.00000668 |
| 427846 | *CDHR3* | -2.2150922 | 0.0025701 |
| 428306 | *CDK12* | 1.03169092 | 0.0020865 |
| 423575 | *CDKL1* | -1.2311117 | 0.00711884 |
| 395320 | *CDX4* | -2.49013 | 0.00180036 |
| 107052707 | *CEBPD* | -1.4291427 | 0.00070767 |
| 373923 | *CELF1* | -0.5934075 | 0.00621212 |
| 416420 | *CEND1* | 0.7895053 | 0.00510179 |
| 770044 | *CEND1* | 0.94712311 | 0.00163213 |
| 424260 | *CEND1* | 1.08086847 | 0.00040311 |
| 395922 | *CENPC* | 0.56166713 | 0.00352901 |
| 426563 | *CENPL* | 0.83125056 | 0.00750585 |
| 693246 | *CENPN* | 1.0166431 | 0.0025421 |
| 421716 | *CENPW* | 1.17867123 | 0.00091066 |
| 100859489 | *CEP126* | -1.64178 | 0.00930459 |
| 419547 | *CEP85* | 1.21350498 | 0.00038327 |
| 426648 | *CERS3* | 0.75616725 | 0.00078689 |
| 100859681 | *CFAP126* | -1.5993702 | 0.00746101 |
| 100857593 | *CFD* | -1.4750319 | 0.00446053 |
| 770709 | *CHCHD7* | 0.8474153 | 0.00510179 |
| 421312 | *CHGB* | 1.34290511 | 0.00741637 |
| 422318 | *CHIC1* | 0.97162417 | 0.00140342 |
| 419009 | *CHORDC1* | -0.8281045 | 0.0000243 |
| 395608 | *CHRNA1* | 1.45566403 | 0.00972207 |
| 386578 | *CHRNA3* | 2.35809618 | 0.00272087 |
| 395606 | *CHRNA4* | 1.25961201 | 0.00613257 |
| 768659 | *CIDEA* | 1.03621545 | 0.00216946 |
| 425789 | *CIRBP* | 0.58330086 | 0.00706931 |
| 374002 | *CKMT1A* | -2.0798923 | 0.0001384 |
| 416240 | *CLINT1* | 1.01333685 | 0.00017434 |
| 100858576 | *CLK1* | -0.6223469 | 0.0062821 |

| | | | |
|---|---|---|---|
| 428223 | *CLSPN* | 1.18391888 | 0.00063552 |
| 770023 | *CMC2* | 0.74567969 | 0.00381898 |
| 427537 | *CMTR2* | 1.04529811 | 0.00023 |
| 421013 | *CNDP2* | 0.88966939 | 0.0012295 |
| 428975 | *CNNM2* | 1.04546404 | 0.00039468 |
| 420669 | *CNOT10* | 0.8657343 | 0.0000454 |
| 417936 | *CNOT4* | 1.25396856 | 0.00027513 |
| 428633 | *CNR1* | 1.08040432 | 0.00036063 |
| 396149 | *CNTN4* | 0.89661458 | 0.0037643 |
| 430532 | *CNTNAP4* | 1.44179781 | 0.0048307 |
| 395779 | *COCH* | -1.7545005 | 0.00237845 |
| 395875 | *COL12A1* | -0.7509508 | 0.00584057 |
| 396243 | *COL1A2* | -1.1057355 | 0.00606763 |
| 769778 | *COMMD6* | 0.70610381 | 0.00173803 |
| 416683 | *CPA5* | 1.60399545 | 0.0018998 |
| 422832 | *CPEB2* | 1.17023116 | 0.00644019 |
| 419853 | *CR1L* | -1.0917123 | 0.00024675 |
| 404297 | *CRH* | -2.0917386 | 0.00070767 |
| 428191 | *CROCC* | -1.2589049 | 0.00639201 |
| 426184 | *CSAD* | -1.577382 | 0.0024623 |
| 421899 | *CSMD1* | 1.07994685 | 0.00070041 |
| 772072 | *CTD-2510F5.6* | 0.87208196 | 0.00275124 |
| 428852 | *CTNND1* | -0.6421879 | 0.00243611 |
| 428163 | *CTSA* | -1.1511895 | 0.00623472 |
| 420467 | *CUL2* | 0.91906926 | 0.0001782 |
| 100858121 | *CXORF23* | 0.97323051 | 0.00216438 |
| 408182 | *CYB5R2* | -1.0681552 | 0.00028645 |
| 421837 | *CYB5R4* | 0.95871004 | 0.00156329 |
| 107051085 | *CYHR1* | 1.11348399 | 0.00980138 |
| 414838 | *CYP11A1* | -1.3699881 | 0.00988931 |
| 420548 | *CYP51A1* | 1.24539208 | 0.00217951 |
| 429052 | *CYTIP* | -4.9874767 | 0.00000255 |
| 374083 | *DAB1* | 0.88893933 | 0.00978418 |
| 723789 | *DACT1* | -0.9284796 | 0.00051724 |
| 421561 | *DACT2* | 1.26937216 | 0.0000415 |
| 420536 | *DBF4* | 0.89807528 | 0.0022047 |
| 420821 | *DCDC2* | -3.4058799 | 0.00000218 |
| 426942 | *DDA1* | 0.75189908 | 0.0000074 |
| 423429 | *DDX24* | 0.97847059 | 0.00842272 |
| 421327 | *DEGS1* | 0.68284246 | 0.00854962 |
| 424710 | *DEPDC1* | 1.08854507 | 0.00888929 |
| 772367 | *DGKB* | 1.08287468 | 0.00078689 |
| 415876 | *DHODH* | -1.389701 | 0.00190129 |
| 417499 | *DHX33* | -0.7100234 | 0.0001586 |

| | | | |
|---|---|---|---|
| 419798 | *DIXDC1* | 1.21400824 | 0.00021191 |
| 421903 | *DLGAP2* | 1.84280268 | 0.00253208 |
| 429107 | *DMRTB1* | 1.88923488 | 0.00227169 |
| 416818 | *DNAH10* | -1.2665735 | 0.00130353 |
| 420921 | *DNAH5* | -1.6421315 | 0.00111343 |
| 417314 | *DNAH9* | -1.6408487 | 0.00070041 |
| 417453 | *DNAI2* | -1.6717065 | 0.0020826 |
| 415360 | *DNAJA4* | -0.8044262 | 0.00088374 |
| 419060 | *DNAJB13* | -0.7510034 | 0.00492566 |
| 424550 | *DNAJB4* | -0.7280703 | 0.00358647 |
| 427409 | *DNAJB5* | 1.69713121 | 0.00358647 |
| 770080 | *DNAJB8* | 3.05904785 | 0.00090185 |
| 424698 | *DNAJC6* | 1.37602241 | 0.00019477 |
| 423640 | *DNAJC9* | 0.95573364 | 0.00016309 |
| 417196 | *DOLPP1* | 1.25911288 | 0.00107753 |
| 420266 | *DPYS* | -2.1781126 | 0.0011367 |
| 395155 | *DPYSL2* | 0.90432044 | 0.00513672 |
| 428527 | *DSC2* | -1.3957879 | 0.00550162 |
| 421104 | *DTNA* | 0.72445365 | 0.00126778 |
| 421340 | *DUSP10* | 0.8580734 | 0.00107753 |
| 417657 | *DUSP14* | 0.56883117 | 0.00933488 |
| 769430 | *E2F4* | 0.8615012 | 0.00334457 |
| 100858704 | *ECM1* | -1.7346103 | 0.00039543 |
| 415294 | *EDC3* | 0.63385906 | 0.00760581 |
| 416021 | *EEFSEC* | 0.88056714 | 0.00042078 |
| 423106 | *EFCAB4B* | 1.118119 | 0.00730641 |
| 417587 | *EFCAB5* | -0.9242359 | 0.00621623 |
| 395896 | *EFNB1* | -0.8048483 | 0.00891231 |
| 423316 | *EGLN3* | 1.29504557 | 0.00281102 |
| 395514 | *EIF5B* | 0.71375924 | 0.00011311 |
| 770158 | *ELAVL2* | 1.24814909 | 0.0000462 |
| 395634 | *ELAVL4* | 1.11118563 | 0.00017428 |
| 419832 | *ELK4* | -0.6513468 | 0.00694932 |
| 420124 | *ELL* | 1.12047261 | 0.00282207 |
| 418971 | *ELMOD1* | 1.14123889 | 0.00311056 |
| 420858 | *ELOVL2* | 1.15323646 | 0.00854962 |
| 772178 | *EMCN* | 0.87587712 | 0.00281102 |
| 374180 | *ENAH* | 0.61785151 | 0.00160225 |
| 421447 | *EPHX1L* | -1.087551 | 0.00598425 |
| 418837 | *EPSTI1* | 1.61363896 | 0.00090474 |
| 769533 | *ERC2* | 1.14455716 | 0.00022838 |
| 418847 | *ERICH6B* | 1.84613483 | 0.00935213 |
| 395575 | *ESR2* | 1.31105128 | 0.00857552 |
| 770095 | *EVC2* | 0.86308381 | 0.00011834 |

| | | | |
|---|---|---|---|
| 419454 | *EXOSC10* | 0.57624472 | 0.00510179 |
| 619530 | *EXOSC9* | 0.35796566 | 0.00881307 |
| 374165 | *FABP4* | -1.7068116 | 0.00694895 |
| 416154 | *FABP6* | -1.7030992 | 0.00018444 |
| 421552 | *FAM120B* | -1.3625073 | 0.00363611 |
| 422237 | *FAM122A* | -0.5688796 | 0.00298192 |
| 422515 | *FAM13A* | -2.2339369 | 0.00000056 |
| 423602 | *FAM13C* | -1.1199462 | 0.00682632 |
| 769188 | *FAM161A* | -1.2204359 | 0.00050968 |
| 431595 | *FAM169A* | 0.73846827 | 0.00285265 |
| 418395 | *FAM172BP* | 1.01315942 | 0.0079328 |
| 421727 | *FAM184A* | 1.25169326 | 0.00011367 |
| 100859145 | *FAM184B* | 1.4551613 | 0.0003587 |
| 421045 | *FAM210A* | 1.59738585 | 0.0000329 |
| 768548 | *FAM222A* | -1.1819819 | 0.00167716 |
| 421744 | *FAM26F* | -0.8515118 | 0.00928035 |
| 426544 | *FAM46C* | 1.25355672 | 0.00018272 |
| 378905 | *FAM53A* | -0.6668924 | 0.00179198 |
| 768603 | *FAM53B* | -0.6287505 | 0.00477535 |
| 428169 | *FAM65C* | 1.5649824 | 0.00020931 |
| 423282 | *FAM71BL* | 3.5685723 | 0.000014 |
| 415396 | *FAM81A* | -1.3581373 | 0.00584057 |
| 770521 | *FAM83A* | 1.65958309 | 0.00111456 |
| 421289 | *FBXO11* | 0.77763811 | 0.0001333 |
| 421906 | *FBXO25* | 0.90349607 | 0.00105026 |
| 776097 | *FBXO39* | 1.48282929 | 0.00989473 |
| 420245 | *FBXO43* | 1.35010877 | 0.00310208 |
| 424904 | *FCGBP* | -3.369307 | 3.89E-07 |
| 395704 | *FGF12* | 1.54467718 | 0.00020759 |
| 414831 | *FGF13* | 1.18420366 | 0.00048514 |
| 395453 | *FGF18* | 1.19084625 | 0.00658155 |
| 770457 | *FGF5* | 1.43827149 | 0.00446053 |
| 422484 | *FHDC1* | 1.46197958 | 9.73E-07 |
| 428618 | *FLRT1* | -2.4127211 | 0.00076537 |
| 775973 | *FMN2* | 1.36224602 | 0.0001514 |
| 100857760 | *FOXF2* | -3.0466362 | 0.00097094 |
| 422726 | *FRG1* | 0.81851386 | 0.00644019 |
| 427452 | *FRMD3* | 1.27257932 | 0.00143197 |
| 429082 | *FRRS1* | 0.99484201 | 0.00621212 |
| 395163 | *FSTL4* | 0.80581399 | 0.00918418 |
| 395726 | *FTCD* | -2.0626314 | 0.0004348 |
| 428717 | *GAB3* | 1.09671246 | 0.00102152 |
| 422289 | *GABRA3* | 1.6076802 | 0.00738786 |
| 418684 | *GABRA5* | 1.97479989 | 0.00019126 |

| | | | |
|---|---|---|---|
| 396289 | *GABRG2* | 0.99309513 | 0.00933488 |
| 416174 | *GABRP* | -1.4439306 | 0.00618181 |
| 395743 | *GAD1* | 1.13158747 | 0.00972207 |
| 417006 | *GAL3ST1* | -2.7090855 | 0.00023 |
| 424854 | *GAL3ST2* | -1.8297104 | 0.0084248 |
| 420978 | *GALNT12* | 0.60633884 | 0.00916381 |
| 416796 | *GALNT9* | 1.23990267 | 0.00100856 |
| 395950 | *GBX2* | 3.20940154 | 0.0000462 |
| 396196 | *GCG* | -2.0890053 | 0.00803625 |
| 428478 | *GCM2* | 1.70324487 | 0.0000833 |
| 419201 | *GDAP1L1* | 1.24280287 | 0.00055707 |
| 417631 | *GDPD1* | -0.7271902 | 0.00282392 |
| 419070 | *GDPD5* | 0.78148017 | 0.00293111 |
| 404771 | *GEM* | -1.0075403 | 0.00802093 |
| 395994 | *GFRA1* | 0.92923901 | 0.00813473 |
| 422121 | *GGA3* | 0.58512957 | 0.00098467 |
| 424743 | *GIGYF2* | 0.81956058 | 0.0000169 |
| 101750732 | *GIN1* | 0.79660433 | 0.00118773 |
| 415824 | *GINS2* | 1.03298417 | 0.00678273 |
| 424779 | *GK5* | -0.6997953 | 0.00721803 |
| 422411 | *GLRB* | 1.23406431 | 0.00520395 |
| 416807 | *GLT1D1* | 1.05235084 | 0.00025599 |
| 429534 | *GLTP* | 0.91309876 | 0.00690802 |
| 423560 | *GMFB* | 0.91583143 | 0.00591388 |
| 421004 | *GMNN* | 0.93954305 | 0.00329906 |
| 415698 | *GNAO1* | 1.00636153 | 0.00661233 |
| 770226 | *GNAZ* | 1.18501051 | 0.00048821 |
| 415417 | *GNB5* | -0.7653867 | 0.00410475 |
| 422772 | *GNPDA2* | 1.41043612 | 0.00223719 |
| 424156 | *GORASP2* | 1.07362896 | 0.00014942 |
| 418445 | *GPA33* | -3.5664933 | 0.00010761 |
| 421355 | *GPATCH2* | 0.56507926 | 0.00625321 |
| 418795 | *GPC5* | 2.31449901 | 0.0054496 |
| 427956 | *GPR15* | 1.77498248 | 0.00672834 |
| 100857561 | *GPR63* | 0.77968793 | 0.00436969 |
| 395648 | *GPRIN2* | 1.26091885 | 0.00025731 |
| 429506 | *GRHL2* | -1.266038 | 0.00895504 |
| 428934 | *GRID1* | 1.13097964 | 0.00021406 |
| 422520 | *GRID2* | 1.37865061 | 0.0000107 |
| 428628 | *GRIK2* | 0.89964159 | 0.00514231 |
| 419619 | *GRIK3* | -1.2514842 | 0.00813473 |
| 416112 | *GRM7* | 1.58721363 | 0.00078574 |
| 418335 | *GSK3B* | 0.86985082 | 0.00916047 |
| 417487 | *GTF2IRD1* | 1.14160994 | 0.00260251 |

| | | | |
|---|---|---|---|
| 425204 | *GTSE1* | 1.16480687 | 0.00550162 |
| 423898 | *HABP2* | 1.37582034 | 0.00930459 |
| 378924 | *HAL* | 1.71267475 | 0.00162415 |
| 424482 | *HCCS* | -0.8979988 | 0.00227169 |
| 426867 | *HCLS1* | -5.3509489 | 0.0000394 |
| 424050 | *HECW2* | 1.04494669 | 0.00045043 |
| 429055 | *HENMT1* | 1.6710635 | 0.00090647 |
| 428365 | *HEY1* | 0.99472886 | 0.00836437 |
| 421597 | *HIPK3* | -0.988415 | 0.00088566 |
| 428208 | *HIVEP3* | 0.86780256 | 0.00760581 |
| 768421 | *HK3* | 1.82343355 | 0.00466832 |
| 378795 | *HMGA1* | -1.0920992 | 0.00232028 |
| 416163 | *HMMR* | 1.09237971 | 0.00518513 |
| 423079 | *HPS5* | -2.8005589 | 5.05E-05 |
| 427474 | *HSD17B3* | -1.8824338 | 0.00689903 |
| 423463 | *HSP90AA1* | -1.5061463 | 0.00025841 |
| 396188 | *HSP90AB1* | -0.9985643 | 0.00010143 |
| 395853 | *HSPA8* | -1.5480919 | 2.48E-08 |
| 427963 | *HTR1F* | 1.16448811 | 0.0069559 |
| 771148 | *IFITM10* | 1.7179105 | 0.00132159 |
| 416123 | *IFT122* | -0.6381615 | 0.004544 |
| 395953 | *IGF2BP1* | 3.510121 | 0.00302362 |
| 420617 | *IGF2BP3* | 1.63626821 | 0.00092697 |
| 416133 | *IK* | 0.68457995 | 0.00520395 |
| 404671 | *IL12B* | 2.4783392 | 0.00015402 |
| 422219 | *IL13RA2* | -1.9714187 | 0.00121129 |
| 416585 | *IL4R* | -0.7057759 | 0.00220843 |
| 424682 | *INADL* | 0.72662318 | 0.00312433 |
| 424852 | *ING5* | 0.68362904 | 0.00217951 |
| 418690 | *INPP4A* | -0.5721813 | 0.00527637 |
| 422454 | *INPP4B* | 0.79148212 | 0.00718063 |
| 423067 | *INSC* | 1.42598593 | 0.00639545 |
| 422537 | *INTS12* | 1.0692235 | 0.00024675 |
| 421374 | *INTS7* | -1.0294885 | 0.00272457 |
| 415554 | *IQCHL* | 1.31097971 | 0.00011311 |
| 418476 | *JAM2* | 1.16126153 | 0.00143654 |
| 418023 | *JOSD1* | 0.68507684 | 0.00140472 |
| 420190 | *JPH1* | 0.89440673 | 0.00151666 |
| 423734 | *KAT6B* | 0.77551285 | 0.00081145 |
| 421626 | *KATNA1* | 0.74750845 | 0.00488785 |
| 426211 | *KAZN* | 1.00184964 | 0.00292612 |
| 420749 | *KBTBD2* | -0.6105756 | 0.00534429 |
| 416085 | *KBTBD8* | 0.66781602 | 0.00133506 |
| 395730 | *KCNAB1* | 1.45324217 | 0.00485472 |

| | | | |
|---|---|---|---|
| 428105 | *KCNE3* | -1.562617 | 0.00514231 |
| 424184 | *KCNH7* | 1.16859668 | 0.00892788 |
| 428902 | *KCNK10* | 2.36142756 | 0.00417612 |
| 772022 | *KCNK16* | -1.6785135 | 0.00317695 |
| 419814 | *KCTD20* | 0.75815313 | 0.00192877 |
| 419526 | *KCTD9* | 0.94060478 | 0.00046782 |
| 101750381 | *KIAA0408* | 1.11320911 | 0.00107715 |
| 415485 | *KIAA1024* | 1.11500781 | 0.00678273 |
| 421598 | *KIAA1549L* | 1.52610907 | 0.0000377 |
| 100857276 | *KIAA1644* | 0.9788763 | 0.00065202 |
| 424141 | *KIAA1715* | 1.22762146 | 0.00021019 |
| 421196 | *KIAA1841* | 1.10003523 | 0.00030459 |
| 421260 | *KIF16B* | 0.50598481 | 0.00603477 |
| 416220 | *KIF20A* | 0.93350684 | 0.00918418 |
| 423489 | *KIF26A* | 0.94982367 | 0.0024623 |
| 416332 | *KIF3A* | 0.88946883 | 0.0000223 |
| 429432 | *KIF3C* | -1.059102 | 0.00010143 |
| 420033 | *KLHL10* | 2.03607677 | 0.00039543 |
| 418840 | *LACC1* | -1.4749923 | 0.00083975 |
| 374016 | *LAMA1* | 1.84130015 | 0.00047331 |
| 420457 | *LARP4B* | 0.81911738 | 0.00417612 |
| 420002 | *LASP1* | -0.6727801 | 0.00933585 |
| 423610 | *LDB3* | 1.54079886 | 0.00053332 |
| 425107 | *LGALS2* | -3.6815575 | 3.89E-07 |
| 423802 | *LGI1* | 1.16385279 | 0.0000425 |
| 428605 | *LGR4* | -0.6811233 | 0.00829394 |
| 396397 | *LHX9* | 1.41450647 | 0.00672834 |
| 422595 | *LIN54* | 0.88586821 | 0.00042566 |
| 415464 | *LMNA* | 1.58998834 | 9.73E-07 |
| 396223 | *LMNB1* | 0.78854808 | 0.00243611 |
| 418179 | *LMO3* | 0.75573236 | 0.00690802 |
| 100857445 | *LOC100857445* | 0.66686967 | 0.00034359 |
| 100857927 | *LOC100857927* | 1.83552951 | 0.00073821 |
| 100858301 | *LOC100858301* | 1.17475651 | 0.00930459 |
| 100858336 | *LOC100858336* | 0.93067104 | 0.00080896 |
| 100858845 | *LOC100858845* | 3.29995939 | 0.00018522 |
| 100859230 | *LOC100859230* | 1.36721475 | 0.00586708 |
| 100859272 | *LOC100859272* | -2.1509118 | 0.0001333 |
| 100859371 | *LOC100859371* | 1.54051453 | 0.00065556 |
| 100859381 | *LOC100859381* | 4.93832142 | 0.0000073 |
| 100859449 | *LOC100859449* | 5.5606747 | 0.001225 |
| 101747255 | *LOC101747255* | 3.41047737 | 0.00050829 |
| 101747522 | *LOC101747522* | -1.723058 | 0.0058422 |
| 101747844 | *LOC101747844* | -5.2524349 | 5.42E-12 |

| | | | |
|---|---|---|---|
| 101748683 | *LOC101748683* | 3.63449859 | 0.00025731 |
| 426430 | *LOC101749001* | -0.9554347 | 0.00027513 |
| 101749216 | *LOC101749216* | -3.6160107 | 0.00025731 |
| 101749269 | *LOC101749269* | 2.39418078 | 0.00389369 |
| 395389 | *LOC101749492* | -1.3236286 | 0.0000933 |
| 101750367 | *LOC101750367* | -4.1289087 | 9.83E-08 |
| 101750448 | *LOC101750448* | -2.0497965 | 0.00464432 |
| 101750583 | *LOC101750583* | 0.82706853 | 0.00935213 |
| 101750794 | *LOC101750794* | 2.29066702 | 0.0072288 |
| 101751597 | *LOC101751597* | -1.5293132 | 0.00054914 |
| 101752063 | *LOC101752063* | 4.25218593 | 0.0000768 |
| 107049002 | *LOC107049002* | -0.8778689 | 0.00520395 |
| 107049275 | *LOC107049275* | 1.42915751 | 0.00989473 |
| 107049666 | *LOC107049666* | -2.3084318 | 0.00490339 |
| 107049720 | *LOC107049720* | -2.7507272 | 0.00437232 |
| 107049863 | *LOC107049863* | -0.6505669 | 0.00888929 |
| 107051352 | *LOC107051352* | 2.26607445 | 0.00904711 |
| 107051435 | *LOC107051435* | -2.5678777 | 0.00088566 |
| 107052033 | *LOC107052033* | -1.1441953 | 0.00989473 |
| 107052453 | *LOC107052453* | -1.3797512 | 0.00282392 |
| 107052725 | *LOC107052725* | -2.6444835 | 0.00020759 |
| 107052834 | *LOC107052834* | 1.85080474 | 0.00119916 |
| 107053055 | *LOC107053055* | 1.80487813 | 0.00010143 |
| 107053196 | *LOC107053196* | 1.44692969 | 0.00160105 |
| 107055024 | *LOC107055024* | -1.252647 | 0.00028492 |
| 107055038 | *LOC107055038* | 1.16018168 | 0.00918418 |
| 107055618 | *LOC107055618* | -3.7843084 | 0.000089 |
| 107056420 | *LOC107056420* | -3.4729335 | 0.00024999 |
| 107057502 | *LOC107057502* | -1.9521197 | 0.00829016 |
| 395647 | *LOC395647* | 1.20388388 | 0.00202964 |
| 396380 | *LOC396380* | 1.23760683 | 0.0019084 |
| 415780 | *LOC415780* | 1.0593428 | 0.00977608 |
| 417131 | *LOC417131* | 1.22200126 | 0.00809507 |
| 417345 | *LOC417345* | -1.2691474 | 0.00591181 |
| 418189 | *LOC418189* | 1.40543386 | 0.00101966 |
| 418356 | *LOC418356* | -3.3691982 | 0.00021091 |
| 421054 | *LOC421054* | -0.8442857 | 0.00132338 |
| 421195 | *LOC421195* | -2.6765397 | 0.0000439 |
| 421690 | *LOC421690* | -1.2937921 | 0.00120492 |
| 422301 | *LOC422301* | -1.1905773 | 0.00666269 |
| 422319 | *LOC422319* | 0.92850239 | 0.00273449 |
| 422372 | *LOC422372* | 0.65395073 | 0.00042078 |
| 423793 | *LOC423793* | 1.49014638 | 0.00094744 |
| 424028 | *LOC424028* | -2.8159643 | 0.00045481 |

| | | | |
|---|---|---|---|
| 424033 | *LOC424033* | 2.15175282 | 0.00016036 |
| 424167 | *LOC424167* | 1.79666009 | 0.00080857 |
| 424473 | *LOC424473* | 1.28273597 | 0.00626456 |
| 424892 | *LOC424892* | 1.19870896 | 0.0001586 |
| 424917 | *LOC424917* | -2.3682192 | 0.0048307 |
| 425431 | *LOC425431* | -1.2383327 | 0.00023515 |
| 426093 | *LOC426093* | -2.0494561 | 0.00293111 |
| 426385 | *LOC426385* | 1.3199822 | 0.00572575 |
| 427491 | *LOC427491* | 0.91803646 | 0.00619939 |
| 428510 | *LOC428510* | -1.9742398 | 0.00280223 |
| 430303 | *LOC430303* | 1.52569956 | 0.00412888 |
| 769052 | *LOC769052* | 1.60008055 | 0.00061351 |
| 770248 | *LOC770248* | -0.6303172 | 0.00738606 |
| 771537 | *LOC771537* | 0.58619855 | 0.00310208 |
| 771735 | *LOC771735* | -1.631203 | 0.00429891 |
| 772158 | *LOC772158* | -2.3591468 | 0.00133908 |
| 422742 | *LONRF1* | 0.69848867 | 0.00400354 |
| 420974 | *LPCAT1* | 0.53973972 | 0.00302362 |
| 424477 | *LPPR5* | 1.03777063 | 0.00534907 |
| 424713 | *LRRC40* | 0.78876498 | 0.00078689 |
| 420324 | *LRRC6* | -1.4568248 | 0.00237214 |
| 422761 | *LRRC66* | -1.6938535 | 0.00065501 |
| 424711 | *LRRC7* | 1.07508931 | 0.00061536 |
| 417201 | *LRRC8A* | 0.68874604 | 0.00044091 |
| 424513 | *LRRC8D* | 0.83192948 | 0.00445052 |
| 420205 | *LRRCC1* | 0.70613789 | 0.00146923 |
| 428261 | *LRRN2* | 1.31720188 | 0.00618487 |
| 771330 | *LSM14B* | 0.44990143 | 0.00976346 |
| 423001 | *LUZP2* | 1.39971642 | 0.00078689 |
| 421514 | *LYST* | 0.6755877 | 0.00738208 |
| 418265 | *MAN1A2* | -0.6895504 | 0.00281102 |
| 426428 | *MANSC1* | 1.32636355 | 0.00547651 |
| 427144 | *MAP3K1* | -1.4065885 | 0.00028494 |
| 429048 | *MAP3K19* | -1.5030149 | 0.0047811 |
| 426292 | *MAPK15* | -0.9058971 | 0.00366657 |
| 420130 | *MAST3* | 1.03431209 | 0.0000275 |
| 420487 | *MASTL* | 1.21049794 | 0.0003587 |
| 395428 | *MBD4* | 0.7394344 | 0.00447067 |
| 422253 | *MCF2* | 0.7895931 | 0.00319303 |
| 418748 | *MCF2L* | 0.93839904 | 0.00053491 |
| 424959 | *MCF2L2* | 1.80403949 | 0.0000747 |
| 428942 | *MCU* | 0.77553527 | 0.00065202 |
| 421431 | *MDGA1* | 2.04614925 | 0.00362532 |
| 417639 | *MED13* | -0.6199806 | 0.00678273 |

| | | | |
|---|---|---|---|
| 416941 | *MED15* | 1.04369933 | 0.00810465 |
| 419000 | *MED17* | 0.54080502 | 0.00350921 |
| 418859 | *MED4* | 0.79424701 | 0.00168429 |
| 415545 | *MEGF11* | 1.78088831 | 0.0000344 |
| 416539 | *MEIOB* | 1.34663563 | 0.00954031 |
| 425865 | *METTL22* | 1.03552543 | 0.00806367 |
| 100858555 | *MFAP2* | 1.52121649 | 0.00147641 |
| 396127 | *MFI2* | -1.3142866 | 0.00312154 |
| 423851 | *MGEA5* | 0.68686508 | 0.00451538 |
| 395912 | *MGP* | -2.3016794 | 0.00019126 |
| 418250 | *MKLN1* | 0.60522773 | 0.00026241 |
| 421564 | *MLLT4* | 0.77393934 | 0.00366657 |
| 418981 | *MMP10* | -2.0102724 | 0.0000127 |
| 395683 | *MMP13* | -2.50783 | 9.37E-05 |
| 395387 | *MMP9* | -2.4140317 | 1.31E-06 |
| 421799 | *MMS22L* | 0.67315658 | 0.00839752 |
| 427840 | *MNT* | 0.93984474 | 0.00051221 |
| 426437 | *MOGAT2* | -1.2969289 | 0.00091092 |
| 554283 | *MORF4L1* | 1.58962303 | 0.0000604 |
| 418441 | *MPZL1* | -0.8129135 | 0.00107753 |
| 420686 | *MRPL3* | 0.92577435 | 0.0000441 |
| 420196 | *MRPL53* | 1.36981452 | 0.00282392 |
| 424824 | *MRPS22* | 0.83566785 | 0.00460435 |
| 427318 | *MSH3* | 1.03196395 | 0.00534429 |
| 418641 | *MSL3* | 0.50135992 | 0.00850092 |
| 395245 | *MSX2* | -3.6467724 | 0.0000281 |
| 396212 | *MT4* | -1.1801718 | 0.00625321 |
| 415823 | *MTHFSD* | 0.70495487 | 0.0041647 |
| 418938 | *MTMR6* | 0.50827688 | 0.00375392 |
| 422033 | *MTMR9* | 0.64181439 | 0.00582726 |
| 422729 | *MTUS1* | 0.79879972 | 0.00513202 |
| 100859223 | *MUC13* | -3.6435966 | 7.42E-08 |
| 423101 | *MUC2* | -2.9276693 | 0.00010143 |
| 425514 | *MUM1* | 0.85562232 | 0.0000897 |
| 396258 | *MYBL2* | 1.40550025 | 0.00032399 |
| 420841 | *MYLIP* | -0.5572355 | 0.00516461 |
| 420893 | *MYLK4* | 2.21601553 | 0.00470192 |
| 396072 | *MYO1A* | -2.3909513 | 0.00352956 |
| 415398 | *MYO1E* | 1.11298283 | 0.00192688 |
| 424157 | *MYO3B* | 1.44450174 | 0.00297075 |
| 419690 | *MYOM3* | -1.0958759 | 0.00908098 |
| 424671 | *MYSM1* | 0.77532588 | 0.00331683 |
| 419249 | *MYT1* | 1.71764192 | 0.00052257 |
| 100859189 | *MZB1* | -2.6086005 | 0.00416696 |

| | | | |
|---|---|---|---|
| 418912 | *N4BP2L2* | 0.82323603 | 0.0000841 |
| 418832 | *NAA16* | 0.89949391 | 0.00842272 |
| 424046 | *NABP1* | 0.84123479 | 0.00079227 |
| 769997 | *NAF1* | 0.88532179 | 0.00957482 |
| 426856 | *NARS* | 0.50897985 | 0.00756217 |
| 420007 | *NBR1* | 0.74052564 | 0.00040311 |
| 427379 | *NCBP1* | 0.42851795 | 0.00477535 |
| 424283 | *NCKAP5* | 1.15353644 | 0.000089 |
| 772284 | *NDC1* | 0.66158943 | 0.00177008 |
| 395134 | *NDC80* | 0.96483216 | 0.0014749 |
| 418541 | *NDUFV3* | 0.93405676 | 0.00126778 |
| 770083 | *NECAP1* | 0.85001074 | 0.00204206 |
| 770845 | *NELL1* | 1.07846934 | 0.00018894 |
| 419824 | *NFASC* | -0.7873742 | 0.00416054 |
| 386574 | *NFKB2* | -0.7863521 | 0.00784643 |
| 418404 | *NFKBIZ* | -1.0940097 | 0.00133333 |
| 424744 | *NGEF* | -1.7024018 | 0.00042078 |
| 373910 | *NHLH1* | 1.3957709 | 0.00613257 |
| 415986 | *NINJ1* | -0.7221422 | 0.00282392 |
| 416245 | *NIPAL4* | 1.38558879 | 0.00826153 |
| 107049081 | *NKAIN1* | 1.06368222 | 0.00970744 |
| 422368 | *NKAP* | 1.02762226 | 0.0000394 |
| 107053698 | *NKX2-3* | -1.6528529 | 0.00261771 |
| 428461 | *NME8* | 3.58361256 | 0.0000462 |
| 404779 | *NOCT* | 0.79780717 | 0.00508279 |
| 769466 | *NOL4* | 1.20589456 | 0.00018894 |
| 415955 | *NOL8* | 1.13022633 | 0.00048035 |
| 423755 | *NOLC1* | 0.75499885 | 0.00385558 |
| 424087 | *NOP58* | 1.05878032 | 0.0011367 |
| 415316 | *NPTN* | -0.6060061 | 0.0049692 |
| 100859014 | *NR2F1B* | -1.0878935 | 0.00137307 |
| 395961 | *NR5A2* | -2.053025 | 0.000056 |
| 424635 | *NRD1* | 0.62109126 | 0.00183642 |
| 423404 | *NRDE2* | 1.13140622 | 0.00028219 |
| 420873 | *NRN1* | 2.47413209 | 0.00042986 |
| 395398 | *NRXN1* | 0.85523054 | 0.00137307 |
| 416036 | *NTN4L* | -1.292336 | 0.00160277 |
| 420433 | *NUB1* | 0.50927123 | 0.00738208 |
| 417897 | *NUDT4* | 0.86088032 | 0.00080911 |
| 418841 | *NUFIP1* | 0.68785562 | 0.0048307 |
| 768717 | *NUMA1* | -1.4252518 | 0.00552349 |
| 395639 | *NUMB* | 0.84892548 | 0.00048973 |
| 418937 | *NUP58* | 0.94634725 | 0.00048973 |
| 423213 | *NUSAP1* | 0.91083586 | 0.00697824 |

| | | | |
|---|---|---|---|
| 419916 | *OARD1* | 1.20450602 | 0.00593099 |
| 420257 | *ODF1* | 4.43707272 | 0.0000167 |
| 428582 | *OPN3* | 1.42465796 | 0.00119881 |
| 424640 | *ORC1* | 0.98642838 | 0.00753553 |
| 417714 | *ORC5* | 1.02101945 | 0.00023 |
| 423981 | *ORMDL1* | 0.6723986 | 0.00813473 |
| 421079 | *OSBPL1A* | 0.89718599 | 0.00966186 |
| 419227 | *OSBPL2* | 0.7709476 | 0.00941158 |
| 395735 | *OTC* | -2.2110311 | 0.00070041 |
| 420502 | *OTUD1* | -1.3811497 | 0.00000547 |
| 422463 | *OTUD4* | 1.06546516 | 0.00037714 |
| 415380 | *OTUD7A* | 0.63223969 | 0.00510179 |
| 769290 | *OVCH2* | -3.0857626 | 0.0000345 |
| 396151 | *OVST* | 5.37638121 | 8.18E-07 |
| 420270 | *OXR1* | 1.01114078 | 0.00000021 |
| 429367 | *P2RX6* | 1.12205905 | 0.00270758 |
| 416326 | *P4HA2* | 0.77578619 | 0.0017162 |
| 421576 | *PACRG* | -1.4664832 | 0.00090474 |
| 428171 | *PADI1* | 1.40672017 | 0.00699326 |
| 396534 | *PAICS* | 0.7607405 | 0.00447067 |
| 422342 | *PAK3* | 0.87580042 | 0.00256926 |
| 416734 | *PAK7* | 0.66358102 | 0.00313077 |
| 770425 | *PAPD7* | 0.56920695 | 0.00225265 |
| 423678 | *PAPSS2* | -1.1477891 | 0.00188052 |
| 418958 | *PARP4* | 1.08382775 | 0.00033445 |
| 418092 | *PARPBP* | 1.83907374 | 0.00088374 |
| 395319 | *PCLO* | 1.54933066 | 0.00032788 |
| 424116 | *PDE1A* | 1.7565255 | 0.00164968 |
| 422677 | *PDE5A* | -1.0044361 | 0.00063495 |
| 428372 | *PDP1* | 1.02876607 | 0.00851275 |
| 427550 | *PDP2* | 0.82493127 | 0.0062508 |
| 428984 | *PDZD8* | 1.35608613 | 0.00018272 |
| 421279 | *PELI1* | -0.712016 | 0.00472255 |
| 421854 | *PHIP* | -0.4594404 | 0.00878085 |
| 374241 | *PI15* | -2.1386452 | 0.00227169 |
| 374268 | *PIK3AP1* | 0.8454697 | 0.00857942 |
| 422044 | *PKHD1* | 1.28379116 | 0.00705661 |
| 768530 | *PLA2G15* | 1.12214508 | 0.00327073 |
| 427365 | *PLAA* | 1.44579171 | 0.0000527 |
| 416730 | *PLCB4* | -0.9386808 | 0.0025857 |
| 418417 | *PLCXD2* | 1.00846249 | 0.00039326 |
| 418182 | *PLCZ1* | 3.72193241 | 0.00015098 |
| 423940 | *PLEKHA1* | -0.7737581 | 0.00116941 |
| 423069 | *PLEKHA7* | 0.99753819 | 0.00317695 |

| | | | |
|---|---|---|---|
| 101750407 | *PLEKHS1* | -2.7549839 | 0.00017324 |
| 424586 | *PLK3* | 1.1376163 | 0.00865236 |
| 423980 | *PMS1* | 0.84184016 | 0.00025194 |
| 769547 | *PNO1* | 0.97100645 | 0.00119916 |
| 418233 | *PNPLA3* | -0.9080586 | 0.00048821 |
| 427210 | *POC5* | 0.93679315 | 0.00275124 |
| 418326 | *POLQ* | 1.13644591 | 0.00738208 |
| 426805 | *POLR1E* | 0.71322687 | 0.00606763 |
| 428240 | *POU2F3* | 1.83962265 | 0.00105299 |
| 395521 | *POU4F3* | 2.7498375 | 0.00079769 |
| 421764 | *PPIL6* | -1.7936524 | 0.00134959 |
| 415941 | *PPM1M* | -1.1380183 | 0.00204206 |
| 421287 | *PPP1R21* | 0.9367322 | 0.00049734 |
| 422858 | *PPP2R2C* | 0.4643394 | 0.00625914 |
| 416098 | *PPP4R2* | 0.6003035 | 0.00930459 |
| 423848 | *PPRC1* | 1.03803302 | 0.00133506 |
| 425977 | *PRCC* | 0.71740032 | 0.00364014 |
| 419478 | *PRDM2* | 0.7138372 | 0.00070767 |
| 427017 | *PRKAB2* | 0.86044082 | 0.00096643 |
| 419399 | *PRKCZ* | 0.74029831 | 0.00310208 |
| 428122 | *PRKRIR* | 0.77194577 | 0.0019445 |
| 428985 | *PRLHR* | 1.40153121 | 0.00615541 |
| 422975 | *PRMT3* | 0.93807827 | 0.00137008 |
| 418639 | *PRPS2* | 1.29214954 | 0.00281102 |
| 416957 | *PRR14L* | 0.65008179 | 0.00440456 |
| 426937 | *PSMA5* | 0.75922252 | 0.00271251 |
| 395806 | *PTCH1* | -0.5987313 | 0.00956256 |
| 396451 | *PTGS2* | -1.1439816 | 0.00168399 |
| 417725 | *PTPN12* | 0.71998521 | 0.00124629 |
| 421049 | *PTPRM* | 0.90960443 | 0.00727196 |
| 420449 | *PTPRN2* | 1.16179332 | 0.00010143 |
| 419554 | *PUM1* | -0.4195145 | 0.00972207 |
| 430546 | *PURA* | -0.5507049 | 0.00757939 |
| 421879 | *RAB23* | 0.88431266 | 0.00011019 |
| 416953 | *RAB36* | -1.0716511 | 0.00494726 |
| 421170 | *RABIF* | 0.97687508 | 0.00225568 |
| 426865 | *RABL2B* | -0.6365613 | 0.00743416 |
| 431623 | *RAD23B* | 0.74988645 | 0.00036469 |
| 420203 | *RALYL* | 1.27751581 | 0.00032788 |
| 418057 | *RASD2* | 1.66850151 | 0.00027273 |
| 416645 | *RBFOX1* | 1.30584028 | 0.00026788 |
| 417029 | *RBM19* | 1.26549069 | 0.0025701 |
| 107049003 | *RBM44* | 1.2003727 | 0.00050829 |
| 422404 | *RBM46* | 1.04539729 | 0.00813473 |

| | | | |
|---|---|---|---|
| 396449 | *RBP* | 5.84769348 | 2.33E-08 |
| 395678 | *RBPMS2* | 2.16881036 | 0.0020535 |
| 421380 | *RCOR3* | -1.0622294 | 0.00234781 |
| 421678 | *REPS1* | 1.34263062 | 0.0000492 |
| 419435 | *RERE* | 0.73747508 | 0.00456046 |
| 418180 | *RERGL* | 1.11829504 | 0.00293511 |
| 426549 | *RETSAT* | -1.0338512 | 0.00614293 |
| 422788 | *RFC1* | 0.78109313 | 0.00138653 |
| 418070 | *RFX4* | 1.41376239 | 0.00366436 |
| 418713 | *RFX8* | 1.47633618 | 0.00143197 |
| 101751119 | *RGS22* | -1.9421146 | 0.00054172 |
| 424800 | *RHBDD1* | 0.8211373 | 0.0033604 |
| 423059 | *RIC3* | 0.8454807 | 0.00698837 |
| 428142 | *RIMS4* | 0.94068383 | 0.0063964 |
| 421465 | *RMDN2* | 0.72828036 | 0.00930459 |
| 374021 | *RNF111* | 0.54199455 | 0.00598425 |
| 420085 | *RNF126* | 0.86516434 | 0.00662778 |
| 422382 | *RNF128* | 0.73748734 | 0.00854962 |
| 101751836 | *RNF186* | -3.1173838 | 0.00214391 |
| 421435 | *RNF8* | 0.99898557 | 0.00000263 |
| 421123 | *RP1* | -2.247506 | 0.00147758 |
| 418037 | *RP1-37E16.12* | 0.85292078 | 0.00856431 |
| 415326 | *RP11-152F13.10* | 1.0427308 | 0.0000937 |
| 771731 | *RP11-166N6.1* | -2.5354296 | 0.00181814 |
| 420326 | *RP11-240B13.2* | 1.5510073 | 0.00579273 |
| 420185 | *RP11-463D19.2* | 0.56427703 | 0.00552349 |
| 415812 | *RP11-505K9.4* | 0.76065965 | 0.00246711 |
| 395848 | *RP13-512J5.1* | 0.57526309 | 0.00477535 |
| 418675 | *RP2* | -0.8253609 | 0.00021531 |
| 768335 | *RP4-613B23.5* | 0.76248543 | 0.00618201 |
| 423795 | *RPP30* | 1.49031666 | 0.00048973 |
| 420253 | *RRM2B* | 0.61196098 | 0.00425282 |
| 421446 | *RSPH9* | -1.0898236 | 0.00321372 |
| 426519 | *RUFY1* | 0.77370119 | 0.00071244 |
| 421508 | *RYR2* | 0.88170042 | 0.0072473 |
| 426356 | *S100A9* | -2.1287043 | 0.00633379 |
| 395359 | *SALL3* | 1.50746687 | 0.00018894 |
| 769286 | *SALL4* | 1.30886986 | 0.0000454 |
| 100857189 | *SAMD3* | 1.0891354 | 0.00458661 |
| 419125 | *SAMHD1* | -0.748283 | 0.00584057 |
| 417375 | *SAP30BP* | 0.58225497 | 0.0081189 |
| 770088 | *SCFD2* | -0.7943018 | 0.00402741 |
| 422486 | *SCLT1* | 0.88792387 | 0.00032788 |
| 771555 | *SCN1A* | 0.77343808 | 0.00527637 |

| | | | |
|---|---|---|---|
| 395945 | *SCN2A* | 0.92177167 | 0.00742005 |
| 395947 | *SCN5A* | 1.41939933 | 0.00293511 |
| 396050 | *SCNN1A* | -1.3201048 | 0.00099197 |
| 772136 | *SDCCAG3* | 1.46929223 | 0.0000772 |
| 421318 | *SDE2* | 0.57104749 | 0.00243849 |
| 415806 | *SDR42E1* | 1.01793683 | 0.00933488 |
| 426850 | *SEC11C* | 1.02661129 | 0.0037328 |
| 415945 | *SEMA3G* | -1.5871364 | 0.00079928 |
| 107057370 | *SEMA4C* | -1.5665931 | 0.00621623 |
| 396228 | *SERPINH1* | -0.8248061 | 0.00293111 |
| 423445 | *SETD3* | 0.63779797 | 0.00472483 |
| 101750922 | *SETD6* | 1.14716208 | 0.0049692 |
| 427717 | *SF3A1* | 0.96407861 | 0.00019126 |
| 420077 | *SF3A2* | 0.90938607 | 0.00081363 |
| 420732 | *SFRP4* | -1.7841235 | 0.00760581 |
| 107049626 | *SFTPC* | 3.7232591 | 0.00061536 |
| 422739 | *SGCZ* | 1.16569436 | 0.00416054 |
| 776038 | *SGIP1* | 2.0543453 | 0.00017329 |
| 378907 | *SGMS1* | -0.7309501 | 0.00126778 |
| 424070 | *SGOL2* | 0.98343328 | 0.00302817 |
| 426396 | *SGTA* | 0.58469932 | 0.00138653 |
| 769640 | *SH2D4B* | 1.52236489 | 0.00778806 |
| 421851 | *SH3BGRL2* | -1.7417586 | 0.00026241 |
| 770341 | *SHC2* | -1.1582403 | 0.00033445 |
| 426482 | *SHC4* | 1.13654812 | 0.00548938 |
| 395843 | *SIX6* | -3.0351389 | 0.0000394 |
| 416808 | *SLC15A4* | -0.7453143 | 0.00756791 |
| 395383 | *SLC16A3* | -1.1236321 | 0.00011311 |
| 423921 | *SLC18A2* | 1.27576829 | 0.00814087 |
| 427352 | *SLC1A1* | -1.1422423 | 0.00091201 |
| 421582 | *SLC22A3* | 1.5066094 | 0.0030967 |
| 417700 | *SLC26A3* | -2.5519815 | 0.0000521 |
| 427845 | *SLC26A4* | -2.019142 | 0.00091092 |
| 427459 | *SLC28A3* | 2.11825751 | 0.0041984 |
| 417807 | *SLC38A2* | -1.0383853 | 0.00163213 |
| 420308 | *SLC45A4* | -0.8178574 | 0.00248752 |
| 421420 | *SLC4A1AP* | 0.78917991 | 0.00165791 |
| 424916 | *SLC51A* | -3.1326498 | 0.0000287 |
| 419805 | *SLC6A17* | 0.89843071 | 0.0041984 |
| 415768 | *SLC7A9* | -3.7081986 | 0.0000102 |
| 418719 | *SLC9A2* | -2.4099225 | 2.48E-05 |
| 100859280 | *SLITRK5* | 1.03762162 | 0.00644019 |
| 419641 | *SMAP2* | 0.75015679 | 0.00400354 |
| 422522 | *SMARCAD1* | 0.59615914 | 0.00510179 |

| | | | |
|---|---|---|---|
| 396156 | *SMC2* | 0.75349306 | 0.00833575 |
| 101749387 | *SMIM24* | -3.9533048 | 0.000089 |
| 423889 | *SMNDC1* | 0.66058567 | 0.00206142 |
| 396444 | *SNAP25* | 1.23283462 | 0.00020131 |
| 427480 | *SNX1* | 1.00579065 | 0.00177021 |
| 420202 | *SNX16* | 0.87120859 | 0.00173427 |
| 416417 | *SNX29* | -0.4699176 | 0.00552018 |
| 421654 | *SNX9* | 0.77410079 | 0.00408648 |
| 421775 | *SOBP* | 0.72935562 | 0.00977608 |
| 421713 | *SOGA3* | 3.45882907 | 0.00147758 |
| 423885 | *SORCS1* | 0.99040571 | 0.00013631 |
| 415332 | *SORD* | -0.9314295 | 0.00510179 |
| 395526 | *SOX14* | 2.18262487 | 0.001706 |
| 428534 | *SOX17* | -1.7606961 | 0.00863482 |
| 416243 | *SOX30* | 1.93557907 | 0.00026788 |
| 395143 | *SOX4* | -1.5976175 | 0.00642402 |
| 416172 | *SPDL1* | 1.05910994 | 0.00010761 |
| 417595 | *SPECC1* | 1.00930689 | 0.00063552 |
| 415572 | *SPG11* | 0.59731327 | 0.0031356 |
| 424789 | *SPHKAP* | 0.98826916 | 0.00929878 |
| 395879 | *SPI1* | -1.2041925 | 0.00760696 |
| 416235 | *SPINK5* | -2.6962248 | 0.00130353 |
| 415855 | *SPIRE2* | -1.7474101 | 0.00011311 |
| 768915 | *SPO11* | 1.95154271 | 0.00025849 |
| 423225 | *SPTBN5* | -2.1232561 | 0.00018272 |
| 425008 | *SPTSSB* | -2.1789016 | 0.00053831 |
| 420335 | *SQLE* | -0.696554 | 0.00177021 |
| 426281 | *SRCIN1* | 1.04078198 | 0.00737173 |
| 420538 | *SRI* | -1.7608787 | 0.00097839 |
| 419731 | *ST14* | -0.7692361 | 0.00839752 |
| 395138 | *ST3GAL6* | 0.68381135 | 0.00293511 |
| 404746 | *ST6GALNAC3* | 0.902027 | 0.00284107 |
| 395331 | *ST7L* | 2.48319486 | 0.00705661 |
| 414796 | *ST8SIA3* | 2.81287407 | 0.00644183 |
| 419213 | *STAU1* | 0.77780839 | 0.00113314 |
| 420184 | *STAU2* | 1.06089116 | 0.00039543 |
| 378801 | *STK25* | 0.75505151 | 0.00039104 |
| 420621 | *STK31* | 1.09782793 | 0.00525031 |
| 422850 | *STK32B* | 1.67631689 | 0.0000228 |
| 101751124 | *STK39* | 0.67621315 | 0.00876847 |
| 417118 | *STOM* | -0.7535142 | 0.00550702 |
| 423587 | *STYX* | 1.25520645 | 0.00091092 |
| 427508 | *SV2B* | 1.16049838 | 0.00534907 |
| 100859817 | *SWT1* | 0.57412474 | 0.00697824 |

| | | | |
|---|---|---|---|
| 100859541 | *SYCE2* | 2.64423711 | 0.00081363 |
| 396359 | *SYNPR* | 0.91094082 | 0.00161584 |
| 426394 | *SYTL2* | 0.98576936 | 0.00022798 |
| 428796 | *TADA2B* | 0.88389156 | 0.00185168 |
| 416784 | *TANGO2* | 0.66549899 | 0.00235458 |
| 426162 | *TBC1D1* | 0.47853459 | 0.00842272 |
| 421181 | *TBC1D22B* | 0.71761815 | 0.00815639 |
| 429029 | *TBR1* | 1.86547681 | 0.00126864 |
| 373943 | *TBX20* | 2.35174615 | 0.00052257 |
| 373988 | *TBX5* | 1.55992846 | 0.00929878 |
| 395782 | *TBXT* | 1.45755509 | 0.00769148 |
| 416276 | *TCOF1* | 0.86564408 | 0.00111343 |
| 421567 | *TCP10* | -2.3330414 | 0.00210399 |
| 100857412 | *TCTN3* | -0.5038722 | 0.00246711 |
| 423403 | *TDP1* | 0.94141405 | 0.00246711 |
| 428669 | *TDRD6* | 1.12774913 | 0.00595551 |
| 423488 | *TDRD9* | 1.16157826 | 0.00510179 |
| 403089 | *TEAD1* | -0.5689945 | 0.00087316 |
| 395668 | *TENM1* | 0.94358553 | 0.00342451 |
| 419327 | *TFAP2C* | 1.9975647 | 0.00086763 |
| 431620 | *THAP1* | 1.26306811 | 0.00033445 |
| 771761 | *THL* | 2.8857503 | 0.0081189 |
| 378897 | *THY1* | 2.11135104 | 0.00078689 |
| 415489 | *TICRR* | 0.91900079 | 0.0034418 |
| 416250 | *TIMD4* | -1.9472942 | 0.00393445 |
| 420070 | *TJP3* | -1.4228891 | 0.00048821 |
| 775978 | *TM2D1* | 0.72953741 | 0.00063545 |
| 415462 | *TM6SF1* | -0.8679316 | 0.0022047 |
| 420985 | *TMEFF1* | 1.15847595 | 0.0012876 |
| 768542 | *TMEFF2* | 1.17815764 | 0.00070041 |
| 416809 | *TMEM132C* | 1.19238397 | 0.00028492 |
| 421284 | *TMEM17* | 0.98551728 | 0.00329906 |
| 107053202 | *TMEM192* | 0.78160719 | 0.00048183 |
| 772254 | *TMEM196* | -1.6667232 | 0.0037643 |
| 419725 | *TMEM45B* | -2.4260088 | 0.0000293 |
| 419590 | *TMEM57* | 0.86283544 | 0.00714085 |
| 427967 | *TMPRSS15* | 1.88652733 | 0.00119916 |
| 418528 | *TMPRSS2* | -1.937098 | 0.0001512 |
| 378894 | *TNFSF10* | -1.0273937 | 0.00415282 |
| 421138 | *TOX* | -1.0548577 | 0.00930459 |
| 428145 | *TP53RK* | 0.88419888 | 0.00728738 |
| 374269 | *TP63* | 1.77260846 | 0.00020759 |
| 418885 | *TPTE2* | 0.8158726 | 0.00254999 |
| 395403 | *TRA2B* | -0.9312832 | 0.00520395 |

| | | | |
|---|---|---|---|
| 416884 | *TRAFD1* | -0.6408915 | 0.00803636 |
| 421095 | *TRAPPC8* | 0.56645871 | 0.0022047 |
| 378919 | *TRIB2* | -0.7414554 | 0.0040723 |
| 418872 | *TRIM13* | -0.9210366 | 0.0000889 |
| 419883 | *TRIM33* | 0.7836185 | 0.00218874 |
| 417628 | *TRIM37* | 0.63082038 | 0.00468683 |
| 424818 | *TRIM42* | 2.2689097 | 0.00048973 |
| 424785 | *TRIP12* | 0.50934593 | 0.00697824 |
| 420798 | *TRIP13* | 1.10146205 | 0.00076806 |
| 426860 | *TRMT1L* | 0.83760208 | 0.00020175 |
| 768091 | *TSC22D3* | -1.17244 | 0.000395 |
| 769262 | *TSNAXIP1* | -1.3050715 | 0.00204206 |
| 417854 | *TSPAN8* | -3.2648141 | 9.96E-07 |
| 421916 | *TSSC1* | 0.69118407 | 0.00126778 |
| 770784 | *TTC12* | -0.7105585 | 0.00140342 |
| 423407 | *TTC7B* | 0.69952887 | 0.00933488 |
| 421849 | *TTK* | 0.85940996 | 0.00759336 |
| 423061 | *TUB* | -0.5776514 | 0.00827574 |
| 415590 | *TUBGCP4* | 0.49934034 | 0.00811282 |
| 426085 | *TULP3* | 0.60216735 | 0.00179709 |
| 420603 | *TWISTNB* | 0.90616235 | 0.00943755 |
| 107051021 | *TXNIP* | -1.184333 | 0.0000889 |
| 419748 | *UBASH3B* | -0.5390149 | 0.00767653 |
| 416678 | *UBE2H* | -0.4500183 | 0.00393445 |
| 771251 | *UBFD1* | 2.13746083 | 5.62E-07 |
| 418921 | *UBL3* | -0.6534936 | 0.00833575 |
| 416398 | *UBN1* | 0.79127584 | 0.00167716 |
| 417614 | *ULK2* | -0.5984903 | 0.00965537 |
| 395101 | *UNC5C* | 0.95810211 | 0.00272457 |
| 416458 | *UNCX* | 1.56942666 | 0.00044091 |
| 424553 | *USP33* | 0.92014702 | 0.00061542 |
| 769457 | *USP48* | 0.92619427 | 0.00095758 |
| 421364 | *VASH2* | 1.0495412 | 0.00075895 |
| 430410 | *VAT1L* | 0.99638012 | 0.00392186 |
| 396423 | *VIL1* | -5.0292306 | 9.08E-09 |
| 421702 | *VNN1* | -1.66147 | 0.00308 |
| 415750 | *VPS35* | 0.52612057 | 0.00762862 |
| 423350 | *VRTN* | 1.35161684 | 0.00226833 |
| 395537 | *VSX1* | 2.04352695 | 0.00111456 |
| 424533 | *VTG2* | 1.68600137 | 0.0000399 |
| 416616 | *VWA3A* | -1.1819713 | 0.00091092 |
| 421761 | *WASF1* | 1.41199443 | 0.00057849 |
| 418145 | *WASH1* | 0.78491064 | 0.00687565 |
| 424806 | *WDFY1* | 0.69484096 | 0.00245568 |

| | | | |
|---|---|---|---|
| 423558 | *WDHD1* | 0.93679018 | 0.00727196 |
| 424344 | *WDR47* | 0.54906274 | 0.00281199 |
| 420427 | *WDR48* | 0.74192961 | 0.0063964 |
| 427444 | *WDR70* | 1.10171668 | 0.00036584 |
| 423985 | *WDR75* | 0.50504837 | 0.00636833 |
| 429114 | *WDR78* | -1.0985971 | 0.00757939 |
| 419300 | *WFDC2* | -5.4764285 | 2.48E-08 |
| 417831 | *WIF1* | -1.7642591 | 0.00439941 |
| 415984 | *WNK2* | 0.83734795 | 0.00124629 |
| 777580 | *WNK4* | 1.5891487 | 0.00150054 |
| 100858171 | *XKR4* | 1.35633754 | 0.00967653 |
| 428363 | *XKR9* | 2.28803745 | 0.000571 |
| 414780 | *XYLT2* | 1.9189847 | 0.0043001 |
| 428916 | *ZBTB1* | -0.5277814 | 0.00039197 |
| 422847 | *ZBTB49* | 1.6714756 | 0.0000897 |
| 107055117 | *ZBTB7B* | -1.4226397 | 0.00758249 |
| 419663 | *ZBTB8OS* | 0.89008239 | 0.0028496 |
| 428890 | *ZC2HC1C* | 1.37777273 | 0.00078689 |
| 415838 | *ZC3H18* | 0.78251494 | 0.00625914 |
| 417791 | *ZCRB1* | 1.09350436 | 0.00222538 |
| 419342 | *ZFP64* | 1.10544525 | 0.00014942 |
| 422889 | *ZFYVE28* | 0.88815443 | 0.00249406 |
| 424639 | *ZFYVE9* | -0.7677769 | 0.00017434 |
| 425428 | *ZMAT3* | 0.56990413 | 0.00909905 |
| 101749232 | *ZNF281* | 0.68026241 | 0.00668242 |
| 416144 | *ZNF346* | 0.60657397 | 0.00281102 |
| 421881 | *ZNF451* | 1.00963881 | 0.00034944 |
| 769868 | *ZNF512* | 0.70907371 | 0.00359314 |
| 422841 | *ZNF518B* | 0.71339511 | 0.00157165 |
| 373921 | *ZNF622* | 0.58301304 | 0.00910575 |
| 770560 | *ZNF703* | 1.69360112 | 0.00458661 |
| 423295 | *ZNF770* | 0.85541814 | 0.00218249 |
| 423994 | *ZNF804A* | 1.32367031 | 0.00026114 |
| 422465 | *ZNF827* | 0.75867775 | 0.00283738 |
| 420152 | *ZNRF4* | 4.28072713 | 0.0000337 |
| 404532 | *ZPBP2* | 1.85278462 | 0.00323173 |
| 418405 | *ZPLD1* | 3.09182008 | 0.000089 |
| 423953 | *ZRANB1* | 0.96422327 | 0.0001181 |
| 424291 | *ZRANB3* | 0.98096382 | 0.00527637 |
| 424645 | *ZYG11B* | 0.57790662 | 0.00904711 |

**Table 6:** Sex-specific, restraint stress responsive genes that were differentially expressed in the male gonads.

| Entrez ID | Gene Name | logFC | FDR |
|---|---|---|---|
| 101748426 | *LOC101748426* | -2.1186961 | 0.00614228 |
| 101749540 | *LOC101749540* | 4.03908776 | 0.00561745 |
| 422387 | *MTMR1* | 1.17189367 | 0.00478273 |
| 395878 | *PAPOLA* | 0.65626868 | 0.00967625 |
| 423433 | *SERPINA4* | -3.6531707 | 0.00478273 |
| 423414 | *TRIP11* | -1.4596995 | 0.00967625 |

**Table 7:** Differentially expressed genes found in the hypothalamus of both sexes for restraint stress.

| Entrez ID | Gene Name | Male logFC | Male FDR | Female logFC | Female FDR |
|---|---|---|---|---|---|
| 423548 | *AP5M1* | 2.663776185 | 0.002159163 | 2.942159249 | 0.000106574 |
| 378911 | *GHSR* | 3.820809349 | 0.004702544 | 4.025969752 | 0.000703197 |
| 420716 | *LOC420716* | 4.679893006 | 0.0000435 | 4.40610442 | 0.001241845 |
| 417158 | *RPL7A* | 1.573845101 | 0.008847658 | 1.645257722 | 0.001405972 |

**Table 8:** Differentially expressed genes found in the pituitary of both sexes for restraint stress.

| Entrez ID | Gene Name | Male logFC | Male FDR | Female logFC | Female FDR |
|---|---|---|---|---|---|
| 101750033 | *ANGPTL7* | 2.059675268 | 0.005145346 | 2.623379629 | 0.0000594 |
| 769889 | *APOLD1* | -2.695563361 | 0.00000154 | -2.791016195 | 1.26E-06 |
| 421369 | *ATF3* | -3.086151839 | 4.06E-09 | -2.962941643 | 2.76E-08 |
| 395468 | *CCL4* | 2.424131523 | 0.008263933 | 3.263222669 | 0.000183615 |
| 107052707 | *CEBPD* | -2.567830821 | 3.10E-09 | -1.885295619 | 1.73E-05 |
| 395335 | *CISH* | -1.764782855 | 3.16E-10 | -1.807300415 | 1.91E-10 |
| 378903 | *CREM* | -0.76528046 | 0.003887469 | -0.850364519 | 0.000648189 |
| 420425 | *CSRNP1* | -0.787358144 | 0.00050493 | -0.806016798 | 0.000277564 |
| 428137 | *EMILIN3* | 2.539871512 | 0.000247545 | 3.837617407 | 0.00000104 |
| 421416 | *FOSL2* | -2.110305949 | 0.00014188 | -2.944149432 | 0.000000405 |
| 425213 | *GRP* | -2.230872468 | 0.004796056 | -3.277770507 | 0.00000439 |
| 395128 | *HES4* | 1.69213783 | 1.18E-08 | 1.399863714 | 3.70E-06 |
| 428247 | *HTR3A* | -1.88753632 | 0.0000438 | -2.230419611 | 0.00000138 |
| 395925 | *KCNJ5* | -1.480040484 | 0.007270196 | -1.548871029 | 0.004088001 |
| 419829 | *KLHDC8A* | -0.888920223 | 0.000800229 | -0.850985647 | 0.001309962 |
| 373901 | *LMO4* | 2.148245199 | 0.000215575 | 1.673832449 | 0.00660783 |
| 100859084 | *LOC100859084* | 2.73843229 | 0.003120538 | 2.754098737 | 0.001811334 |
| 100859853 | *LOC100859853* | -2.537574391 | 3.1E-09 | -1.967896101 | 0.00000804 |
| 101751319 | *LOC101751319* | 5.085123808 | 0.0000137 | 4.71420299 | 0.0000896 |
| 107049603 | *LOC107049603* | -3.386594158 | 0.000051 | -4.149269713 | 0.00000524 |
| 107050337 | *LOC107050337* | -3.076897507 | 0.000426361 | -3.112363698 | 0.001026693 |
| 107050461 | *LOC107050461* | 3.960655909 | 0.000116488 | 2.887819778 | 0.004699478 |
| 107051321 | *LOC107051321* | 2.441903465 | 0.000426361 | 2.000660878 | 0.004120316 |
| 374126 | *NAB1* | -0.512982492 | 0.002566327 | -0.592217228 | 0.000206412 |
| 420996 | *NR4A3* | -2.852356984 | 5.25E-10 | -3.419441009 | 4.87E-12 |
| 396214 | *PLP1* | -3.786766846 | 0.001590204 | -10.6939925 | 4.37E-12 |
| 429116 | *PTGER3* | -1.816054283 | 0.000790523 | -1.507306835 | 0.007497314 |
| 100857976 | *S1PR2* | 3.287102232 | 0.0000267 | 2.174033231 | 0.006797303 |
| 378907 | *SGMS1* | -0.85600155 | 0.000325349 | -0.755694608 | 0.001824287 |
| 421035 | *SLMO1* | -0.813644712 | 0.00000506 | -1.096475088 | 5.98E-10 |
| 432368 | *SNAI2* | 1.330695786 | 0.004433902 | 1.373591676 | 0.001410526 |
| 419225 | *SS18L1* | 1.033425848 | 0.002811186 | 0.980008791 | 0.004255233 |
| 419443 | *TMEM201* | -0.919424013 | 0.001555148 | -1.537415124 | 1.73E-08 |
| 768091 | *TSC22D3* | -1.023907228 | 0.00657757 | -1.647421547 | 1.16E-06 |
| 419759 | *ZBTB16* | -2.158469442 | 2.95E-05 | -2.85962211 | 5.57E-08 |

**Table 9:** Differentially expressed genes found in the gonads of both sexes for restraint stress.

| Entrez ID | Gene Name | Male logFC | Male FDR | Female logFC | Female FDR |
|---|---|---|---|---|---|
| 428310 | *HSP25* | -2.5402025 | 0.00478273 | -2.9095266 | 4.96E-07 |
